# Contribution of the IGCR1 regulatory element and the 3*’Igh* CBEs to Regulation of *Igh* V(D)J Recombination

**DOI:** 10.1101/2023.04.21.537836

**Authors:** Zhuoyi Liang, Lijuan Zhao, Adam Yongxin Ye, Sherry G. Lin, Yiwen Zhang, Chunguang Guo, Hai-Qiang Dai, Zhaoqing Ba, Frederick W. Alt

## Abstract

Immunoglobulin heavy chain variable region exons are assembled in progenitor-B cells, from V_H_, D, and J_H_ gene segments located in separate clusters across the *Igh* locus. RAG endonuclease initiates V(D)J recombination from a J_H_-based recombination center (RC). Cohesin-mediated extrusion of upstream chromatin past RC-bound RAG presents Ds for joining to J_H_s to form a DJ_H_-RC. *Igh* has a provocative number and organization of CTCF-binding-elements (CBEs) that can impede loop extrusion. Thus, *Igh* has two divergently oriented CBEs (CBE1 and CBE2) in the IGCR1 element between the V_H_ and D/J_H_ domains, over 100 CBEs across the V_H_ domain convergent to CBE1, and 10 clustered 3’*Igh*-CBEs convergent to CBE2 and V_H_ CBEs. IGCR1 CBEs segregate D/J_H_ and V_H_ domains by impeding loop extrusion-mediated RAG-scanning. Down-regulation of WAPL, a cohesin unloader, in progenitor-B cells neutralizes CBEs, allowing DJ_H_-RC-bound RAG to scan the VH domain and perform VH-to-DJH rearrangements. To elucidate potential roles of IGCR1-based CBEs and 3’*Igh*-CBEs in regulating RAG-scanning and elucidate the mechanism of the “ordered” transition from D-to-J_H_ to V_H_-to-DJ_H_ recombination, we tested effects of deleting or inverting IGCR1 or 3’*Igh*-CBEs in mice and/or progenitor-B cell lines. These studies revealed that normal IGCR1 CBE orientation augments RAG-scanning impediment activity and suggest that 3’*Igh*-CBEs reinforce ability of the RC to function as a dynamic loop extrusion impediment to promote optimal RAG scanning activity. Finally, our findings indicate that ordered V(D)J recombination can be explained by a gradual WAPL down-regulation mechanism in progenitor B cells as opposed to a strict developmental switch.

**SIGNIFICANCE STATEMENT:** To counteract diverse pathogens, vertebrates evolved adaptive immunity to generate diverse antibody repertoires through a B lymphocyte-specific somatic gene rearrangement process termed V(D)J recombination. Tight regulation of the V(D)J recombination process is vital to generating antibody diversity and preventing off-target activities that can predispose the oncogenic translocations. Recent studies have demonstrated V(D)J rearrangement is driven by cohesin-mediated chromatin loop extrusion, a process that establishes genomic loop domains by extruding chromatin, predominantly, between convergently-oriented CTCF looping factor-binding elements (CBEs). By deleting and inverting CBEs within a critical antibody heavy chain gene locus developmental control region and a loop extrusion chromatin-anchor at the downstream end of this locus, we reveal how these elements developmentally contribute to generation of diverse antibody repertoires.

## INTRODUCTION

Variable region exons that encode antigen-binding sites of antibodies are assembled in progenitor(“pro”)-B cells from germline V_H_, D, and J_H_ gene segments (1). V(D)J recombination is initiated by RAG1/2 endonuclease (RAG) (2). RAG introduces DNA double-stranded breaks (DSBs) between V_H_, D, and J_H_ coding segments and flanking recombination signal sequences (RSSs) (2). RSSs comprise a conserved heptamer, a spacer of 12 or 23 base pairs, and an AT-rich nonamer. To robustly initiate V(D)J recombination, RAG must bind and cleave a pairs of gene segments flanked by RSSs with complementary 12 and 23 bp spacers (termed 12-RSSs and 23-RSSs respectively) (2). After RAG cleavage, 12/23-RSS matched gene segment ends and, separately, their RSS ends are fused by the classical nonhomologous end-joining (3). The mouse IgH locus (*Igh*) spans 2.7 megabases (Mbs) on chromosome 12 with over 100 V_H_s interspersed within a several Mb distal portion (1). This V_H_ domain lies 100 kb upstream of a 50 kb region containing up to 13 Ds, with 4 J_H_s embedded within a 2 kb region just downstream of the most proximal D (DQ52) (1). RAG initiates *Igh* V(D)J recombination from a recombination center (RC) formed within highly transcribed chromatin that spans DQ52, the four J_H_s, and the intronic enhancer (iE*μ*) (4, 5). The V_H_s and J_H_s have 23RSSs and cannot be directly joined. Ds are flanked on either side by 12RSSs, allowing them to join to a downstream J_H_ and an upstream V_H_ to form a V(D)J exon (3). V(D)J recombination is developmentally ordered with Ds joined to a J_H_ to form a DJ_H_ RC, after which V_H_s are joined to the upstream D12RSS of the DJ_H_ RC (3).

The CTCF chromatin looping factor binds target DNA sequences, termed CTCF binding elements (“CBEs or “CTCF sites”), in an orientation-specific manner (6, 7). In this regard, the numerous genomic CBEs, when in adjacent regions, can occur in the same, divergent, or in convergent orientations (8). The cohesin complex mediates extrusion of chromatin loops genome-wide, forming contact loops when extrusion in each direction reaches CTCF-bound CBEs that impede extrusion (9, 10). Such CBE-anchored chromatin loops occur most dominantly between CBEs in convergent orientation (8, 11), an orientation that forms the most stable anchor (9, 10). Convergent CBE orientation has been implicated in mediating physiological functions (12–17). However, CTCF-bound CBEs can impede loop extrusion regardless of orientation (18), and non-CBE-based impediments, for example highly transcribed chromatin, can impede extrusion and contribute to developmental or tissue-specific regulation of loop extrusion (18–24). The *Igh* contains numerous CBEs with remarkable relative orientation. The IGCR1 element in the V_H_ to D domain just upstream of the distal DFL16.1 has two divergently-oriented CBEs, namely upstream CBE1 and downstream CBE2 (25). The V_H_ domain has over 100 CBEs with most lying in convergent orientation to IGCR1 CBE1 (26). Ten consecutive CBEs, termed 3’*Igh* CBEs (27, 28) lie at the just downstream of *Igh* in convergent orientation to IGCR1 CBE2 and V_H_ domain CBEs (25, 26). This organization has been proposed to have various functions in regulating V(D)J recombination (25, 29–31).

Cohesin-mediated loop extrusion provides the mechanistic underpinnings for RC-bound RAG to scan chromatin across the *Igh* locus for substrates (18, 19, 22, 24, 32). In pro-B cells, RC-bound RAG initiates scanning upon binding a J_H_-23RSS into one of its two active sites (1, 18, 24). For this scanning process, the active RC, which lacks CBEs, serves as a transcription-based dynamic downstream loop extrusion anchor, while IGCR1 serves as a CBE-based upstream anchor that terminates RAG scanning, preventing scanning from entering the V_H_ domain (18, 24, 33). The upstream orientation of the RAG-bound J_H_ programs RAG scanning of upstream D-containing chromatin extruded past the RC (24). During RAG scanning of the D locus, only downstream D-12RSSs, convergently oriented to the J_H_23RSSs, are used, resulting in D-to-J_H_ joining that deletes all sequences between the participating D and J_H_ (24). Predominant utilization of RSSs, or cryptic RSSs, in convergent orientation to the initiating RC RSS is a mechanistic property of the linear RAG scanning process (22, 32).

While V_H_s-23RSSs are compatible for joining to D12-RSSs, the IGCR1 impedes access of V_H_s to the D locus during the D-to-J_H_ rearrangement and, thereby, enforces ordered D-to-J_H_ before V_H_-to-DJ_H_ V(D)J recombination in pro-B cells (18, 25, 32). Mutational inactivation of IGCR1 CBEs allows RC-bound RAG to scan directly into the proximal V_H_ locus, causing the most proximal functional V_H_5-2 to robustly rearrange and dominate the V_H_ repertoire, with little rearrangement of more distal V_H_s (18, 25, 32). The mechanism by which V_H_5-2 dominates rearrangement when IGCR1 CBEs are inactivated is based on its RSS-associated CBE that impedes extrusion past the RC, making it accessible for rearrangement (18). Indeed, dozens of the proximal V_H_s have RSS-associated CBEs. Thus, while deletion of the V_H_5-2 CBE results in a 50-fold reduction of V_H_5-2 rearrangement along with greatly reduced RC interaction, the next upstream V_H_ becomes dominantly rearranged based on its RSS-associated CBE (18). Because dozens of the D-proximal V_H_s have RSS-associated CBEs, this portion of the V_H_ locus is a major barrier to RAG scanning to further upstream V_H_s when IGCR1 CBEs are inactivated (18).

Fluorescent *in situ* hybridization and chromosome conformation capture (3C)-based studies revealed the V_H_ domain to undergo large-scale contraction in pro-B cells (34–42). V_H_ locus contraction was proposed to bring distal V_H_s into proximity with the DJ_H_ RC for recombinational access (43). Recent studies revealed that locus contraction is mediated by loop extrusion, which extends across the several Mb V_H_ locus due to approximately 4-fold down-regulation of the WAPL cohesin unloading factor in pro-B cells (44). In this context, depletion of WAPL in non-lymphoid cells extends genome-wide loop extrusion by increasing cohesin density, allowing it to bypass CBEs and potentially other impediments (45–47). WAPL down-regulation in pro-B cells down-regulates impediment activity of IGCR1 CBEs, proximal V_H_-associated CBEs, and other impediments, allowing RAG scanning from a DJ_H_ RC to extend linearly across the V_H_ domain (22). Abelson murine leukemia virus-transformed pro-B cell lines (“*v-Abl* lines”) can be viably arrested in the G1-cell cycle phase in which V(D)J recombination occurs (48). Introduction of RAG into G1-arrested RAG-deficient *v-Abl* lines activates robust RAG scanning across the D domain (19, 22, 24). However, scanning is impeded at IGCR1 and there is little V_H_-to-DJ_H_ joining (19, 22, 24). Inactivation of IGCR1 CBEs leads to dominant rearrangement of V_H_5-2 in *v-Abl* lines due to its RSS-associated CBE (18). In this regard, *v-Abl* lines have high WAPL levels and do not neutralize V_H_ locus CBEs (19, 22). Neutralization of CBE impediments by depletion of CTCF or WAPL extends cohesin loop extrusion and RAG-scanning past IGCR1 and proximal V_H_s to the most distal V_H_s (19, 22). Thus, *v-Abl* pro-B cell studies have provided substantial mechanistic insights into locus contraction and RAG (1).

Despite recent advances in elucidating functions of *Igh* CBEs during V(D)J recombination, many questions remain. While an early study indicated that IGCR1 CBEs act synergistically to segregate the V_H_ domain from the D/J_H_ domain (49), the question of whether orientation of these CBEs is critical to their function remained. Likewise, 3’*Igh* CBEs confine *Igh* class switch recombination (CSR) activity to the *Igh* (50). However, their roles in V(D)J recombination have not been resolved (1). Finally, a long-standing question is how the developmental transition from D-to-J_H_ joining to V_H_-to-DJ_H_ joining is regulated. While a role for the DJ_H_ intermediate in signaling the transition (51, 52) has been considered, such a transition could, in theory, be mediated by gradual WAPL-down-regulation during pro-B cell development (1). We now describe studies that address these questions.

## RESULTS

### Role of IGCR1 CBEs in D-to-J_H_ and Proximal V_H_-to-DJ_H_ rearrangements

To gain insight into factors that determine primary BM pro-B cell V_H_ repertoires, we assessed effects of IGCR1 CBE1 and CBE2 inactivation via high resolution HTGTS-V(D)J-Seq (24). For these assays, we purified B220^+^CD43^high^IgM^−^ pro-B cells from bone marrow (BM) of wild-type (WT) 129SV controls and previously generated IGCR1/CBE1&2^−/−^ mice (25). We performed HTGTS-V(D)J-Seq on DNA from these samples using a J_H_4 bait primer to compare their levels of D-to-J_H_ and V_H_-to-DJ_H_ rearrangements (Fig.1A and SI Appendix, Table S1) (32). A J_H_4 bait primer was used for HTGTS-V(D)J-Seq analyses in this study to eliminate potential confounding effects of rearrangements within extra-chromosomal deletion products (24). For these analyses, individual peaks found in HTGTS libraries can be normalized as a fraction of total HTGTS reads for a given experiment, which reveals absolute V(D)J levels of each rearranging gene segment. Alternatively, peaks can be normalized as a fraction of total recovered junctions across a given locus or section of a locus to reveal the relative utilization of a given gene segments to each other as a percentage of all junctions in the analyzed region (SI Appendix, Fig. S1). Such analyses are useful for examining effects of potential regulatory element mutations. For example, finding a decrease in total reads between two samples in the absence of differences in the junction profile, reflects decreased RAG or RC activity without changes in long range scanning patterns (22).

**Figure 1.**
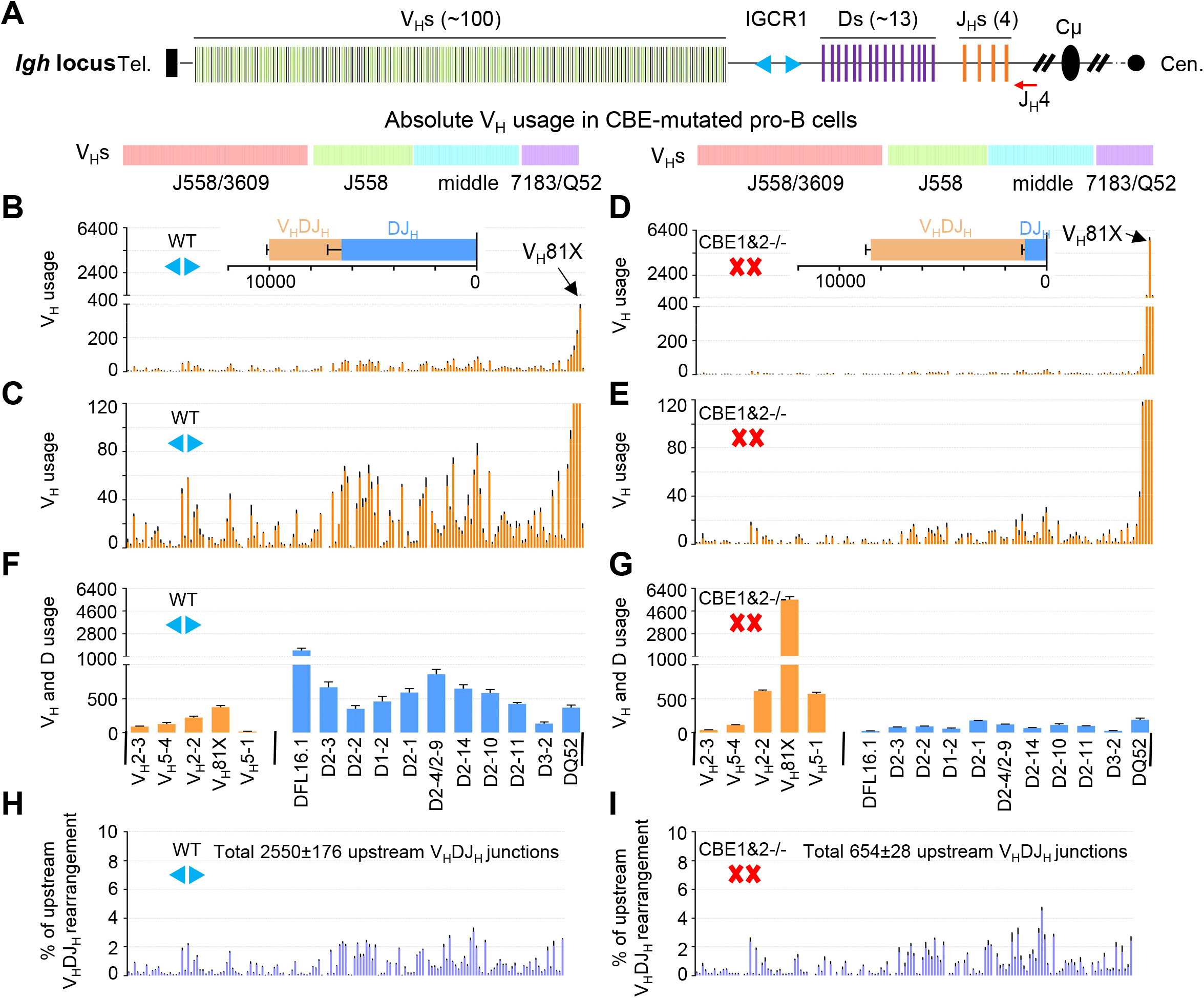
Role of IGCR1/CBE1 and CBE2 in D-to-J_H_ and proximal V_H_-to-DJ_H_ rearrangements. (A) Schematic of the murine *Igh* locus showing V_H_s, Ds, J_H_s, C_H_s and IGCR1. The red arrow indicates the J_H_4 coding end (CE) bait primer. (B-E) Utilization of V_H_s across the entire *Igh* locus in WT (B-C) and CBE1&2^−/−^ (D-E) pro-B cells. The V_H_DJ_H_ and DJ_H_ junctions are shown in the insets. (*n*=3 mice, mean±SEM; all HTGTS libraries are normalized to 78,091 total reads; see SI Appendix, Table S1). (F-G) Proximal V_H_ usage in V_H_DJ_H_ junctions and D usage DJ_H_ junctions of WT (F) and CBE1&2^−/−^ (G) pro-B cells (*n*=3 mice, mean±SEM). (H-I) Each panel shows the relative percentage of upstream V_H_s beyond the five most proximal V_H_s normalized to the indicated V_H_DJ_H_ junction number. Upstream V_H_s junctions are extracted from the data of Fig. 1B, D (*n*=3 mice, mean±SEM).

WAPL down-regulation in mouse pro-B cells (22, 44) allows RAG to scan the entire V_H_ locus, which results in highly reproducible utilization of the different V_H_s (Fig. 1B,C). However, in WT pro-B cells, V_H_5-2 (“V_H_81X”) and three immediately upstream V_H_s with RSS-associated CBEs are much more highly utilized than any V_H_s further upstream (Fig. 1B, C). In such steady-state BM pro-B cell populations, DJ_H_ rearranged alleles are more prevalent than V_H_(D)J_H_ rearranged alleles (Fig. 1B, inset), likely reflecting steady-state distributions within pro-B cells entering the compartment and successively generating DJ_H_ and V_H_-to-DJ_H_ rearrangements before leaving the compartment. Inactivation of IGCR1 by mutational inactivation of CBE1 and CBE2 (18, 25, 32), allows V_H_5-2 to dominate the V_H_DJ_H_ repertoire (Fig.1D,E). Moreover, the great majority of V_H_5-2 rearrangements are non-productive (SI Appendix, Fig. S2 and Table S2), consistent with selection against productive V_H_5-2 rearrangements (53). This finding also confirms that dominant V_H_5-2 rearrangements in pro-B cells do not result from cellular selection (25). Rearrangement frequencies of more distal V_H_s are dramatically decreased upon IGCR1 inactivation (Compare Fig. 1 panels B and C with panels D and E; also see SI Appendix, Table S1). Strikingly, however, in CBE1&2^−/−^ pro-B cell populations as compared to WT pro-B cell populations, the absolute level of V_H_DJ_H_ rearrangements is greatly increased with the vast majority utilizing V_H_5-2, while the absolute level of DJ_H_ rearrangements and upstream V_H_ rearrangements is, correspondingly, decreased (Fig. 1 E-G and SI Appendix, Table S1).

The above findings support the notion that, in the absence of IGCR1 CBE activity in normal pro-B cells, RAG scanning continues between the DJ_H_ RC and V_H_5-2, which allows V_H_5-2 to dominate V_H_-to-DJ_H_ rearrangements due to its robust CBE-mediated interaction with the DJ_H_ RC (18). Moreover, the finding that the increased frequency of V_H_-to-DJ_H_ rearrangements in the steady state CBE1&2^−/−^ pro-B results almost totally from increased V_H_5-2 rearrangements indicates that these dominant rearrangements occur at the normal DJ_H_ scanning stage before sufficient WAPL down-regulation neutralizes proximal V_H_-RSS associated CBE impediments to allow upstream scanning. These high resolution HTGTS-V(D)J-seq studies also revealed another notable finding. Thus, despite the dramatically reduced levels of upstream V_H_ rearrangements in CBE1&2^−/−^ pro-B cells, the upstream V_H_s beyond the several most proximal V_H_s, relative to each other, have rearrangement junction patterns (i.e. relative levels compared to each other), that were nearly identical to those of WT pro-B cells (Fig.1, Compare panels H and I). Thus, V_H_81X, and to a lesser extent immediately upstream V_H_s, dominate initial rearrangements in the absence of IGCR1 CBE activity, and, in doing so, terminate most RAG upstream scanning. However, RAG scanning that does proceed beyond V_H_5-2 continues through the remainder of the V_H_ locus, with similar V_H_ usage patterns as those of WT cells. Overall, these findings indicate that, while most V_H_5-2 rearrangements in CBE1&2^−/−^ pro-B cells occur before WAPL down-regulation, a small fraction of CBE1&2^−/−^ pro-B that do not form V_H_5-2 rearrangements on one or both *Igh* alleles undergo upstream V_H_-to-DJ_H_ recombination events at normal frequencies when WAPL-down-regulation reaches appropriate levels.

### Role of IGCR1/CBE1 and CBE2 in regulating RAG scanning into the V_H_ locus

To further assess potential mechanisms by which IGCR1 CBE impediment activity is modulated to promote RAG scanning of the V_H_ domain, we applied the highly sensitive 3C-HTGTS chromatin interaction assay to explore interactions of the RC-based iE*μ* enhancer element with upstream and downstream *Igh* locus chromatin domains in WT *rag2*^*-/-*^ and IGCR1/CBE1&2^−/−^*rag2*^*-/-*^ cultured pro-B cells derived from the corresponding mouse lines.

RAG-deficient cells must be used for such assays to eliminate confounding effects of V(D)J recombination events on such interactions (18, 19, 22, 24). These studies revealed that the iEμ/RC interacts robustly with 15 highly focused regions across the 2.4 Mb V_H_ locus in WT *rag2*^*-/-*^ pro-B cells (Fig. 2A; SI Appendix, Fig. S3) (19, 22). Among the most robust of these iE*μ* /RC interacting peaks are peaks associated with robustly transcribed PAX5-activated intergenic repeat (PAIR) elements (38, 54, 55) in the J558, and J558/3609 V_H_-containing regions in the distal portion of the V_H_ locus (Fig. 2A and SI Appendix, Fig. S3; Peaks 1,4,6,8-10). RC interactions with transcribed PAIR element-associated sequences are considered a hallmark of loop extrusion-mediated V_H_ locus contraction in pro-B cells (38). In this regard, locus contraction results from an approximately 4-fold developmental down-regulation of WAPL in pro-B cells(44), which at least partially neutralizes IGCR1-CBEs, proximal V_H_ CBEs, and likely other V_H_ locus CBEs, and potentially transcription-associated impediments to RC-based RAG linear scanning (22, 44). Although V_H_5-2 and proximal V_H_s are the most dominantly utilized V_H_s in normal pro-B cells (Fig 1B), they show only low-level interactions with the RC in RAG2-deficient WT pro-B cells at steady-state (Fig. 2A, B), consistent with most CBE-based interactions being diminished by WAPL-downregulation in a large fraction of the pro-B cells (22). In this regard, some WT *rag2*^*-/-*^ pro-B cells retain interactions between the RC and IGCR1(Fig. 2B, upper), indicating that some cells in the population have not fully down-regulated WAPL and/or that IGCR1 impediment activity is not completely neutralized by physiological levels of WAPL down-regulation (Fig. 2B; upper).

**Figure 2.**
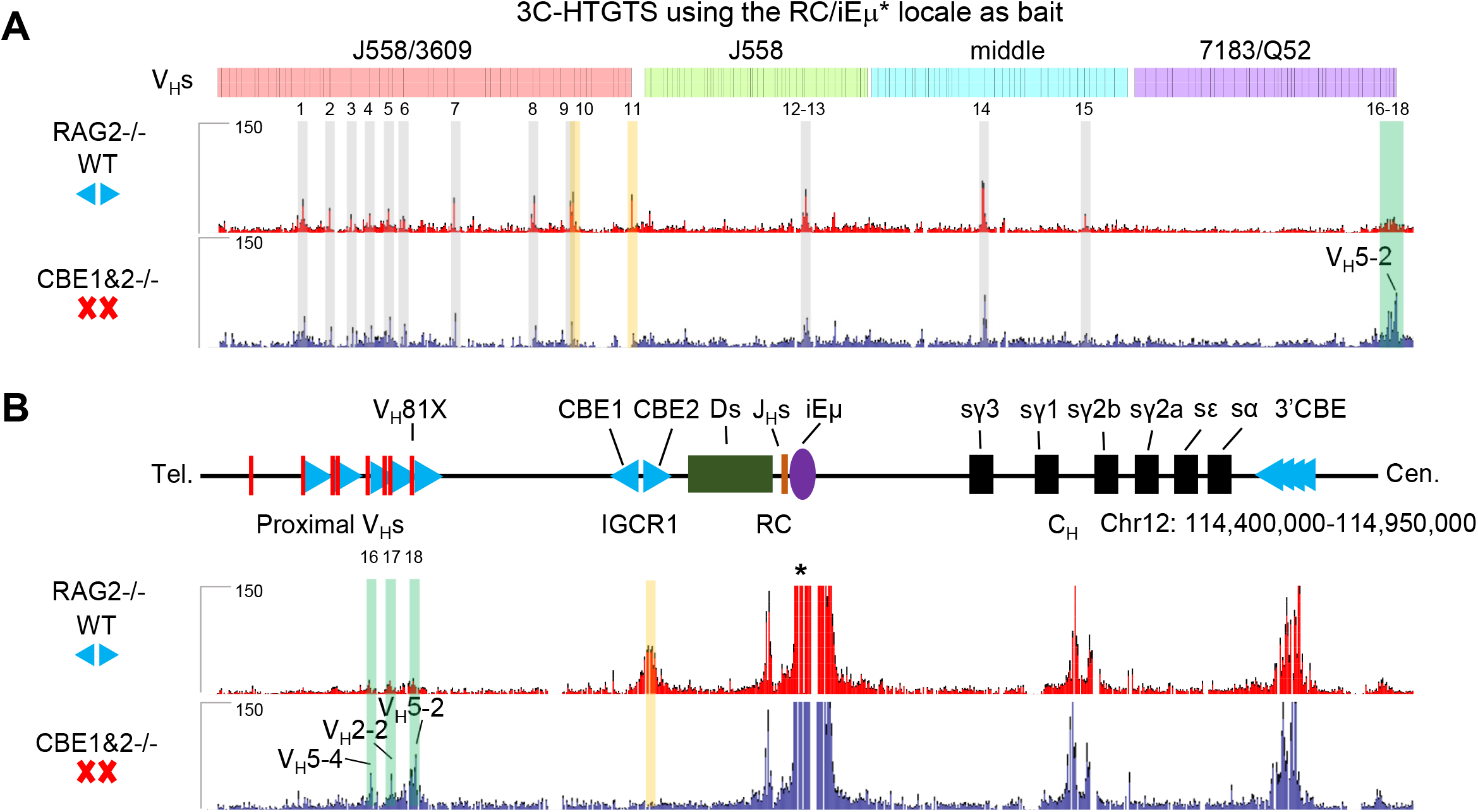
Role of IGCR1/CBE1 and CBE2 in regulating RAG scanning into the V_H_ locus. (A) 3C-HTGTS signal counts of all V_H_s in WT (red) and CBE1&2^−/−^ (blue) RAG2-deficient pro-B cells baiting from RC/iEμ (*). Each library was normalized to 160,314 total junctions (*n*=3 mice, mean±SEM). 18 peaks across V_H_s region are called by MACS2 pipeline and highlighted in gray (peaks 1-9, 12-15 are called in both conditions), orange (peaks 10-11 are called only in WT), green (peaks 16-18 are called only in CBE1&2^−/−^) (see SI Appendix, Fig. S3). (B) Zoom-in 3C-HTGTS profiles of *Igh* locus from proximal V_H_S to 3’*Igh* CBEs.

Notably, in CBE1&2^−/−^*rag2*^*-/-*^ cultured pro-B cells, the RC gained greatly increased interaction with proximal V_H_5-2 and the 3 proximal V_H_s immediately upstream, which, based on prior studies (18), is dependent on their RSS-associated CBEs (Fig. 2A,B, and SI Appendix, Fig. S3; Peaks 16-18). Strikingly, nearly all major further upstream interaction peaks were also present in chromatin from CBE1&2^−/−^*rag2*^*-/-*^ pro-B cells, mostly at similar relative levels to those in WT *rag2*^*-/-*^ pro-B cells (Fig. 2A. upper and lower). Given that the chromatin interactions are investigated in RAG-deficient cells and upstream interactions are not impacted by proximal V_H_- to DJ_H_ recombination events, it is not unexpected that large number of IGCR1/CBEs mutated pro-B cells would have robust interactions between the RC to the upstream V_H_s after developmental WAPL down-regulation. As previously described (18), the RC also robustly interacts downstream with the transcribed enhancer-like element between C*γ*1 and C*γ*2b (38) and with the 3’*Igh* CBEs in pro-B cells (Fig. 2B, upper); these interactions were not altered by IGCR1 CBE1 and CBE2 deletion (Fig. 2B, lower).

### Influence of IGCR1 CBEs and their orientation on V_H_-utilization in pro-B cells

We have previously generated CBE1^−/−^ and CBE2^−/−^ mice (49). To assess whether orientation of IGCR1/CBEs is critical for *Igh* V(D)J recombination control, we replaced CBE1 or CBE2 with their inverted sequences to generate an CBE1^inv^ or CBE2^inv^ alleles in mouse 129SV ES cells (SI Appendix, Fig. S4 A-B) (25, 49) (see Methods). We employed HTGTS-V(D)J-Seq to assay J_H_4-based utilization of the various V_H_s in pro-B populations harboring IGCR1 CBE deletion or inversion mutations (Fig. 3). CBE1^−/−^ pro-B cell populations have markedly increased V_H_5-2 rearrangements and markedly decreased rearrangements of more distal V_H_s, with the degree of increases and decreases modestly, but significantly, less than those observed for CBE1&2^−/−^ pro-B cells (Fig. 3A, E and SI Appendix, Table S1). In contrast, CBE2^−/−^ pro-B cells had very modestly increased V_H_5-2 rearrangements (2.5-fold; Fig. 3 B, E) and little impact on more distal V_H_ rearrangements (Fig. 3B, E and SI Appendix, Table S1). These findings indicate that CBE1 and CBE2 cooperatively provide the full impact of IGCR1 scanning impediment activities and unequivocally demonstrate that CBE1 plays a much more dominant role. CBE1^inv/inv^ BM pro-B cells also have significantly increased levels of proximal V_H_5-2 utilization relative to WT; but to a much lower extent than CBE1^−/−^ pro-B cells, indicating that ability of CBE1 to impede RAG scanning is dampened, but not abrogated, when inverted (Fig. 3C, E and SI Appendix, Table S1). In contrast, CBE2^inv/inv^ pro-B cells were very similar to WT pro-B cells with respect to utilization of V_H_5-2 and upstream V_H_s (Fig. 3D, E and SI Appendix, Table S1). Consistent with our findings for IGCR1/CBE1&2^−/−^ pro-B cells (Fig.1), the rearrangement pattern of upstream V_H_s, relative to each other, was not markedly impacted by CBE1 or CBE2 deletions or inversions (SI Appendix, Fig. S5 A-F and Table S1).

**Figure 3.**
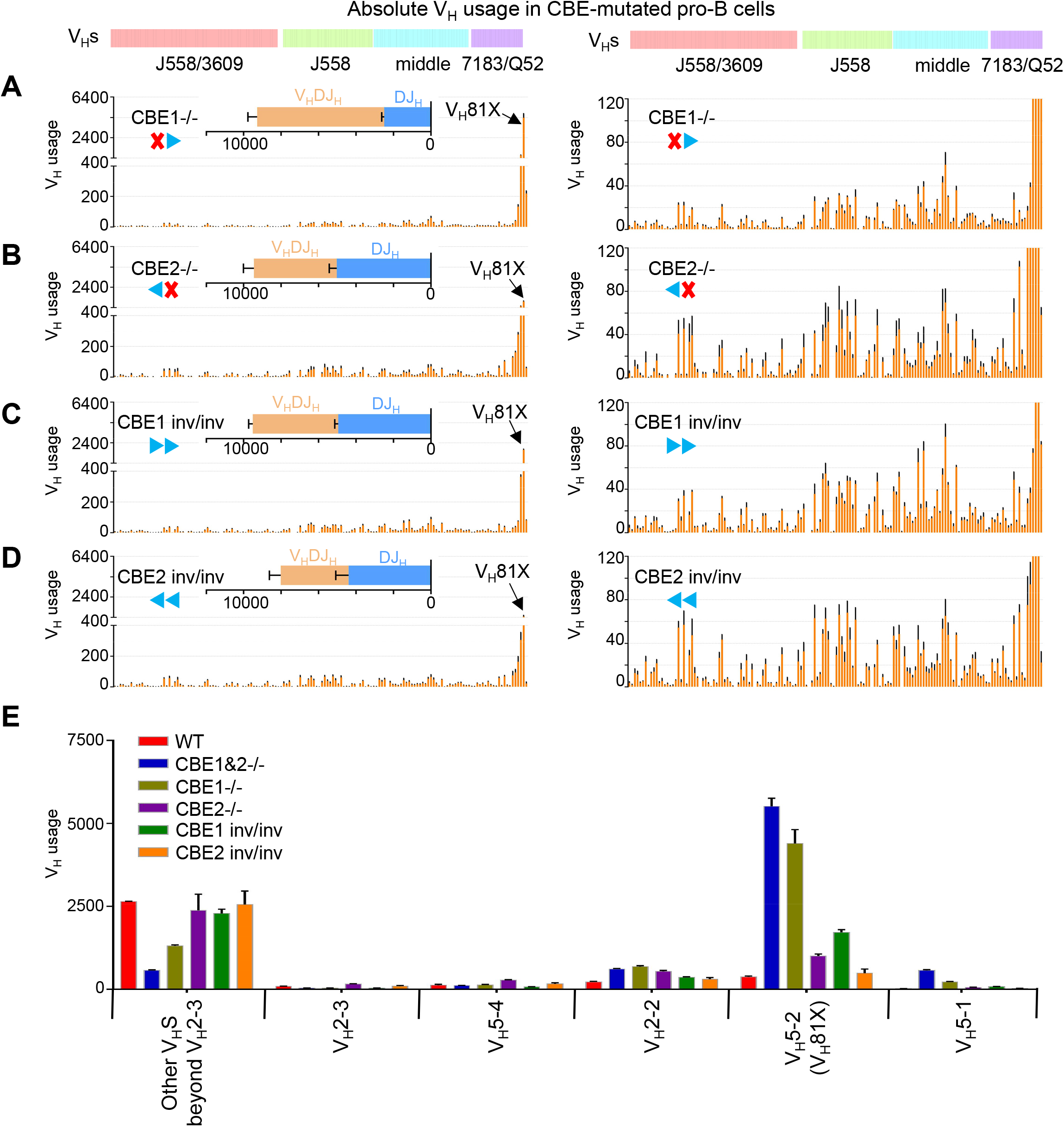
The mutation of IGCR1/CBEs alters V_H_s utilization in pro-B cells. (A-D) Each panel shows the utilization of V_H_s across the entire *Igh* locus in indicated IGCR1/CBEs mutated pro-B cells. The V_H_DJ_H_ and DJ_H_ junctions are shown in insets. (*n*=3 mice, mean±SEM; All HTGTS libraries are normalized to 78,091 total reads; see SI Appendix, Table S1). (E) V_H_s usage in WT and indicated IGCR1/CBEs mutated pro-B cells (*n*=3 mice, mean±SEM).

We performed 3C-HTGTS on RAG-deficient IGCR1/WT, IGCR1/CBE1&2^−/−^, IGCR1/CBE1^inv/inv^ and IGCR1/CBE2^inv/inv^ *v-Abl* lines with bait primers to the iEμ in the RC (SI Appendix, Fig. S6 A) and V_H_5-2 (SI Appendix, Fig. S6 B) locales. In CBE1^−/−^ pro-B cell populations, we observed robust, albeit somewhat diminished, interactions between the iEμ/RC bait and the proximal V_H_-CBEs compared to interactions in CBE1&2^−/−^ pro-B cells (SI Appendix, Fig. S6 A, CBE1^−/−^ vs. CBE1&2^−/−^). In CBE2^−/−^ pro-B cells we observed a modest increase in the interactions between the iE*μ*RC bait and the proximal V_H_-CBEs (SI Appendix, Fig. S6 A, CBE2^−/−^ vs. CBE1&2^−/−^). These findings indicate that IGCR1 CBEs play a cooperative role in impeding loop extrusion-mediated proximal V_H_-CBEs and RC interactions and, again, that CBE1 has a more dominant role. In contrast, when CBE1 or CBE2 was inverted, interactions between proximal V_H_-CBEs and iE*μ*/RC showed only very modest changes compared to those when IGCR1 CBEs are in normal orientation (SI Appendix, Fig. S6 A-B). These studies confirm unequivocally that CBE1 and CBE2 function synergistically to provide the full RAG scanning impediment activity of IGCR1 and that CBE1 provides the major portion of this activity. Moreover, reminiscent of the effects of V_H_5-2 RSS-associated CBE inversion as compared to complete inactivation on proximal V_H_5-2 rearrangements (18), IGCR1 CBEs in their normal orientation provide physiological levels of RAG-scanning impediment activity, but retain significant activity when inverted.

### Influence of 3’*Igh* CBEs on D-to-J_H_ and V_H_-to-DJ_H_ rearrangement patterns and levels

To investigate potential contributions of 3’*Igh* CBEs on RAG scanning, we deleted all 3’CBEs in RAG1-deficient C57BL/6 WAPL-degron *v-Abl* cells that contain a single copy of the *Igh* locus (22). For analyses, WT or 3’CBEs-deleted (3’CBE^−^) WAPL-degron *v-Abl* cells were arrested in G1 and then left untreated or treated with Dox plus IAA to degrade WAPL (22).

Subsequent introduction of RAG into untreated WT WAPL-degron *v-Abl* cells activated V(D)J recombination leading to robust D-to-J_H_ recombination, very low-level V_H_5-2 to DJ_H_ recombination, and extremely low-level upstream V_H_-to-DJ_H_ recombination (Fig. 4A and SI Appendix, Table S3), as *v-Abl* lines have high WAPL levels and RAG scanning is impeded at IGCR1(22). Dox plus IAA treatment completely depletes WAPL in these G1-arrested *v-Abl* lines, but they remain substantially viable (22). Introduction of RAG into WAPL-depleted WT *v-Abl* cells activated D-to-J_H_ rearrangement, but the level of DJ_H_ rearrangements was reduced 6.5-fold compared to that of untreated cells (Fig. 4I and SI Appendix, Table S3). This reduction was associated with dramatically decreased distal DFL16.1-J_H_ and, to a lesser degree, most other DJ_H_ absolute rearrangement levels (Fig. 4M, N). Notably, however, DQ52 rearrangement levels showed little change (Fig. 4M). WAPL-depletion also led to RAG-scanning and utilization of V_H_s across the 2.4 Mb V_H_ locus (Fig. 4 A,B and SI Appendix, Table S3) as expected (22). The absolute level of V_H_ DJ_H_ rearrangements in WAPL-depleted WT *v-Abl* cells was similar to that of the low level of proximal V_H_ rearrangements in untreated cells; but represented a 5.3-fold increase with respect to their fraction of DJ_H_ rearrangements (Fig. 4 I, J; see Discussion). In both untreated and WAPL-depleted 3’CBE^−^ lines, we observed a further 25% decrease in DJ_H_ rearrangement levels from their baseline levels. We observed a similar 25% decrease in V_H_DJ_H_ rearrangements in untreated 3’CBE^−^ lines and a nearly 50% decrease in WAPL-depleted 3’CBE-lines (compare Fig. 4 panels B and D, Fig.4 I-L). Despite the further reduction of V_H_DJ_H_ rearrangement levels in WAPL-depleted 3’CBE^−^ *v-Abl* cells, their relative V_H_ usage pattern was very similar to that of WAPL-depleted WT lines (compare Fig. 4, panel F and H).

**Figure 4.**
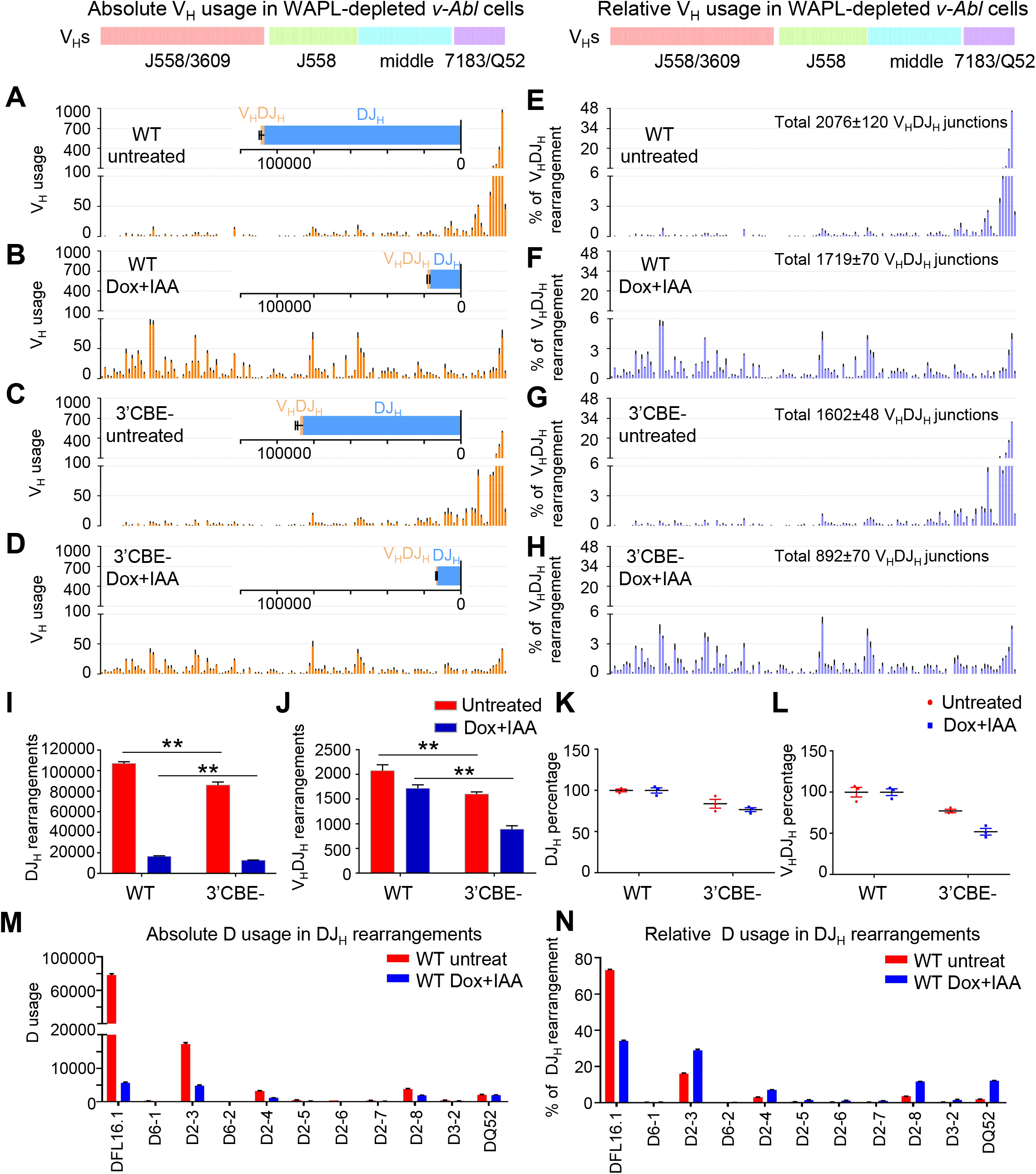
Role of 3’*Igh* CBEs in RC activity during loop extrusion. (A-D) Utilization of V_H_s across the entire *Igh* locus in (A, B) and 3’CBE^−^ (C, D) *v-Abl* cells with or without Dox/IAA treatments. The V_H_DJ_H_ and DJ_H_ junctions are shown in insets. (*n*=3 repeats from 3 independent clones, mean±SEM; all HTGTS libraries are normalized to 1,964,102 total reads; see SI Appendix, Table S3). (E-H) Relative percentage of V_H_s utilization normalized to the indicated V_H_DJ_H_ junction number (*n*=3 repeats, mean±SEM; percentages are plotted from the data of Fig. 4 A-D). (I-J) Absolute level of V_H_DJ_H_ and DJ_H_ rearrangements in WT and 3’CBE-lines (*n*=3 repeats, mean±SEM; *t*-test, *P*<0.01, **). (K-L) Relative percentages of V_H_DJ_H_ and DJ_H_ normalized to untreated or treated WT conditions. (M-N) Absolute (M) and relative (N) D usage in DJ_H_ rearrangements in untreated and WAPL-depleted WT *v-Abl* cells (relative percentage was normalized to 106,936 DJ_H_ junctions in untreated or 16,565 DJ_H_ junctions in WAPL-depleted cells; see SI Appendix, Table S3).

We performed 3C-HTGTS in RAG-deficient WT and 3’CBE^−^ *v-Abl* cells baiting from iEμ/RC (Fig. 5 A, B and SI Appendix Fig. S7 B, C, S8). These studies revealed, as previously described (22), that complete depletion of WAPL substantially increased a number of interaction peaks in the J558/3609, J558 and middle V_H_ regions (Fig. 5 A and SI Appendix, Fig. S8 Dox+IAA vs untreated; Peaks 1-13). Many of the sequences that contribute to these peaks were highly transcribed including the well-characterized PAIR elements (Fig. 5A and SI Appendix, Fig. S8; Peaks, 1-8). In contrast, peaks in the proximal 7183/Q52 region that are dominant in untreated *v-Abl* cells are mainly associated with proximal V_H_ RSS-CBEs and were significantly diminished by WAPL depletion (Fig. 5 A, and SI Appendix, Fig. S8, Dox+IAA vs untreated; Peaks 14-16). Notably, in 3’CBE^−^ cells, these same major interaction peaks, including those associated with proximal V_H_RSS-CBEs in untreated and those associated with the upstream V_H_ regions in treated and untreated cells, remain robust and largely correspond to those in the same locations as in the WT line (Fig. 5 A and SI Appendix, Fig. S8, 3’CBE^−^ vs WT). While the intensity of some peaks in the untreated or WAPL-depleted 3’CBE^−^ *v-Abl* cells were somewhat diminished compared those of the untreated or WAPL-depleted WT *v-Abl* cells, when viewed at high resolution they are clearly still associated with same transcriptional or CBE impediments (Fig. 5 A, and SI Appendix, Fig. S8). Upon the deletion of 3’*Igh* CBEs, multiple CBEs downstream of the 3’*Igh* CBEs appear to gain robust interactions with the iEμ/RC (Fig. 5B).

**Figure 5.**
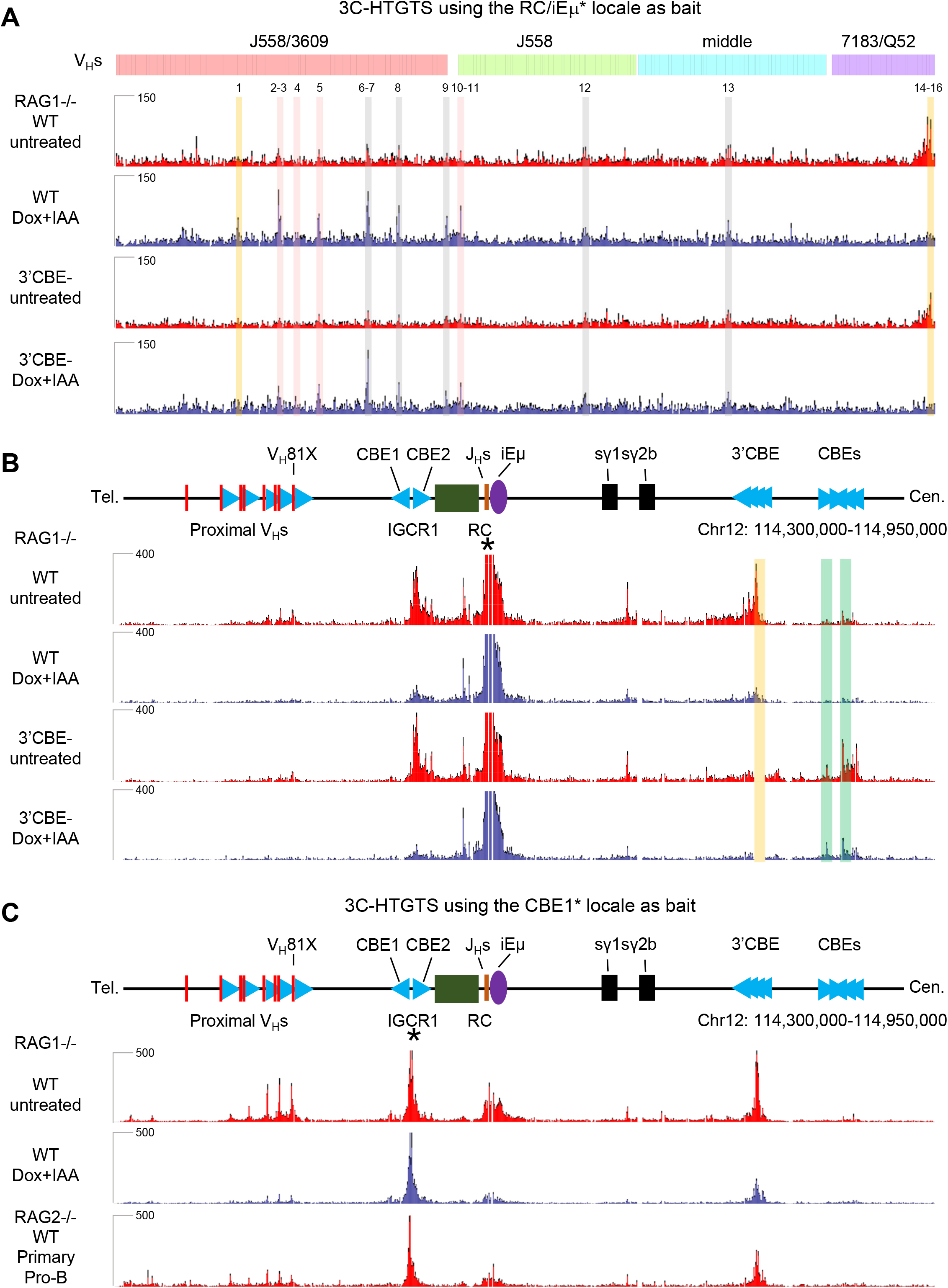
3C-HTGTS profiles at *Igh* locus bating from RC in WT and 3’CBE^−^ WAPL-degron *v-Abl* cells. (A) 3C-HTGTS signal counts at V_H_s domains of WT and 3’CBE^−^ RAG1-deficient *v-Abl* lines baiting from nRC/iEμ (*) with (red) or without (blue) Dox/IAA treatment. Each library was normalized to 160,314 total junctions (*n*=3 repeats from 3 independent clones, mean±SEM). 16 peaks across V_H_s region are called by MACS2 pipeline and highlighted in gray (peak 1, 6-9, 12-13 are called in WAPL-depleted WT *v-Abl* cells and WT pro-B cells), red (peaks 2-5, 10-11 are called in WAPL-depleted *v-Abl* cells), orange (peak1 is called only in WAPL-depleted WT *v-Abl* cells), green (peaks are called in 3’CBE^−^ *v-Abl* cells); Peaks 14-16 are present in untreated WT and 3’CBE^−^ *v-Abl* lines (See SI Appendix, Fig. S8). (B) Zoom-in 3C-HTGTS profiles of *Igh* locus from proximal V_H_S to 3’*Igh* CBEs. (C) 3C-HTGTS signal counts of *Igh* locus from proximal V_H_s to 3’*Igh* CBEs in RAG-deficient WT *v-Abl* cells (red), WAPL-depleted *v-Abl* cells (blue) and pro-B cells (red) baiting from IGCR1/CBE1 (*). Each library was normalized to 112,525 total junctions (*n*=3 repeats from 3 independent clones or 3 mice, mean±SEM).

Finally, WAPL-depletion also substantially diminishes interactions of IGCR1 CBE1 with both CBE-based (proximal V_H_s and 3’*Igh* CBEs and transcription-based (RC and *γ*1-*γ*2b enhancer) loop extrusion impediments (Fig. 5C).

## DISCUSSION

HTGTS-V(D)J-Seq analyses provided a deep analysis of V_H_ repertoires in primary WT and IGCR1/CBE1&2^−/−^ pro-B cells (Fig. 1). In addition, 3C-HTGTS-Seq analyses of iE*μ*/RC interactions across the *Igh* locus in RAG2-deficient primary WT and IGCR1/CBE1&2^−/−^ pro-B cells complemented HTGTS-V(D)J-Seq studies to reveal a likely mechanism by which ordered transition from D-to-J_H_ versus V_H_-to-DJ_H_ rearrangement is regulated (Fig. 2). Our overall findings suggest a model in which robust D-to-J_H_ rearrangements occur in WT pro-B cells before WAPL is sufficiently down-regulated to allow scanning to pass IGCR1 or proximal V_H_-associated CBEs. In CBE1&2^−/−^ pro-B cells, robust iE*μ*/RC interactions with proximal V_H_RSS-associated CBEs promote their robust rearrangements and suppress scanning to upstream V_H_s.

Our finding that low-level rearrangements of upstream V_H_s in CBE1&2^−/−^ pro-B cells have normal RAG-scanning patterns is consistent with these rearrangements occurring after WAPL-down-regulation in the cells that have not formed proximal V_H_DJ_H_ rearrangements on both alleles. In CBE1&2^−/−^*rag2*^*-/-*^ pro-B cells, lack of V(D)J recombination upon WAPL down-regulation allows extrusion of the iE*μ*/RC to continue into the upstream V_H_ domain where it reaches normal impediments in a substantial fraction of the cells. In this context, the relatively robust contribution of V_H_5-2 and immediately upstream V_H_s to the WT pro-B repertoire indicates that these V_H_s are dominantly utilized in normal pro-B cells until WAPL levels are sufficiently down-regulated Finally, in support of this model, proximal V_H_s are poorly utilized in *v-Abl* cells in which RAG is introduced after complete WAPL-depletion (22) (Fig 4).

Our findings on the impact of WAPL-depletion on chromatin interactions and RAG scanning activity support a model in which 3’*Igh* CBEs reinforce RC activity during *Igh* V(D)J recombination (SI Appendix, Fig. S9) (1). In WT *v-Abl* cells with high WAPL expression, the RC robustly interacts with the downstream *γ*1-*γ*2b enhancer and the 3’*Igh* CBEs, which, as proposed (1), could reinforce its loop-extrusion impediment activity (Fig. 5B). Complete WAPL depletion in *v-Abl* cells substantially diminishes downstream RC interactions (Fig. 5B). In addition, interactions of the IGCR1/CBE1 with the RC, as well as with the *γ*1-*γ*2b enhancer and 3’*Igh* CBEs, are also greatly diminished in WAPL-depleted *v-Abl* cells (Fig. 5C), consistent with WAPL-depletion diminishing transcription-based RC impediment activity. In contrast, in WT pro-B cells, in which WAPL levels are modestly reduced (44), these interactions are relatively robust (Fig. 2B). We propose that the 6.5-fold decrease in DJ_H_ rearrangements in WAPL-depleted *v-Abl* cells results from decreased RC impediment activity (Fig. 4I and 5B), as previously proposed for a similar reduction in V_*κ*_J_*κ*_ rearrangements upon WAPL depletion in this *v-Abl* line (22). Notably, WAPL-depletion reduced rearrangement levels of all Ds, other than DQ52, leading to DQ52 contributing more substantially to residual DJ_H_ rearrangements (Fig. 4N). A similar overall trend was obtained when previously reported data that employed J_H_ 1-4 baits (22) was analyzed for absolute levels, as well as relative percentages (SI Appendix, Fig. S7D, E; Table S4). We propose that DQ52 recombination, after WAPL depletion, may be less affected, because it accesses RAG by diffusion (versus scanning) from its RC location (24).

Finally, diverse V_H_DJ_H_ rearrangements in WAPL-depleted cells occurred at a similarly low, absolute level to those of proximal V_H_s in untreated cells. However, V_H_ rearrangements in WAPL-depleted cells contributed to a 5.3-fold increase in the proportion of V_H_DJ_H_/DJ_H_ rearrangements (Fig. 4I, J). This latter finding may reflect increased levels of V_H_DJ_H_ recombination in WAPL-depleted *v-Abl* cells compensating for reduced RC activity (Fig 4). However, the net effect is that overall V(D)J recombination levels are much lower in WAPL-depleted *v-Abl* lines than in BM pro-B cells as reported (22).

Deletion of 3’*Igh* CBEs decreased DJ_H_ and V_H_DJ_H_ rearrangement levels in both untreated and WAPL-depleted *v-Abl* cells. These decreases support the model that 3’ *Igh* CBEs contribute to reinforcing RC impediment activity (SI Appendix, Fig. S9). In this regard, the similarity in V_H_ usage patterns across the V_H_ locus, despite the nearly 50% decrease in V(D)J junctions, in WAPL-depleted 3’CBE^−^ versus untreated 3’CBE^−^ *v-Abl* cells indicates that the impact on V(D)J recombination in these WAPL-depleted lines affects V(D)J recombination activity *per se*, but apparently not scanning patterns. We note that the extent to which the 3’ *Igh* CBEs reinforce RC activity may be compensated, in its absence, by interactions of the RC with downstream CBEs; it is also notable that these downstream CBE interactions are diminished by complete WAPL down-regulation (Fig. 5B). Such compensatory activity of downstream CBEs, in the absence of 3’ *Igh* CBEs, was also implicated in the context of *Igh* class switch recombination (50). Finally, normal pro-B cells do not completely down-regulate WAPL levels (44), which may contribute to preserving 3’*Igh* CBEs/RC interactions and RC activity in these cells. In this regard, WT and 3’CBE^−^ pro-B cells have indistinguishable *Igh* V(D)J recombination patterns (22).

When the RC-interacting peaks found in the 129SV pro-B cells (Fig. 2A), and C57BL/6 *v-Abl* cells (Fig. 5A), a number of the peaks C57BL/6 Peaks 6-9, 12-13 (Fig. 5A) are shared; but others are unique due to the significant differences in the V_H_ loci in these two cell types. Major peaks in J558/3609, distal V_H_s region, are often associated with PAIR elements (Peaks 1, 4, 6, 8-10 in Fig. 2A and peaks 1-8 in Fig. 5A), while other peaks in J558/3609, J558, and middle V_H_ regions (Peaks 2, 3, 5, 7, 11-15 in Fig. 2A and peaks 9-13 in Fig. 5A) are associated with transcription or CBE binding motifs. These observations support the notion that when WAPL is down regulated, upon the neutralization of IGCR1, various transcription sites and CBEs still form sufficiently active loop extrusion impediments to promote interactions with the RC during RAG scanning of upstream V_H_s locus sequences.

## METHODS

### Mice

Wildtype 129SV mice were purchased from Charles River Laboratories International. RAG2-deficient mice in 129SV background were purchased from Taconic. All animal experiments were performed under protocols approved by the Institutional Animal Care and Use Committee of Boston Children’s Hospital.

### Generation of IGCR1 CBE-inversion mice

A previously described pLNTK targeting vector (49) containing inversion mutations of the 20-bp CBE1 and corresponding upstream activating sequence (WT sequence: 5’- TGCTTCCCCCTTGTGGCCATGAGCATTACTGCA-3’; inverted: 5’- TGCAGTAATGCTCATGGCCACAAGGGGGAAGCA-3’); or the 19-bp CBE2 (WT sequence: 5’-TCTCCACAAGAGGGCAGAA-3’; inverted sequence: 5’-TTCTGCCCTCTTGTGGAGA- 3’) sites within IGCR1 were electroporated into TC1 ES cells. Successfully targeted clones with CBE1 or CBE2 inversion integration were assessed by Southern blot analyses using StuI-digested (13.9 kb untargeted; 10 kb targeted) or SpeI-digested (16.3 kb untargeted; 12.7 kb targeted) genomic DNA with appropriate probes. Two independently targeted clones containing each inversion mutation were subjected to adenovirus mediated Cre deletion to remove the NeoR gene, karyotyped, and injected for germline transmission. Homozygous mice were generated through breeding and genotype was confirmed by PCR genotyping (Primer sequences are listed in Table S5).

### Generation of *v-Abl* cell lines

The WT *v-Abl*-kinase-transformed pro-B cell line (*v-Abl* pro-B cells) was derived by retroviral infection of BM pro-B cells derived from *rag*2^−/−^ mice as described (48). IGCR1 mutated RAG2-deficient *v-Abl* lines were established by breeding each IGCR1 mutant mice with *rag*2^−/−^ germline mice to generate RAG2-deficient homozygous IGCR1 mutated mice (i.e. IGCR1/CBE1^−/−^*rag*2^−/−^), and deriving *v-Abl* lines as described (48). All these mutations were confirmed by PCR genotyping. 3’*Igh* CBEs deletion in single *Igh* WAPL-degron *v-Abl* (22) were generated by designed sgRNAs and screened by PCR. The sequence of all sgRNAs and oligos used are listed in Table S5.

### HTGTS-V(D)J-seq and data analyses

Pro-B cells used in HTGTS-V(D)J-seq experiments were purified from WT or IGCR1 mutated mice as described (53). 2ug pro-B cell genomic DNA were used to generate each library. The sequence of the J_H_4 coding end primer (129SV background) used to generate HTGTS-V(D)J-seq libraries is listed in Table S5. HTGTS-V(D)J-seq libraries were prepared as described (24). HTGTS-V(D)J-seq libraries were sequenced using paired-end 300-bp sequencing on a Mi-Seq (Illumina) machine. The WT 129SV pro-B cells data shown in Fig.1 and Fig. S1-2 was extracted from a prior publication (GSM2183881-GSM2183883) (53). All libraries were normalized to total reads (junctions+germine reads) or junctions across a given locus, and the V_H_DJ_H_ and DJ_H_ junctions were described in Table S1, S2. When normalized to total reads, all libraires were normalized to the smallest libraries from the same batch of experiments. The number of normalized reads or junctions is indicated in the figure and figure legends. D usage from the V_H_DJ_H_ joins was analysed via the VDJ_annotation pipeline. Productive and nonproductive V_H_DJ_H_ joins were analyzed via VDJ_productivity_annotation pipeline (see ‘code availability’).

RAG recombination and treatment of WAPL-degron *v-Abl* cells was performed as described (22), and the J_H_4 coding end primer (C57BL/6 background) used to generate HTGTS-V(D)J-seq libraries. All libraries were normalized to total reads or junctions, and the V_H_DJ_H_ and DJ_H_ junctions were described in Table S3. D usage from the V_H_DJ_H_ joins was analyzed by the VDJ_annotation pipeline.

### 3C-HTGTS and data analyses

RAG2-deficient pro-B cells for 3C-HTGTS were purified and cultured as described (19). Cycling or G1-arrested RAG2-deficient *v-Abl* pro-B cells for 3C-HTGTS were prepared as described (18). Treatment of WAPL-degron *v-Abl* cells was performed as described (22). 3C-HTGTS was performed as described (18). Briefly, 10 million cells were crosslinked with 2% (v/v) formaldehyde for 10 min at RT. Cells were lysed in 50 mM Tris-HCl, pH 7.5, containing 150 mM NaCl, 5 mM EDTA, 0.5% NP-40, 1% Triton X-100 and protease inhibitors (Roche, #11836153001). Nuclei were digested with 700 units of NlaIII (NEB, #R0125) restriction enzyme at 37℃ overnight, followed by ligation (T4 DNA ligase NEB M0202L) at 16℃overnight. Crosslinks were reversed and samples were treated with Proteinase K (Roche, #03115852001) and RNase A (Invitrogen, #8003089) prior to DNA precipitation. 3C-HTGTS libraries were generated using LAM-HTGTS (56), and primers are listed in Table S5.

3C-HTGTS libraries were sequenced using paired-end 150-bp sequencing on a Next-seq550 (Illumina) or paired-end 300-bp sequencing on a Mi-Seq (Illumina) machine. Data were processed as described previously (18). In addition, the PCR artificial junctions at Chr12: 114,692,680 in Fig. S6 A were removed from the total junctions. The junctions from Chr12 were extracted and counted for normalization. All 3C-libraires were normalized to the smallest libraries from the same batch of experiments. The number of normalized junctions is indicated in the figure legends. For peak analysis, 3C-HTGTS profiles were analyzed by MACS2 pipeline to call robust interaction peaks (macs2 bdgpeakcall -c20 -l400 -g1000 was used for pro-B 3C-HTGTS in Fig. 2 and macs2 bdgpeakcall -c30 -l400 -g1000 was used for *v-Abl* 3C-HTGTS in Fig. 5). The peaks that showed >2-fold intensity change were annotated as unique peaks of indicated condition.

## Data availability

HTGTS-V(D)J-seq, 3C-HTGTS and GRO-seq sequencing data reported in this study are available through GEO (GSExxxx). All study data are included in the article and/or supporting information. Previously published data were used for this work [GSE151910 (22) and GSE821126 (53)].

## Code availability

3C-HTGTS and HTGTS-V(D)J-seq data were processed through published pipelines (http://robinmeyers.github.io/transloc_pipeline/). D usage in V_H_DJ_H_ joins was processed via a custom pipeline (https://github.com/Yyx2626/VDJ_annotation/). Productive and non-productive junctions were processed via another pipeline (https://github.com/Yyx2626/VDJ_annotation/).

## ACKNOWLEDGMENTS

We thank members of the Alt laboratory for stimulating discussions. This work was supported by the National Institutes of Health Grant R01AI020047 to F.W.A. and Grant F31-AI117920 to S.G.L.; Z.B. and H-Q.D. were supported in part by Cancer Research Institute Irvington Fellowships. F.W.A. is an investigator of the Howard Hughes Medical Institute.

