## Supplementary Information for "Contribution of the IGCR1 regulatory element and the 3*’Igh* CBEs to Regulation of *Igh* V(D)J Recombination"

**This PDF file includes:**

Figures S1 to S9

Tables S1 to S5

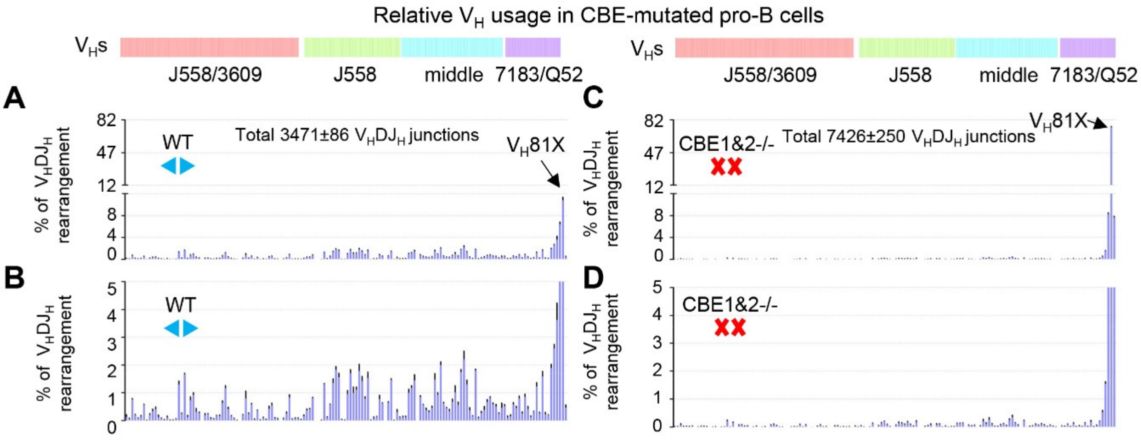


**Figure S1. Relative $V_H$ utilization in WT and CBE1&2$^{-/-}$ pro-B cells.** (A-D) Relative percentage of $V_H$s utilization normalized to the indicated $V_HDJ_H$ junction number in WT (A, B) and CBE1&2$^{-/-}$ (C, D) pro-B cells (*n*=3 mice, mean±SEM; percentages are plotted from the data of Fig. 1B, D).

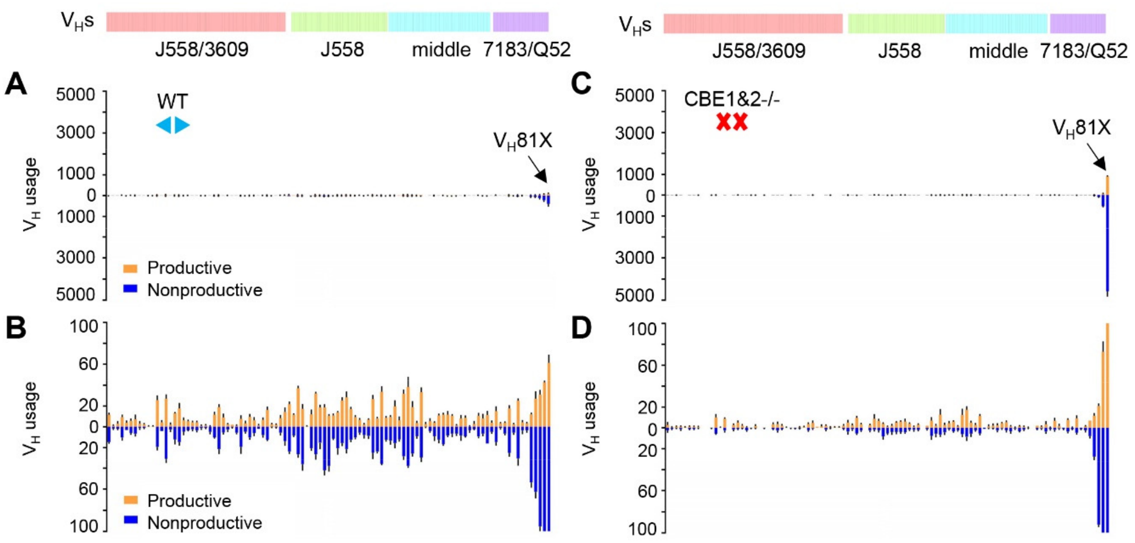


**Figure S2. Productive and nonproductive rearrangements in WT and CBE1&2$^{-/-}$ pro-B cells.** (A-D) Each panel shows the productive (orange) or nonproductive (blue) VHs usage in WT and CBE1&2$^{-/-}$ pro-B cells. Productive and nonproductive rearrangements are analyzed using a custom pipeline (*n*=3 mice, mean±SEM; see Methods and SI Appendix, Table S2).

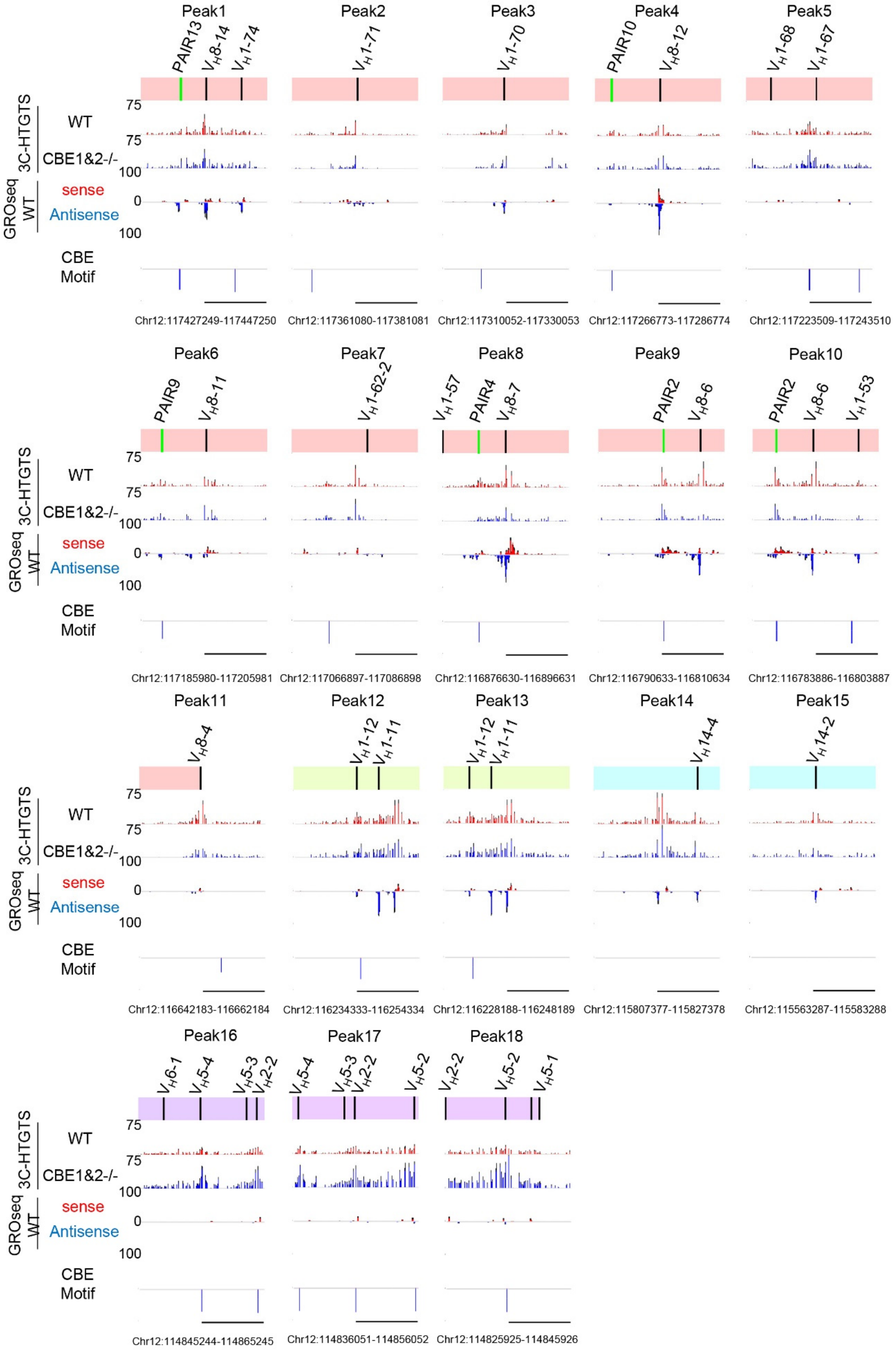
Peak1
Peak2
Peak3
Peak4
Peak5
3C-HTGTS
WT
CBE1&2-/-
GROseq WT
sense
Antisense
CBE Motif
Chr12:117427249-117447250
Chr12:117361080-117381081
Chr12:117310052-117330053
Chr12:117266773-117286774
Chr12:117223509-117243510
Peak6
Peak7
Peak8
Peak9
Peak10
Chr12:117185980-117205981
Chr12:117066897-117086898
Chr12:116876630-116896631
Chr12:116790633-116810634
Chr12:116783886-116803887
Peak11
Peak12
Peak13
Peak14
Peak15
Chr12:116642183-116662184
Chr12:116234333-116254334
Chr12:116228188-116248189
Chr12:115807377-115827378
Chr12:115563287-115583288
Peak16
Peak17
Peak18
Chr12:114845244-114865245
Chr12:114836051-114856052
Chr12:114825925-114845926

**Figure S3. The major RC interactions, transcriptions, and CBE motifs in cultured RAG2-deficient primary pro-B cells.** Zoom-in profiles of 3C-HTGTS, GRO-seq, CBE motif sites signals for ±10kb regions of 18 representative peaks in Fig. 2A from WT (red) and CBE1&2$^{-/-}$ (blue) cultured RAG2-deficient primary pro-B cells (*n*=3 mice, mean±SEM). PAIR elements (green bars) located in each peak are also shown above. Peaks 1-9, 12-15 are called in WT and CBE1&2$^{-/-}$; Peaks 10-11 are called only in WT; Peaks 16-18 are called only in CBE1&2$^{-/-}$.

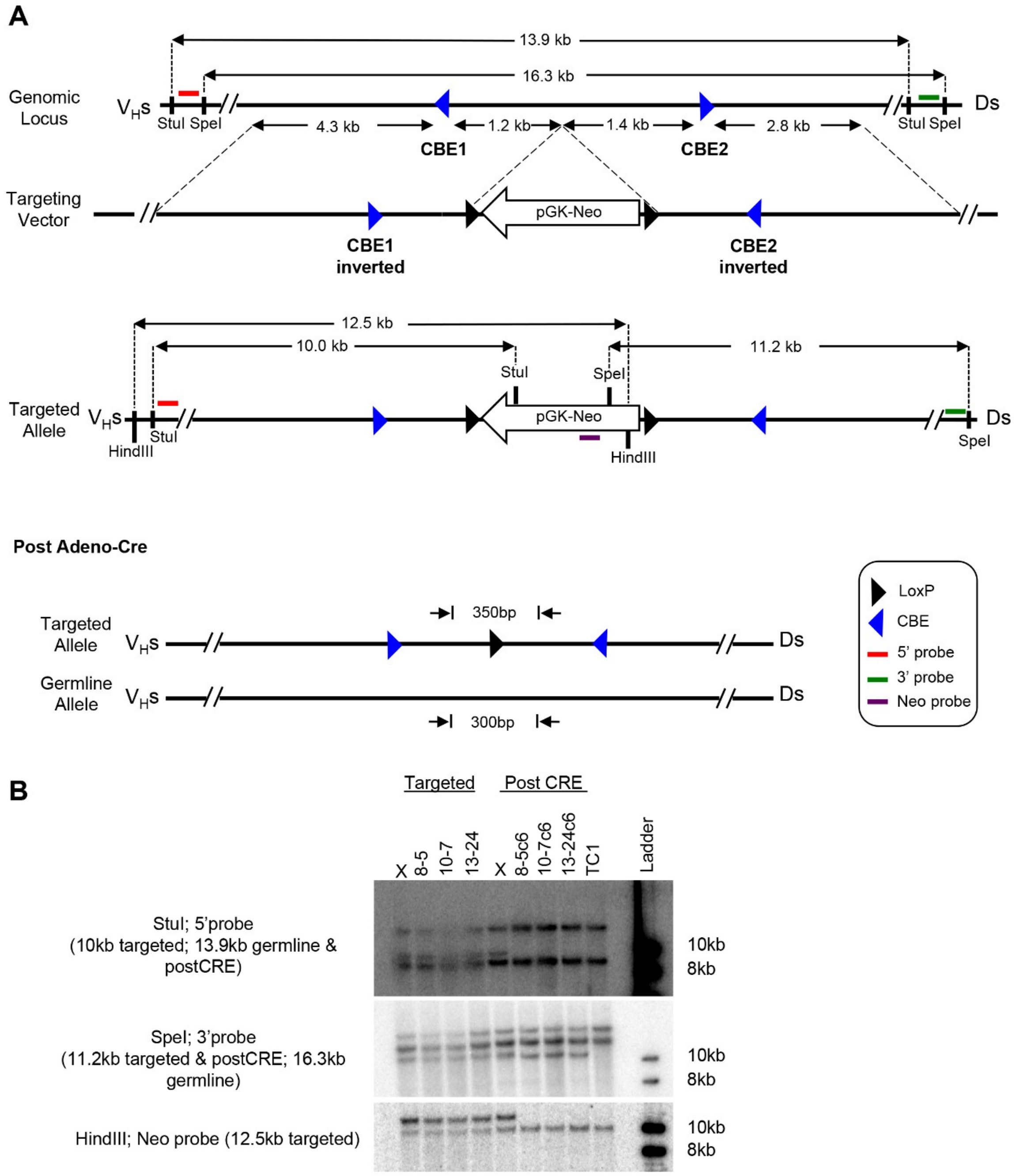


**Figure S4. Generation and validation of CBE1 or CBE2 inversion mice.** (A) Schematic diagram of the targeting strategy to generate CBE1 or CBE2 inversion in ES cells. Green and red lines indicate positions of probes used to confirm the insertion (see Methods). (B) Selection of inversion positive ES Cell clones by Southern blot (TC1: parental ES cell clone; 8-5c6, 10-7c6 and 13-24c6 indicate ES cell clones with insertion replacements after Adeno-Cre).

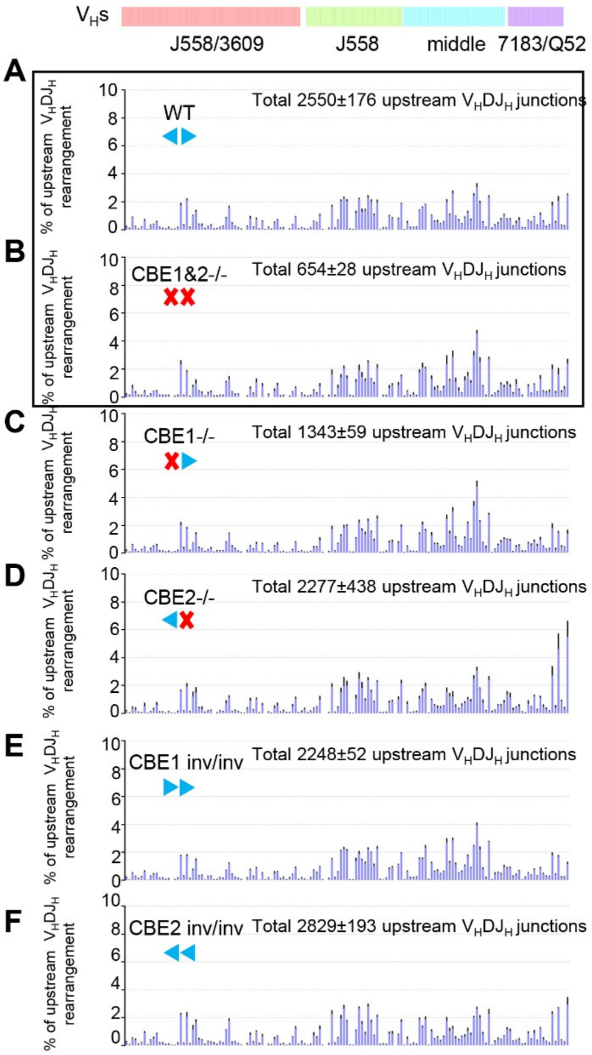


**Figure S5 Relative $V_H$ utilization in WT and indicated IGCR1/CBEs mutated pro-B cells.** (A-F) Each panel shows the relative percentage of upstream $V_H$s beyond the five most proximal $V_H$s normalized to the indicated $V_HDJ_H$ junction number. Upstream $V_H$s junctions are extracted from the data of SI Appendix, Fig. S5 A-F. Panel A, B are reproduced from Fig. 1H, I (*n*=3 mice, mean±SEM).

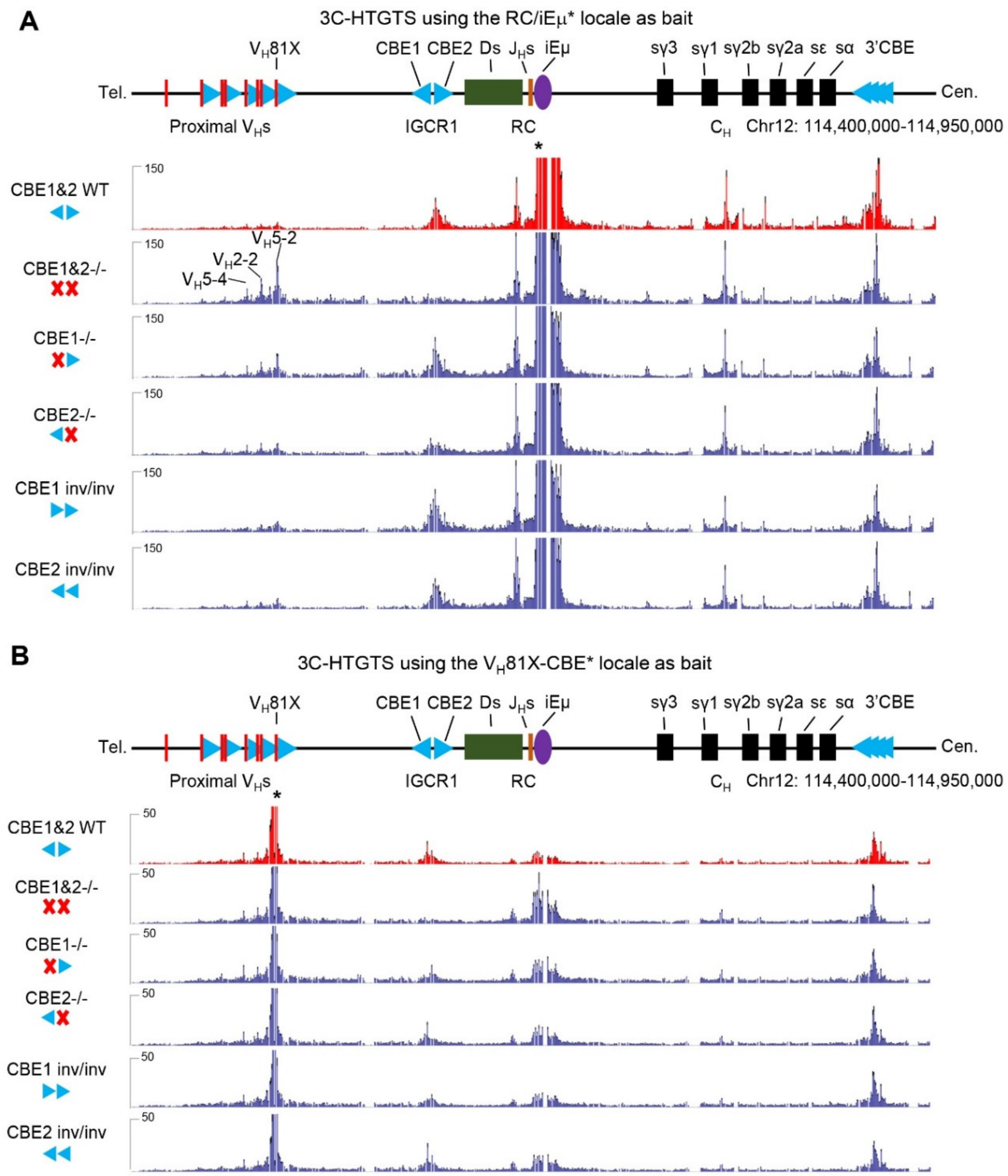


**Figure S6. 3C-HTGTS profiles bating from RC and $V_H$5-2-CBE in IGCR1/WT and IGCR1/CBEs mutated *v-Abl* cells.** (A) 3C-HTGTS signal counts of IGCR1/WT (red) and IGCR1/CBEs mutated (blue) RAG2-deificient *v-Abl* lines baiting from nRC/iEμ (*). Each library was normalized to 70,566 total junctions (*n*=6 repeats from 2 independent clones, mean±SEM). (B) 3C-HTGTS signal counts of IGCR1/WT (red) and IGCR1/CBEs mutated (blue) RAG2-deificient *v-Abl* lines baiting from $V_H$5-2-CBE (*). Each library was normalized to 14,018 total junctions (*n*=6 repeats from 2 independent clones, mean±SEM).

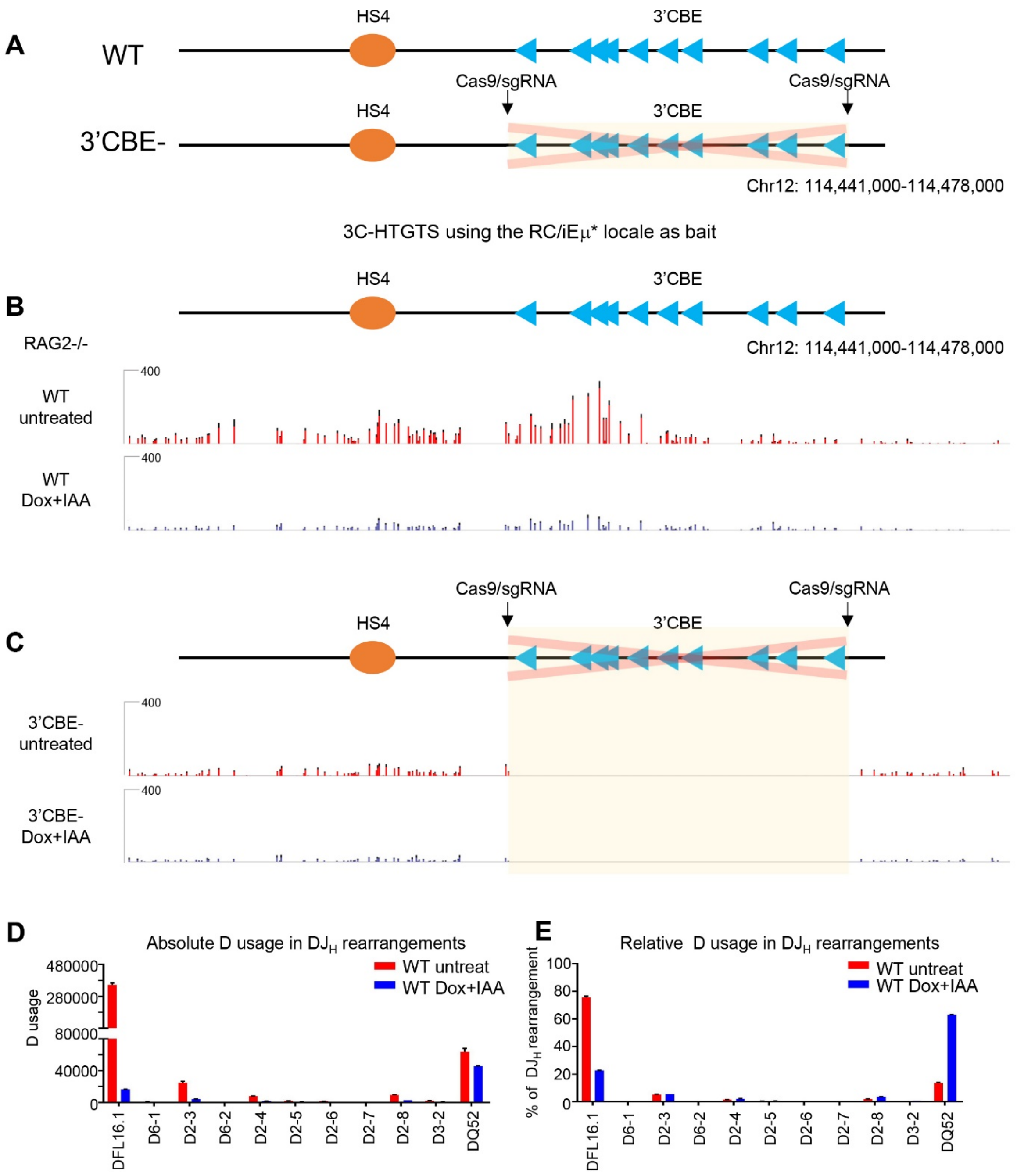


**Figure S7. Role of 3'*Igh* CBEs in RC activity during loop extrusion.** (A) Generation of 3'CBE⁻ by Cas9/sgRNAs in single *Igh* WAPL-degron *v-Abl* lines. (B-C) Zoom-in 3C-HTGTS profiles of 3'*Igh* CBE in WT (B) and 3'CBE⁻ (C) WAPL-degron *v-Abl* cells with or without Dox/IAA treatments (Chr12: 114,441,000-114,478,000; see Fig. 5 B). (D, E) Absolute (D) and relative (E) D usage in D to $J_H$1-4 rearrangements in untreated and WAPL-depleted WT *v-Abl* cells. HTGTS-V(D)J-Seq data was extracted from a prior study in which employed primers specific to each of the four mouse $J_H$s (22: GSM4593296-GSM4593303). HTGTS-V(D)J-Seq libraries were generated using combined $J_H$1-4 primers and relative percentage was normalized to 465,530 $DJ_H$ absolute junctions in untreated or 72,283 $DJ_H$ absolute junctions in WAPL-depleted cells. The general trends for absolute and relative D usage based on utilizing combined primers for all four $J_H$s are similar to our findings based on utilization $J_H$4 primer (Fig. 4 M, N). The absolute usage data for all four $J_H$s in these experiments is shown in Table S4.

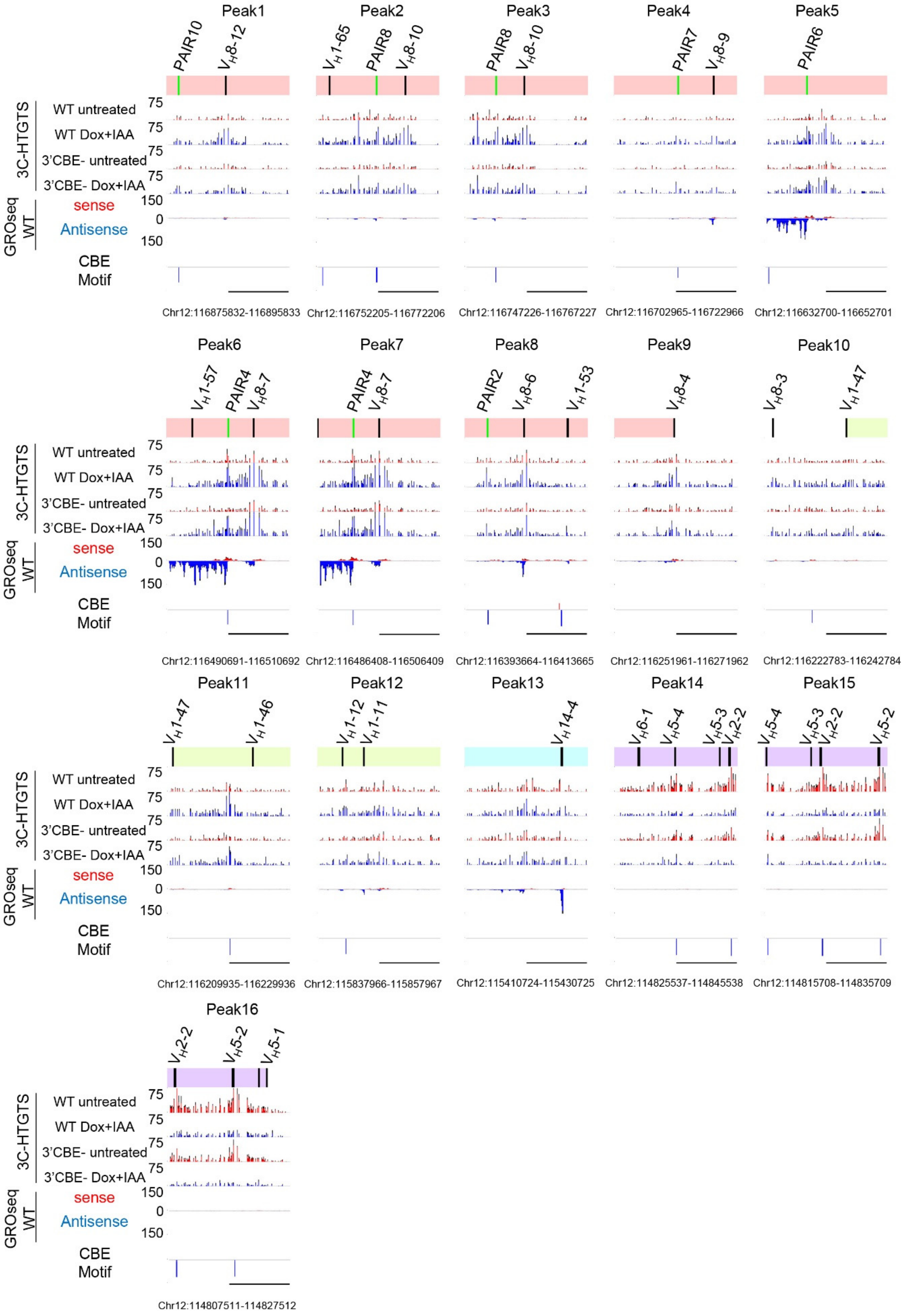

Peak1
Peak2
Peak3
Peak4
Peak5
Peak6
Peak7
Peak8
Peak9
Peak10
Peak11
Peak12
Peak13
Peak14
Peak15
Peak16
3C-HTGTS
WT untreated
WT Dox+IAA
3'CBE- untreated
3'CBE- Dox+IAA
GROseq WT
sense
Antisense
CBE Motif
75
150
0
PAIR10
PAIR8
PAIR7
PAIR6
PAIR4
PAIR2
Chr12:116875832-116895833
Chr12:116752205-116772206
Chr12:116747226-116767227
Chr12:116702965-116722966
Chr12:116632700-116652701
Chr12:116490691-116510692
Chr12:116486408-116506409
Chr12:116393664-116413665
Chr12:116251961-116271962
Chr12:116222783-116242784
Chr12:116209935-116229936
Chr12:115837966-115857967
Chr12:115410724-115430725
Chr12:114825537-114845538
Chr12:114815708-114835709
Chr12:114807511-114827512

**Figure S8. The major RC interactions, transcriptions, and CBE motifs in WT, 3'CBE$^{-}$ and 3'CBE$^{inv}$ WAPL-depleted *v-Abl* cells.** Zoom-in profiles of 3C-HTGTS, GRO-seq, CBE motif sites signals for ±10kb regions of 16 representative peaks in Fig. 5A from WT and 3'CBE$^{-}$ WAPL-depleted *v-Abl* cells (*n*=3 repeats from 3 independent clones, mean±SEM). PAIR elements (green bars) located in each peak are also shown above. Peaks 1, 6-9, 12-13 are shared in WAPL-depleted WT *v-Abl* cells and WT pro-B cells (SI Appendix, Fig. S3); Peaks 2-5, 10-11 are called in WAPL-depleted *v-Abl* cells; Peak1 is called only in WAPL-depleted WT *v-Abl* cells; Peaks 14-16 are only present in untreated WT and 3'CBE$^{-}$ *v-Abl* lines. The reference GRO-seq sequencing data was extracted from GSM4593308-GSM4593311.

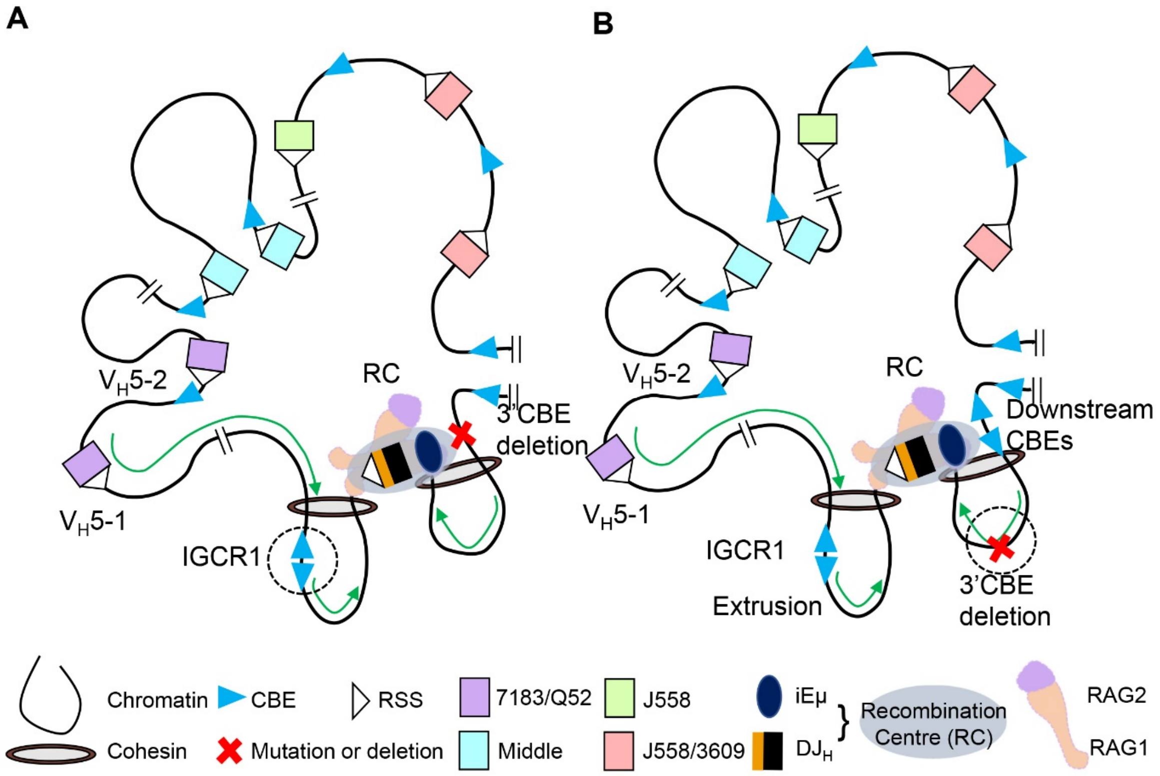


**Figure S9. Roles of IGCR1/CBEs and 3'*Igh* CBEs in *Igh* loop-extrusion mediated RAG scanning.** (A) 3'*Igh* CBEs reinforce the impediment activity of RC and promote the optimal RAG scanning. (B) Downstream CBEs appear to gain robust interactions with the RC upon the deletion of 3'*Igh* CBEs.

**Supplementary Table 1 | Absolute utilization of Igh VH and D segments in WT and IGCR1/CBEs mutated pro-B cells.**

| | | WT (n=3) | | | | | CBE1&2 -/- (n=3) | | | | | CBE1 -/- (n=3) | | | | | CBE2 -/- (n=3) | | | | | CBE1 inv/inv (n=3) | | | | | CBE2 inv/inv (n=3) | | | | |
|---|---|---|---|---|---|---|---|---|---|---|---|---|---|---|---|---|---|---|---|---|---|---|---|---|---|---|---|---|---|---|---|
| **VH domains** | | #1 | #2 | #3 | Average | s.d. | #1 | #2 | #3 | Average | s.d. | #1 | #2 | #3 | Average | s.d. | #1 | #2 | #3 | Average | s.d. | #1 | #2 | #3 | Average | s.d. | #1 | #2 | #3 | Average | s.d. |
| | Ighv1-86P | 9 | 5 | 0 | 5 | 5 | 2 | 0 | 1 | 1 | 1 | 4 | 2 | 3 | 3 | 1 | 7 | 6 | 0 | 4 | 4 | 4 | 9 | 1 | 5 | 4 | 2 | 6 | 3 | 4 | 2 |
| | Ighv1-85 | 4 | 2 | 0 | 2 | 2 | 1 | 0 | 2 | 1 | 1 | 1 | 1 | 2 | 1 | 1 | 1 | 1 | 0 | 1 | 1 | 0 | 1 | 4 | 2 | 2 | 0 | 3 | 2 | 2 | 2 |
| | Ighv1-84 | 24 | 30 | 25 | 26 | 3 | 2 | 0 | 13 | 5 | 7 | 5 | 8 | 7 | 7 | 2 | 22 | 5 | 5 | 11 | 10 | 11 | 16 | 13 | 13 | 3 | 12 | 20 | 6 | 13 | 7 |
| | Ighv1-83P | 8 | 8 | 5 | 7 | 2 | 3 | 1 | 2 | 2 | 1 | 6 | 3 | 2 | 4 | 2 | 14 | 10 | 1 | 8 | 7 | 0 | 8 | 4 | 4 | 4 | 11 | 17 | 6 | 11 | 6 |
| | Ighv1-82 | 2 | 4 | 3 | 3 | 1 | 1 | 3 | 1 | 2 | 1 | 3 | 3 | 0 | 2 | 2 | 6 | 0 | 2 | 3 | 3 | 5 | 3 | 0 | 3 | 3 | 5 | 4 | 2 | 4 | 2 |
| | Ighv1-81 | 3 | 4 | 8 | 5 | 3 | 3 | 0 | 1 | 1 | 2 | 1 | 3 | 2 | 2 | 1 | 3 | 0 | 0 | 1 | 2 | 5 | 6 | 3 | 5 | 2 | 6 | 5 | 5 | 5 | 1 |
| | Ighv1-80 | 18 | 22 | 19 | 20 | 2 | 2 | 4 | 2 | 3 | 1 | 9 | 5 | 5 | 6 | 2 | 34 | 5 | 4 | 14 | 17 | 16 | 12 | 15 | 14 | 2 | 18 | 33 | 19 | 23 | 8 |
| | Ighv1-79P | 0 | 1 | 0 | 0 | 1 | 0 | 0 | 0 | 0 | 0 | 0 | 0 | 0 | 0 | 0 | 1 | 0 | 0 | 0 | 1 | 1 | 2 | 0 | 1 | 1 | 2 | 1 | 0 | 1 | 1 |
| | Ighv1-78 | 10 | 6 | 5 | 7 | 3 | 1 | 0 | 1 | 1 | 1 | 2 | 6 | 2 | 3 | 2 | 1 | 2 | 2 | 2 | 1 | 7 | 5 | 10 | 7 | 3 | 13 | 14 | 10 | 12 | 2 |
| | Ighv1-77 | 16 | 8 | 11 | 12 | 4 | 2 | 3 | 3 | 3 | 1 | 3 | 6 | 5 | 5 | 2 | 11 | 5 | 2 | 6 | 5 | 5 | 8 | 19 | 11 | 7 | 19 | 16 | 7 | 14 | 6 |
| | Ighv1-76 | 10 | 21 | 11 | 14 | 6 | 4 | 1 | 1 | 2 | 2 | 9 | 8 | 3 | 7 | 3 | 24 | 20 | 4 | 16 | 11 | 11 | 7 | 16 | 11 | 5 | 23 | 27 | 15 | 22 | 6 |
| | Ighv1-75 | 9 | 1 | 3 | 4 | 4 | 1 | 0 | 2 | 1 | 1 | 3 | 3 | 0 | 2 | 2 | 5 | 3 | 0 | 3 | 3 | 3 | 5 | 4 | 4 | 1 | 3 | 6 | 2 | 4 | 2 |
| | Ighv1-74 | 3 | 4 | 3 | 3 | 1 | 0 | 0 | 0 | 0 | 0 | 4 | 2 | 4 | 3 | 1 | 3 | 3 | 0 | 2 | 2 | 2 | 7 | 6 | 5 | 3 | 1 | 10 | 5 | 5 | 5 |
| | Ighv1-73P | 0 | 0 | 2 | 1 | 1 | 0 | 1 | 0 | 0 | 1 | 1 | 1 | 1 | 1 | 0 | 0 | 0 | 0 | 0 | 0 | 0 | 1 | 1 | 1 | 1 | 2 | 2 | 1 | 2 | 1 |
| | Ighv8-13 | 4 | 0 | 7 | 4 | 4 | 0 | 0 | 2 | 1 | 1 | 1 | 3 | 1 | 2 | 1 | 3 | 3 | 2 | 3 | 1 | 0 | 0 | 1 | 0 | 1 | 1 | 4 | 3 | 3 | 2 |
| | Ighv1-72 | 1 | 1 | 0 | 1 | 1 | 0 | 0 | 0 | 0 | 0 | 0 | 0 | 1 | 0 | 1 | 0 | 0 | 0 | 0 | 0 | 1 | 1 | 0 | 1 | 1 | 0 | 2 | 0 | 1 | 1 |
| | Ighv1-71 | 1 | 1 | 0 | 1 | 1 | 0 | 0 | 1 | 0 | 1 | 1 | 0 | 1 | 1 | 1 | 0 | 0 | 0 | 0 | 0 | 0 | 3 | 1 | 1 | 2 | 0 | 3 | 0 | 1 | 2 |
| | Ighv1-70P | 2 | 3 | 1 | 2 | 1 | 1 | 0 | 1 | 1 | 1 | 0 | 2 | 4 | 2 | 2 | 4 | 4 | 1 | 3 | 2 | 4 | 2 | 3 | 3 | 1 | 7 | 6 | 5 | 6 | 1 |
| | Ighv8-12 | 42 | 40 | 53 | 45 | 7 | 14 | 12 | 21 | 16 | 5 | 20 | 26 | 22 | 23 | 3 | 65 | 35 | 22 | 41 | 22 | 25 | 30 | 32 | 29 | 4 | 49 | 66 | 48 | 54 | 10 |
| | Ighv1-69 | 6 | 11 | 8 | 8 | 3 | 1 | 0 | 0 | 0 | 1 | 4 | 5 | 4 | 4 | 1 | 2 | 4 | 2 | 3 | 1 | 6 | 18 | 9 | 11 | 6 | 3 | 9 | 1 | 4 | 4 |
| **J558/3609** | Ighv1-67 | 58 | 57 | 58 | 58 | 1 | 10 | 15 | 9 | 11 | 3 | 24 | 26 | 16 | 22 | 5 | 64 | 32 | 41 | 46 | 17 | 41 | 25 | 36 | 34 | 8 | 38 | 81 | 52 | 57 | 22 |
| | Ighv1-66 | 7 | 6 | 1 | 5 | 3 | 0 | 0 | 0 | 0 | 0 | 6 | 0 | 2 | 3 | 3 | 3 | 4 | 0 | 2 | 2 | 2 | 2 | 2 | 2 | 0 | 2 | 2 | 3 | 2 | 1 |
| | Ighv8-11 | 19 | 23 | 34 | 25 | 8 | 7 | 1 | 5 | 4 | 3 | 12 | 12 | 11 | 12 | 1 | 67 | 19 | 14 | 33 | 29 | 13 | 22 | 14 | 16 | 5 | 20 | 31 | 36 | 29 | 8 |
| | Ighv1-64 | 28 | 39 | 30 | 32 | 6 | 8 | 5 | 5 | 6 | 2 | 15 | 19 | 21 | 18 | 3 | 75 | 20 | 23 | 39 | 31 | 35 | 38 | 40 | 38 | 3 | 46 | 74 | 22 | 47 | 26 |
| | Ighv1-63 | 4 | 20 | 9 | 11 | 8 | 2 | 4 | 2 | 3 | 1 | 2 | 4 | 3 | 3 | 1 | 16 | 10 | 5 | 10 | 6 | 4 | 7 | 3 | 5 | 2 | 7 | 23 | 8 | 13 | 9 |
| | Ighv1-62-3 | 7 | 9 | 12 | 9 | 3 | 2 | 3 | 4 | 3 | 1 | 7 | 4 | 4 | 5 | 2 | 10 | 1 | 5 | 5 | 5 | 6 | 6 | 4 | 5 | 1 | 10 | 8 | 7 | 8 | 2 |
| | Ighv1-62P | 0 | 0 | 0 | 0 | 0 | 0 | 0 | 0 | 0 | 0 | 0 | 0 | 0 | 0 | 0 | 0 | 0 | 0 | 0 | 0 | 0 | 0 | 0 | 0 | 0 | 0 | 0 | 0 | 0 | 0 |
| | Ighv1-61 | 10 | 7 | 3 | 7 | 4 | 0 | 1 | 0 | 0 | 1 | 5 | 4 | 1 | 3 | 2 | 7 | 2 | 2 | 4 | 3 | 3 | 6 | 8 | 6 | 3 | 3 | 10 | 4 | 6 | 4 |
| | Ighv1-59 | 9 | 8 | 8 | 8 | 1 | 0 | 2 | 4 | 2 | 2 | 5 | 1 | 2 | 3 | 2 | 5 | 5 | 2 | 4 | 2 | 2 | 8 | 10 | 7 | 4 | 6 | 11 | 4 | 7 | 4 |
| | Ighv1-58 | 5 | 0 | 2 | 2 | 3 | 0 | 0 | 0 | 0 | 0 | 0 | 1 | 2 | 1 | 1 | 0 | 0 | 0 | 0 | 0 | 2 | 0 | 1 | 1 | 1 | 2 | 2 | 5 | 3 | 2 |
| | Ighv8-8 | 2 | 3 | 2 | 2 | 1 | 1 | 0 | 0 | 0 | 1 | 2 | 1 | 2 | 2 | 1 | 0 | 0 | 0 | 0 | 0 | 0 | 1 | 2 | 1 | 1 | 1 | 1 | 1 | 1 | 0 |
| | Ighv8-7P | 6 | 6 | 7 | 6 | 1 | 2 | 1 | 0 | 1 | 1 | 1 | 1 | 1 | 1 | 0 | 11 | 1 | 2 | 5 | 6 | 4 | 2 | 4 | 3 | 1 | 7 | 4 | 6 | 6 | 2 |
| | Ighv1-56 | 4 | 9 | 4 | 6 | 3 | 0 | 0 | 0 | 0 | 0 | 2 | 1 | 3 | 2 | 1 | 7 | 2 | 2 | 4 | 3 | 2 | 3 | 1 | 2 | 1 | 2 | 3 | 4 | 3 | 1 |
| | Ighv1-55 | 26 | 20 | 20 | 22 | 3 | 2 | 7 | 5 | 5 | 3 | 10 | 11 | 6 | 9 | 3 | 40 | 13 | 6 | 20 | 18 | 15 | 19 | 15 | 16 | 2 | 22 | 37 | 21 | 27 | 9 |
| | Ighv1-54 | 35 | 49 | 37 | 40 | 8 | 3 | 9 | 7 | 6 | 3 | 17 | 23 | 23 | 21 | 3 | 42 | 18 | 22 | 27 | 13 | 34 | 33 | 28 | 32 | 3 | 47 | 49 | 34 | 43 | 8 |
| | Ighv1-53 | 11 | 13 | 19 | 14 | 4 | 1 | 5 | 4 | 3 | 2 | 6 | 8 | 2 | 5 | 3 | 5 | 2 | 2 | 3 | 2 | 13 | 14 | 6 | 11 | 4 | 9 | 2 | 2 | 4 | 4 |
| | Ighv1-52 | 4 | 8 | 4 | 5 | 2 | 0 | 1 | 0 | 0 | 1 | 5 | 3 | 3 | 4 | 1 | 5 | 7 | 4 | 5 | 2 | 4 | 5 | 3 | 4 | 1 | 6 | 8 | 9 | 8 | 2 |
| | Ighv1-50 | 3 | 3 | 2 | 3 | 1 | 0 | 1 | 0 | 0 | 1 | 0 | 3 | 1 | 1 | 2 | 5 | 2 | 2 | 3 | 2 | 0 | 2 | 7 | 3 | 4 | 5 | 4 | 3 | 4 | 1 |
| | Ighv8-5 | 1 | 0 | 0 | 0 | 1 | 0 | 0 | 0 | 0 | 0 | 1 | 0 | 2 | 1 | 1 | 2 | 0 | 0 | 1 | 1 | 0 | 2 | 2 | 1 | 1 | 1 | 1 | 0 | 1 | 1 |
| | Ighv1-49 | 0 | 0 | 0 | 0 | 0 | 0 | 0 | 0 | 0 | 0 | 0 | 0 | 0 | 0 | 0 | 0 | 0 | 0 | 0 | 0 | 0 | 0 | 0 | 0 | 0 | 0 | 0 | 0 | 0 | 0 |
| | Ighv8-4 | 7 | 6 | 8 | 7 | 1 | 0 | 1 | 1 | 1 | 1 | 1 | 2 | 4 | 2 | 2 | 3 | 8 | 5 | 5 | 3 | 2 | 7 | 5 | 5 | 3 | 5 | 0 | 2 | 2 | 3 |
| | Ighv1-47 | 44 | 18 | 19 | 27 | 15 | 2 | 1 | 1 | 1 | 1 | 3 | 7 | 11 | 7 | 4 | 21 | 16 | 20 | 19 | 3 | 11 | 11 | 27 | 16 | 9 | 27 | 12 | 9 | 16 | 10 |
| | Ighv1-43 | 8 | 11 | 6 | 8 | 3 | 2 | 0 | 0 | 1 | 1 | 5 | 5 | 2 | 4 | 2 | 9 | 7 | 2 | 6 | 4 | 6 | 7 | 8 | 7 | 1 | 2 | 12 | 4 | 6 | 5 |
| | Ighv1-42 | 22 | 19 | 19 | 20 | 2 | 5 | 4 | 9 | 6 | 3 | 7 | 15 | 9 | 10 | 4 | 29 | 25 | 16 | 23 | 7 | 15 | 26 | 29 | 23 | 7 | 20 | 35 | 18 | 24 | 9 |
| | Ighv1-40P | 3 | 1 | 0 | 1 | 2 | 0 | 1 | 0 | 0 | 1 | 2 | 1 | 1 | 1 | 1 | 1 | 0 | 1 | 1 | 1 | 0 | 3 | 0 | 1 | 2 | 3 | 5 | 0 | 3 | 3 |
| | Ighv1-39 | 13 | 13 | 11 | 12 | 1 | 4 | 8 | 4 | 5 | 2 | 8 | 8 | 6 | 7 | 1 | 17 | 18 | 7 | 14 | 6 | 13 | 10 | 10 | 11 | 2 | 22 | 20 | 6 | 16 | 9 |
| | Ighv1-38P | 0 | 2 | 0 | 1 | 1 | 0 | 0 | 0 | 0 | 0 | 2 | 1 | 0 | 1 | 1 | 0 | 0 | 0 | 0 | 0 | 1 | 0 | 0 | 0 | 1 | 0 | 0 | 0 | 0 | 0 |
| | Ighv1-37 | 5 | 2 | 3 | 3 | 2 | 1 | 1 | 1 | 1 | 0 | 2 | 4 | 2 | 3 | 1 | 4 | 1 | 2 | 2 | 2 | 4 | 5 | 2 | 4 | 2 | 5 | 4 | 1 | 3 | 2 |
| | Ighv1-36 | 0 | 0 | 3 | 1 | 2 | 0 | 0 | 0 | 0 | 0 | 1 | 0 | 0 | 0 | 1 | 0 | 1 | 1 | 1 | 1 | 1 | 1 | 0 | 1 | 1 | 2 | 1 | 0 | 1 | 1 |
| | Ighv1-34 | 13 | 10 | 21 | 15 | 6 | 0 | 4 | 0 | 1 | 2 | 7 | 7 | 4 | 6 | 2 | 23 | 3 | 5 | 10 | 11 | 13 | 20 | 13 | 15 | 4 | 9 | 20 | 2 | 10 | 9 |
| | Ighv1-33P | 23 | 20 | 13 | 19 | 5 | 4 | 8 | 3 | 5 | 3 | 10 | 3 | 7 | 7 | 4 | 29 | 15 | 4 | 16 | 13 | 13 | 18 | 17 | 16 | 3 | 27 | 24 | 12 | 21 | 8 |
| | Ighv1-32P | 3 | 5 | 1 | 3 | 2 | 1 | 1 | 0 | 1 | 1 | 0 | 2 | 4 | 2 | 2 | 2 | 3 | 0 | 2 | 2 | 4 | 3 | 3 | 3 | 1 | 4 | 4 | 3 | 4 | 1 |
| | Ighv1-31 | 4 | 1 | 0 | 2 | 2 | 2 | 1 | 0 | 1 | 1 | 2 | 2 | 2 | 2 | 0 | 4 | 7 | 1 | 4 | 3 | 3 | 6 | 1 | 3 | 3 | 4 | 4 | 2 | 3 | 1 |
| | Ighv1-30P | 0 | 0 | 0 | 0 | 0 | 0 | 0 | 0 | 0 | 0 | 2 | 0 | 0 | 1 | 1 | 2 | 0 | 3 | 2 | 2 | 1 | 2 | 2 | 2 | 1 | 0 | 0 | 1 | 0 | 1 |
| | Ighv1-28P | 4 | 1 | 1 | 2 | 2 | 0 | 0 | 0 | 0 | 0 | 2 | 0 | 0 | 1 | 1 | 0 | 2 | 1 | 1 | 1 | 1 | 1 | 6 | 3 | 3 | 2 | 5 | 3 | 3 | 2 |
| | Ighv1-27P | 5 | 13 | 9 | 9 | 4 | 2 | 5 | 0 | 2 | 3 | 2 | 1 | 1 | 1 | 1 | 15 | 7 | 12 | 11 | 4 | 8 | 12 | 8 | 9 | 2 | 14 | 22 | 7 | 14 | 8 |
| | Ighv1-26 | 34 | 27 | 25 | 29 | 5 | 5 | 3 | 3 | 4 | 1 | 8 | 10 | 12 | 10 | 2 | 45 | 21 | 14 | 27 | 16 | 19 | 23 | 19 | 20 | 2 | 31 | 40 | 19 | 30 | 11 |
| | Ighv1-25P | 2 | 3 | 1 | 2 | 1 | 0 | 1 | 0 | 0 | 1 | 0 | 0 | 2 | 1 | 1 | 3 | 2 | 2 | 2 | 1 | 0 | 1 | 3 | 1 | 2 | 2 | 0 | 2 | 1 | 1 |
| | Ighv1-23 | 2 | 5 | 1 | 3 | 2 | 0 | 2 | 1 | 1 | 1 | 4 | 1 | 0 | 2 | 2 | 2 | 6 | 0 | 3 | 3 | 8 | 9 | 1 | 6 | 4 | 2 | 12 | 3 | 6 | 6 |
| | Ighv1-22 | 3 | 4 | 1 | 3 | 2 | 1 | 1 | 0 | 1 | 1 | 0 | 1 | 1 | 1 | 1 | 2 | 10 | 1 | 4 | 5 | 0 | 1 | 1 | 1 | 1 | 5 | 3 | 3 | 4 | 1 |
| | Ighv1-21P | 1 | 1 | 2 | 1 | 1 | 0 | 2 | 0 | 1 | 1 | 1 | 0 | 0 | 0 | 1 | 3 | 1 | 0 | 1 | 2 | 3 | 2 | 1 | 2 | 1 | 1 | 1 | 0 | 1 | 1 |
| **J558** | Ighv1-21-1P | 2 | 9 | 5 | 5 | 4 | 0 | 0 | 0 | 0 | 0 | 2 | 0 | 2 | 1 | 1 | 8 | 4 | 1 | 4 | 4 | 8 | 3 | 3 | 5 | 3 | 1 | 3 | 0 | 1 | 2 |
| | Ighv1-20 | 8 | 15 | 15 | 13 | 4 | 3 | 1 | 4 | 3 | 2 | 4 | 8 | 3 | 5 | 3 | 14 | 9 | 4 | 9 | 5 | 18 | 22 | 13 | 18 | 5 | 16 | 44 | 10 | 23 | 18 |
| | Ighv1-19 | 15 | 15 | 12 | 14 | 2 | 1 | 1 | 0 | 1 | 1 | 9 | 2 | 3 | 5 | 4 | 21 | 11 | 8 | 13 | 7 | 10 | 15 | 12 | 12 | 3 | 10 | 22 | 15 | 16 | 6 |
| | Ighv1-18 | 29 | 28 | 27 | 28 | 1 | 3 | 5 | 1 | 3 | 2 | 14 | 12 | 4 | 10 | 5 | 52 | 22 | 11 | 28 | 21 | 30 | 15 | 21 | 22 | 8 | 19 | 47 | 32 | 33 | 14 |
| | Ighv1-17P | 0 | 0 | 0 | 0 | 0 | 0 | 0 | 1 | 0 | 1 | 0 | 0 | 0 | 0 | 0 | 0 | 0 | 0 | 0 | 0 | 0 | 0 | 0 | 0 | 0 | 0 | 2 | 1 | 1 | 1 |
| | Ighv1-16 | 0 | 0 | 0 | 0 | 0 | 0 | 0 | 0 | 0 | 0 | 0 | 0 | 0 | 0 | 0 | 0 | 0 | 0 | 0 | 0 | 0 | 0 | 0 | 0 | 0 | 0 | 0 | 0 | 0 | 0 |
| | Ighv1-15 | 0 | 1 | 1 | 1 | 1 | 1 | 1 | 0 | 1 | 1 | 1 | 2 | 1 | 1 | 1 | 5 | 0 | 1 | 2 | 3 | 0 | 2 | 2 | 1 | 1 | 3 | 7 | 2 | 4 | 3 |
| | Ighv1-14 | 45 | 47 | 44 | 45 | 2 | 5 | 10 | 11 | 9 | 3 | 23 | 21 | 34 | 26 | 7 | 46 | 37 | 40 | 41 | 5 | 23 | 30 | 54 | 36 | 16 | 53 | 87 | 49 | 63 | 21 |
| | Ighv1-13P | 3 | 6 | 2 | 4 | 2 | 0 | 0 | 0 | 0 | 0 | 4 | 2 | 1 | 2 | 2 | 4 | 0 | 2 | 2 | 2 | 4 | 0 | 2 | 2 | 2 | 4 | 12 | 3 | 6 | 5 |
| | Ighv1-12 | 20 | 20 | 20 | 20 | 0 | 3 | 3 | 5 | 4 | 1 | 6 | 10 | 8 | 8 | 2 | 14 | 7 | 8 | 10 | 4 | 7 | 9 | 14 | 10 | 4 | 10 | 15 | 10 | 12 | 3 |
| | Ighv1-11 | 41 | 43 | 57 | 47 | 9 | 7 | 11 | 13 | 10 | 3 | 23 | 10 | 16 | 16 | 7 | 64 | 21 | 17 | 34 | 26 | 55 | 54 | 40 | 50 | 8 | 51 | 54 | 41 | 49 | 7 |
| | Ighv1-9 | 66 | 69 | 56 | 64 | 7 | 17 | 16 | 12 | 15 | 3 | 20 | 24 | 26 | 23 | 3 | 91 | 17 | 32 | 47 | 39 | 36 | 63 | 65 | 55 | 16 | 67 | 77 | 44 | 63 | 17 |
| | Ighv15-2 | 59 | 59 | 44 | 54 | 9 | 6 | 9 | 4 | 6 | 3 | 30 | 24 | 28 | 27 | 3 | 76 | 51 | 29 | 52 | 24 | 35 | 50 | 45 | 43 | 8 | 44 | 56 | 30 | 43 | 13 |
| | Ighv1-8P | 2 | 1 | 1 | 1 | 1 | 0 | 0 | 2 | 1 | 1 | 1 | 0 | 1 | 1 | 1 | 0 | 0 | 0 | 0 | 0 | 1 | 1 | 1 | 1 | 0 | 2 | 4 | 1 | 2 | 2 |
| | Ighv1-7 | 1 | 1 | 1 | 1 | 0 | 0 | 0 | 0 | 0 | 0 | 0 | 0 | 1 | 0 | 1 | 0 | 0 | 0 | 0 | 0 | 0 | 0 | 2 | 1 | 1 | 1 | 0 | 0 | 0 | 1 |
| | Ighv10-3 | 40 | 32 | 26 | 33 | 7 | 5 | 5 | 6 | 5 | 1 | 15 | 10 | 19 | 15 | 5 | 35 | 14 | 22 | 24 | 11 | 21 | 19 | 39 | 26 | 11 | 27 | 43 | 18 | 29 | 13 |
| | Ighv1-5 | 60 | 50 | 65 | 58 | 8 | 7 | 8 | 16 | 10 | 5 | 33 | 33 | 28 | 31 | 3 | 106 | 37 | 46 | 63 | 38 | 45 | 37 | 49 | 44 | 6 | 62 | 84 | 52 | 66 | 16 |
| | Ighv1-4 | 47 | 29 | 26 | 34 | 11 | 10 | 4 | 13 | 9 | 5 | 14 | 23 | 22 | 20 | 5 | 75 | 46 | 16 | 46 | 30 | 25 | 40 | 31 | 32 | 8 | 30 | 53 | 34 | 39 | 12 |
| | Ighv10-1 | 41 | 33 | 41 | 38 | 5 | 10 | 6 | 5 | 7 | 3 | 10 | 18 | 16 | 15 | 4 | 26 | 25 | 11 | 21 | 8 | 25 | 24 | 31 | 27 | 4 | 33 | 31 | 5 | 23 | 16 |
| | Ighv1-2P | 75 | 54 | 57 | 62 | 11 | 8 | 15 | 14 | 12 | 4 | 29 | 32 | 28 | 30 | 2 | 84 | 43 | 45 | 57 | 23 | 47 | 56 | 41 | 48 | 8 | 69 | 86 | 49 | 68 | 19 |
| | Ighv8-2 | 56 | 53 | 38 | 49 | 10 | 6 | 4 | 9 | 6 | 3 | 24 | 18 | 19 | 20 | 3 | 49 | 32 | 25 | 35 | 12 | 43 | 53 | 48 | 48 | 5 | 30 | 65 | 35 | 43 | 19 |
| | IGHV6-7 | 31 | 25 | 25 | 27 | 3 | 2 | 1 | 4 | 2 | 2 | 13 | 10 | 7 | 10 | 3 | 23 | 20 | 12 | 18 | 6 | 29 | 18 | 19 | 22 | 6 | 18 | 20 | 15 | 18 | 3 |
| | IGHV6-6 | 63 | 41 | 54 | 53 | 11 | 21 | 9 | 12 | 14 | 6 | 43 | 24 | 23 | 30 | 11 | 87 | 44 | 42 | 58 | 25 | 47 | 39 | 49 | 45 | 5 | 69 | 76 | 40 | 62 | 19 |
| | IGHV6-5 | 1 | 1 | 0 | 1 | 1 | 0 | 0 | 0 | 0 | 0 | 0 | 1 | 0 | 0 | 1 | 2 | 1 | 0 | 1 | 1 | 1 | 1 | 0 | 1 | 1 | 2 | 2 | 0 | 1 | 1 |
| | IGHV6-4 | 6 | 1 | 1 | 3 | 3 | 0 | 0 | 0 | 0 | 0 | 0 | 0 | 0 | 0 | 0 | 3 | 1 | 0 | 1 | 2 | 4 | 2 | 4 | 3 | 1 | 4 | 5 | 1 | 3 | 2 |
| | IGHV6-3 | 4 | 6 | 5 | 5 | 1 | 2 | 2 | 0 | 1 | 1 | 2 | 3 | 2 | 2 | 1 | 5 | 0 | 4 | 3 | 3 | 2 | 5 | 4 | 4 | 2 | 1 | 4 | 0 | 2 | 2 |
| | IGHV12-3 | 30 | 18 | 32 | 27 | 8 | 1 | 5 | 4 | 3 | 2 | 13 | 11 | 7 | 10 | 3 | 14 | 16 | 17 | 16 | 2 | 13 | 18 | 16 | 16 | 3 | 21 | 22 | 13 | 19 | 5 |
| | IGHV13-2 | 19 | 22 | 23 | 21 | 2 | 2 | 4 | 11 | 6 | 5 | 10 | 6 | 12 | 9 | 3 | 27 | 23 | 16 | 22 | 6 | 17 | 18 | 30 | 22 | 7 | 9 | 19 | 18 | 15 | 6 |
| | IGHV3-7P | 0 | 0 | 0 | 0 | 0 | 0 | 0 | 0 | 0 | 0 | 0 | 0 | 0 | 0 | 0 | 0 | 0 | 1 | 0 | 1 | 0 | 0 | 0 | 0 | 0 | 0 | 0 | 0 | 0 | 0 |
| | IGHV9-4 | 23 | 21 | 23 | 22 | 1 | 5 | 1 | 12 | 6 | 6 | 5 | 10 | 11 | 9 | 3 | 36 | 14 | 23 | 24 | 11 | 20 | 33 | 33 | 29 | 8 | 21 | 20 | 13 | 18 | 4 |
| | IGHV3-6 | 47 | 51 | 54 | 51 | 4 | 7 | 10 | 7 | 8 | 2 | 15 | 27 | 21 | 21 | 6 | 73 | 50 | 23 | 49 | 25 | 41 | 63 | 38 | 47 | 14 | 32 | 45 | 40 | 39 | 7 |
| | IGHV13-1P | 0 | 1 | 0 | 0 | 1 | 0 | 0 | 1 | 0 | 1 | 0 | 1 | 0 | 0 | 1 | 0 | 0 | 0 | 0 | 0 | 0 | 2 | 0 | 1 | 1 | 0 | 0 | 1 | 0 | 1 |
| | IGHV3-5 | 15 | 12 | 4 | 10 | 6 | 4 | 1 | 3 | 3 | 2 | 9 | 4 | 3 | 5 | 3 | 14 | 22 | 9 | 15 | 7 | 15 | 25 | 8 | 16 | 9 | 14 | 6 | 7 | 9 | 4 |
| | IGHV3-4 | 5 | 4 | 3 | 4 | 1 | 1 | 0 | 0 | 0 | 1 | 1 | 3 | 0 | 1 | 2 | 1 | 2 | 1 | 1 | 1 | 6 | 7 | 2 | 5 | 3 | 1 | 0 | 3 | 1 | 2 |
| | IGHV7-4 | 6 | 5 | 1 | 4 | 3 | 1 | 2 | 0 | 1 | 1 | 3 | 0 | 0 | 1 | 2 | 5 | 1 | 2 | 3 | 2 | 2 | 2 | 6 | 3 | 2 | 6 | 2 | 3 | 4 | 2 |
| | IGHV3-3P | 3 | 7 | 4 | 5 | 2 | 0 | 1 | 2 | 1 | 1 | 1 | 0 | 1 | 1 | 1 | 0 | 4 | 1 | 2 | 2 | 8 | 2 | 3 | 4 | 3 | 0 | 4 | 2 | 2 | 2 |
| | IGHV14-4 | 24 | 41 | 39 | 35 | 9 | 8 | 12 | 8 | 9 | 2 | 18 | 17 | 19 | 18 | 1 | 24 | 29 | 10 | 21 | 10 | 34 | 47 | 33 | 38 | 8 | 34 | 62 | 45 | 47 | 14 |
| **Middle** | IGHV7-3 | 34 | 46 | 43 | 41 | 6 | 11 | 11 | 8 | 10 | 2 | 28 | 23 | 26 | 26 | 3 | 67 | 25 | 24 | 39 | 25 | 38 | 30 | 37 | 35 | 4 | 37 | 47 | 20 | 35 | 14 |
| | IGHV9-2-1 | 48 | 65 | 38 | 50 | 14 | 13 | 10 | 8 | 10 | 3 | 23 | 21 | 20 | 21 | 2 | 64 | 31 | 40 | 45 | 17 | 56 | 56 | 43 | 52 | 8 | 55 | 58 | 32 | 48 | 14 |
| | IGHV12-2-1 | 0 | 0 | 0 | 0 | 0 | 0 | 0 | 0 | 0 | 0 | 0 | 0 | 1 | 0 | 1 | 1 | 0 | 0 | 0 | 1 | 2 | 0 | 0 | 1 | 1 | 0 | 0 | 0 | 0 | 0 |
| | IGHV9-3-1 | 31 | 28 | 29 | 29 | 2 | 6 | 6 | 5 | 6 | 1 | 20 | 24 | 19 | 21 | 3 | 30 | 23 | 14 | 22 | 8 | 20 | 22 | 31 | 24 | 6 | 25 | 45 | 14 | 28 | 16 |
| | IGHV12-1-1 | 20 | 20 | 19 | 20 | 1 | 3 | 5 | 1 | 3 | 2 | 13 | 15 | 5 | 11 | 5 | 17 | 14 | 7 | 13 | 5 | 13 | 16 | 11 | 13 | 3 | 19 | 16 | 8 | 14 | 6 |
| | IGHV9-3 | 17 | 8 | 13 | 13 | 5 | 6 | 5 | 4 | 5 | 1 | 10 | 3 | 9 | 7 | 4 | 15 | 4 | 12 | 10 | 6 | 14 | 17 | 20 | 17 | 3 | 6 | 22 | 9 | 12 | 9 |
| | IGHV9-2 | 12 | 7 | 12 | 10 | 3 | 0 | 0 | 0 | 0 | 0 | 2 | 9 | 2 | 4 | 4 | 6 | 12 | 11 | 10 | 3 | 12 | 7 | 8 | 9 | 3 | 12 | 9 | 5 | 9 | 4 |
| | IGHV9-1 | 11 | 16 | 20 | 16 | 5 | 4 | 2 | 4 | 3 | 1 | 5 | 4 | 4 | 4 | 1 | 20 | 11 | 4 | 12 | 8 | 12 | 15 | 12 | 13 | 2 | 15 | 21 | 10 | 15 | 6 |
| | IGHV14-3 | 38 | 68 | 48 | 51 | 15 | 8 | 26 | 10 | 15 | 10 | 38 | 41 | 19 | 33 | 12 | 39 | 45 | 13 | 32 | 17 | 45 | 88 | 62 | 65 | 22 | 51 | 61 | 18 | 43 | 23 |

| | | | | | | | | | | | | | | | | | | | | | | | | | | | | | | | |
|---|---|---|---|---|---|---|---|---|---|---|---|---|---|---|---|---|---|---|---|---|---|---|---|---|---|---|---|---|---|---|---|
| | IGHV11-2 | 36 | 31 | 29 | 32 | 4 | 6 | 2 | 15 | 8 | 7 | 13 | 23 | 26 | 21 | 7 | 28 | 36 | 18 | 27 | 9 | 28 | 24 | 15 | 22 | 7 | 25 | 30 | 31 | 29 | 3 |
| | IGHV3-2 | 60 | 72 | 77 | 70 | 9 | 11 | 30 | 17 | 19 | 10 | 37 | 48 | 33 | 39 | 8 | 62 | 12 | 22 | 32 | 26 | 59 | 84 | 84 | 76 | 14 | 42 | 40 | 21 | 34 | 12 |
| | IGHV4-2 | 21 | 19 | 24 | 21 | 3 | 14 | 3 | 4 | 7 | 6 | 13 | 21 | 15 | 16 | 4 | 17 | 14 | 5 | 12 | 6 | 13 | 29 | 28 | 23 | 9 | 18 | 19 | 5 | 14 | 8 |
| | IGHV14-2P | 13 | 15 | 18 | 15 | 3 | 3 | 1 | 2 | 2 | 1 | 11 | 6 | 4 | 7 | 4 | 12 | 7 | 7 | 9 | 3 | 12 | 12 | 10 | 11 | 1 | 17 | 16 | 5 | 13 | 7 |
| | IGHV11-1 | 11 | 8 | 16 | 12 | 4 | 3 | 2 | 0 | 2 | 2 | 3 | 6 | 5 | 5 | 2 | 8 | 18 | 6 | 11 | 6 | 12 | 16 | 12 | 13 | 2 | 16 | 26 | 11 | 18 | 8 |
| | IGHV3-1 | 29 | 19 | 21 | 23 | 5 | 4 | 9 | 4 | 6 | 3 | 19 | 12 | 18 | 16 | 4 | 24 | 11 | 16 | 17 | 7 | 22 | 25 | 21 | 23 | 2 | 14 | 25 | 7 | 15 | 9 |
| | IGHV4-1 | 22 | 33 | 51 | 35 | 15 | 9 | 10 | 9 | 9 | 1 | 28 | 27 | 26 | 27 | 1 | 39 | 27 | 10 | 25 | 15 | 40 | 38 | 38 | 39 | 1 | 34 | 26 | 13 | 24 | 11 |
| | IGHV14-1 | 15 | 28 | 16 | 20 | 7 | 3 | 4 | 5 | 4 | 1 | 16 | 10 | 11 | 12 | 3 | 19 | 23 | 16 | 19 | 4 | 16 | 20 | 19 | 18 | 2 | 20 | 22 | 19 | 20 | 2 |
| | IGHV7-1 | 61 | 67 | 54 | 61 | 7 | 27 | 16 | 13 | 19 | 7 | 34 | 42 | 53 | 43 | 10 | 100 | 53 | 33 | 62 | 34 | 52 | 64 | 45 | 54 | 10 | 55 | 68 | 37 | 53 | 16 |
| | IGHV2-9 | 68 | 65 | 97 | 77 | 18 | 19 | 34 | 26 | 26 | 8 | 37 | 69 | 72 | 59 | 19 | 95 | 55 | 59 | 70 | 22 | 67 | 108 | 91 | 89 | 21 | 55 | 95 | 45 | 65 | 26 |
| | IGHV5-17 | 53 | 38 | 43 | 45 | 8 | 10 | 13 | 9 | 11 | 2 | 28 | 29 | 32 | 30 | 2 | 57 | 40 | 27 | 41 | 15 | 40 | 46 | 47 | 44 | 4 | 36 | 70 | 37 | 48 | 19 |
| | IGHV5-15 | 18 | 13 | 7 | 13 | 6 | 8 | 4 | 3 | 5 | 3 | 6 | 4 | 7 | 6 | 2 | 17 | 26 | 15 | 19 | 6 | 12 | 21 | 18 | 17 | 5 | 25 | 25 | 22 | 24 | 2 |
| | IGHV2-7P (VHQ52.a2! | 6 | 4 | 6 | 5 | 1 | 0 | 0 | 0 | 0 | 0 | 5 | 5 | 13 | 8 | 5 | 4 | 2 | 4 | 3 | 1 | 10 | 2 | 6 | 6 | 4 | 7 | 5 | 2 | 5 | 3 |
| | IGHV2-6-7 | 63 | 62 | 63 | 63 | 1 | 22 | 14 | 14 | 17 | 5 | 29 | 42 | 37 | 36 | 7 | 65 | 50 | 43 | 53 | 11 | 64 | 59 | 59 | 61 | 3 | 50 | 44 | 49 | 48 | 3 |
| | IGHV2-6-6 | 1 | 4 | 5 | 3 | 2 | 1 | 1 | 1 | 1 | 0 | 1 | 1 | 2 | 1 | 1 | 7 | 0 | 1 | 3 | 4 | 0 | 1 | 1 | 1 | 1 | 1 | 0 | 0 | 0 | 1 |
| | IGHV2-6-5 | 13 | 7 | 8 | 9 | 3 | 0 | 4 | 1 | 2 | 2 | 2 | 7 | 7 | 5 | 3 | 5 | 4 | 8 | 6 | 2 | 16 | 11 | 7 | 11 | 5 | 3 | 7 | 8 | 6 | 3 |
| | IGHV2-2-2 | 9 | 9 | 22 | 13 | 8 | 1 | 5 | 2 | 3 | 2 | 5 | 11 | 7 | 8 | 3 | 24 | 11 | 11 | 15 | 8 | 8 | 14 | 14 | 12 | 3 | 21 | 17 | 7 | 15 | 7 |
| | IGHV2-6-4 | 17 | 25 | 28 | 23 | 6 | 1 | 7 | 9 | 6 | 4 | 10 | 6 | 17 | 11 | 6 | 26 | 10 | 3 | 13 | 12 | 12 | 12 | 8 | 11 | 2 | 8 | 14 | 10 | 11 | 3 |
| | IGHV5-6-5 | 26 | 28 | 31 | 28 | 3 | 7 | 3 | 5 | 5 | 2 | 17 | 11 | 10 | 13 | 4 | 13 | 19 | 13 | 15 | 3 | 19 | 23 | 30 | 24 | 6 | 14 | 23 | 13 | 17 | 6 |
| | IGHV5-12-2 | 20 | 24 | 16 | 20 | 4 | 11 | 3 | 9 | 8 | 4 | 15 | 14 | 15 | 15 | 1 | 51 | 37 | 16 | 35 | 18 | 14 | 20 | 14 | 16 | 3 | 26 | 32 | 21 | 26 | 6 |
| | IGHV5-6-4 | 21 | 14 | 18 | 18 | 4 | 7 | 0 | 7 | 5 | 4 | 10 | 14 | 8 | 11 | 3 | 41 | 14 | 29 | 28 | 14 | 21 | 22 | 24 | 22 | 2 | 36 | 26 | 15 | 26 | 11 |
| | IGHV2-6-3 | 9 | 6 | 10 | 8 | 2 | 0 | 3 | 0 | 1 | 2 | 4 | 5 | 0 | 3 | 3 | 2 | 3 | 9 | 5 | 4 | 3 | 7 | 15 | 8 | 6 | 7 | 15 | 8 | 10 | 4 |
| | IGHV5-6-3 | 6 | 17 | 26 | 16 | 10 | 6 | 6 | 5 | 6 | 1 | 4 | 7 | 2 | 4 | 3 | 19 | 9 | 4 | 11 | 8 | 11 | 18 | 7 | 12 | 6 | 8 | 11 | 5 | 8 | 3 |
| | IGHV5-12-1 | 16 | 16 | 22 | 18 | 3 | 2 | 3 | 3 | 3 | 1 | 8 | 4 | 8 | 7 | 2 | 8 | 16 | 7 | 10 | 5 | 10 | 12 | 22 | 15 | 6 | 11 | 9 | 11 | 10 | 1 |
| | IGHV5-9-5 | 1 | 8 | 3 | 4 | 4 | 1 | 0 | 1 | 1 | 1 | 1 | 0 | 4 | 2 | 2 | 7 | 2 | 0 | 3 | 4 | 3 | 2 | 1 | 2 | 1 | 1 | 5 | 4 | 3 | 2 |
| | IGHV2-6-2 | 9 | 2 | 4 | 5 | 4 | 0 | 0 | 0 | 0 | 0 | 1 | 0 | 6 | 2 | 3 | 1 | 1 | 2 | 1 | 1 | 6 | 2 | 2 | 3 | 2 | 4 | 3 | 3 | 3 | 1 |
| **7183/Q52** | IGHV5-6-2 | 16 | 19 | 25 | 20 | 5 | 3 | 3 | 4 | 3 | 1 | 7 | 13 | 15 | 12 | 4 | 6 | 17 | 22 | 15 | 8 | 11 | 11 | 15 | 12 | 2 | 17 | 27 | 24 | 23 | 5 |
| | IGHV2-9-2 | 12 | 15 | 20 | 16 | 4 | 1 | 4 | 1 | 2 | 2 | 11 | 5 | 5 | 7 | 3 | 20 | 10 | 10 | 13 | 6 | 7 | 14 | 8 | 10 | 4 | 12 | 15 | 12 | 13 | 2 |
| | IGHV5-9-4 | 24 | 24 | 34 | 27 | 6 | 8 | 9 | 4 | 7 | 3 | 9 | 5 | 10 | 8 | 3 | 25 | 29 | 26 | 27 | 2 | 13 | 13 | 20 | 15 | 4 | 29 | 16 | 21 | 22 | 7 |
| | IGHV2-4-1 | 10 | 12 | 5 | 9 | 4 | 3 | 1 | 0 | 1 | 2 | 0 | 11 | 3 | 5 | 6 | 6 | 4 | 6 | 5 | 1 | 6 | 7 | 10 | 8 | 2 | 11 | 8 | 6 | 8 | 3 |
| | IGHV5-9-3 | 32 | 34 | 23 | 30 | 6 | 4 | 15 | 7 | 9 | 6 | 6 | 10 | 10 | 9 | 2 | 42 | 29 | 12 | 28 | 15 | 23 | 23 | 17 | 21 | 3 | 33 | 41 | 26 | 33 | 8 |
| | IGHV2-2-1P | 20 | 11 | 12 | 14 | 5 | 3 | 0 | 0 | 1 | 2 | 1 | 7 | 8 | 5 | 4 | 14 | 15 | 3 | 11 | 7 | 10 | 8 | 15 | 11 | 4 | 3 | 19 | 15 | 12 | 8 |
| | IGHV5-9-2 | 2 | 6 | 6 | 5 | 2 | 7 | 0 | 0 | 2 | 4 | 8 | 4 | 6 | 6 | 2 | 9 | 11 | 5 | 8 | 3 | 2 | 4 | 8 | 5 | 3 | 4 | 10 | 7 | 7 | 3 |
| | IGHV2-6-1 | 9 | 12 | 12 | 11 | 2 | 2 | 1 | 0 | 1 | 1 | 3 | 6 | 4 | 4 | 2 | 7 | 4 | 3 | 5 | 2 | 5 | 6 | 9 | 7 | 2 | 5 | 6 | 2 | 4 | 2 |
| | IGHV5-9-1 | 35 | 41 | 52 | 43 | 9 | 11 | 11 | 12 | 11 | 1 | 28 | 38 | 16 | 27 | 11 | 57 | 84 | 41 | 61 | 22 | 39 | 63 | 45 | 49 | 12 | 54 | 72 | 47 | 58 | 13 |
| | IGHV2-6 | 11 | 7 | 9 | 9 | 2 | 1 | 2 | 2 | 2 | 1 | 2 | 8 | 0 | 3 | 4 | 5 | 6 | 12 | 8 | 4 | 12 | 11 | 7 | 10 | 3 | 4 | 12 | 4 | 7 | 5 |
| | IGHV5-12 | 44 | 71 | 51 | 55 | 14 | 8 | 13 | 20 | 14 | 6 | 19 | 14 | 15 | 16 | 3 | 94 | 104 | 111 | 103 | 9 | 28 | 47 | 35 | 37 | 10 | 65 | 82 | 58 | 68 | 12 |
| | IGHV5-9 | 4 | 13 | 7 | 8 | 5 | 1 | 2 | 1 | 1 | 1 | 6 | 11 | 5 | 7 | 3 | 29 | 16 | 21 | 22 | 7 | 6 | 3 | 7 | 5 | 2 | 10 | 15 | 8 | 11 | 4 |
| | IGHV2-4 | 5 | 7 | 8 | 7 | 2 | 4 | 7 | 1 | 4 | 3 | 9 | 11 | 9 | 10 | 1 | 3 | 10 | 12 | 8 | 5 | 4 | 9 | 1 | 5 | 4 | 3 | 10 | 5 | 6 | 4 |
| | **IGHV5-6** | 68 | 59 | 70 | 66 | 6 | 17 | 13 | 16 | 15 | 2 | 25 | 13 | 14 | 17 | 7 | 117 | 137 | 130 | 128 | 10 | 19 | 33 | 29 | 27 | 7 | 116 | 84 | 25 | 75 | 46 |
| | **IGHV2-3** | 81 | 104 | 87 | 91 | 12 | 32 | 46 | 40 | 39 | 7 | 41 | 44 | 32 | 39 | 6 | 164 | 173 | 148 | 162 | 13 | 33 | 31 | 46 | 37 | 8 | 117 | 112 | 54 | 94 | 35 |
| | **IGHV5-4** | 96 | 175 | 110 | 127 | 42 | 116 | 110 | 120 | 115 | 5 | 107 | 150 | 142 | 133 | 23 | 282 | 297 | 254 | 278 | 22 | 67 | 74 | 80 | 74 | 7 | 210 | 185 | 97 | 164 | 59 |
| | **IGHV2-2** | 189 | 256 | 226 | 224 | 34 | 619 | 633 | 585 | 612 | 25 | 632 | 721 | 701 | 685 | 47 | 570 | 571 | 479 | 540 | 53 | 338 | 366 | 386 | 363 | 24 | 368 | 340 | 207 | 305 | 86 |
| | **IGHV5-2 (VH81X)** | 322 | 400 | 403 | 375 | 46 | 5999 | 5293 | 5254 | 5515 | 419 | 3604 | 5012 | 4571 | 4396 | 720 | 1037 | 1085 | 896 | 1006 | 98 | 1559 | 1795 | 1795 | 1716 | 136 | 651 | 565 | 242 | 486 | 216 |
| | **IGHV5-1P** | 10 | 23 | 16 | 16 | 7 | 600 | 586 | 527 | 571 | 39 | 185 | 239 | 234 | 219 | 30 | 67 | 45 | 63 | 58 | 12 | 76 | 85 | 83 | 81 | 5 | 31 | 34 | 2 | 22 | 18 |
| | D usage in DJH joins | | | | | | | | | | | | | | | | | | | | | | | | | | | | | | |
| | **IGHD1-1 (DFL16.1)** | 1167 | 1483 | 1789 | 1480 | 311 | 22 | 26 | 29 | 26 | 4 | 513 | 612 | 579 | 568 | 50 | 534 | 790 | 613 | 646 | 131 | 826 | 985 | 892 | 901 | 80 | 584 | 533 | 301 | 473 | 151 |
| | **IGHD2-3** | 533 | 667 | 803 | 668 | 135 | 86 | 75 | 83 | 81 | 6 | 191 | 222 | 218 | 210 | 17 | 447 | 560 | 457 | 488 | 63 | 437 | 455 | 446 | 446 | 9 | 573 | 417 | 275 | 422 | 149 |
| | **IGHD2-2** | 274 | 342 | 440 | 352 | 83 | 92 | 85 | 104 | 94 | 10 | 126 | 151 | 183 | 153 | 29 | 268 | 297 | 257 | 274 | 21 | 241 | 347 | 358 | 315 | 65 | 283 | 235 | 153 | 224 | 66 |
| | **IGHD3-3** | 1 | 1 | 1 | 1 | 0 | 1 | 0 | 0 | 0 | 1 | 0 | 0 | 1 | 0 | 1 | 0 | 0 | 2 | 1 | 1 | 0 | 1 | 4 | 2 | 2 | 0 | 0 | 2 | 1 | 1 |
| | **IGHD1-2** | 329 | 475 | 580 | 461 | 126 | 69 | 42 | 69 | 60 | 16 | 181 | 190 | 147 | 173 | 23 | 314 | 467 | 351 | 377 | 80 | 413 | 487 | 347 | 416 | 70 | 560 | 419 | 268 | 416 | 146 |
| | **IGHD2-1** | 492 | 587 | 692 | 590 | 100 | 179 | 179 | 183 | 180 | 2 | 276 | 275 | 313 | 288 | 22 | 514 | 677 | 541 | 577 | 87 | 540 | 538 | 448 | 509 | 53 | 664 | 529 | 335 | 509 | 165 |
| | **IGHD2-4/2-9** | 781 | 797 | 1002 | 860 | 123 | 113 | 128 | 126 | 122 | 8 | 237 | 277 | 262 | 259 | 20 | 612 | 767 | 766 | 715 | 89 | 613 | 670 | 689 | 657 | 40 | 748 | 656 | 423 | 609 | 168 |
| | **IGHD2-14** | 595 | 586 | 763 | 648 | 100 | 65 | 59 | 82 | 69 | 12 | 157 | 211 | 167 | 178 | 29 | 424 | 604 | 518 | 515 | 90 | 420 | 551 | 542 | 504 | 73 | 577 | 409 | 345 | 444 | 120 |
| | **IGHD2-10** | 581 | 493 | 672 | 582 | 90 | 106 | 91 | 148 | 115 | 30 | 226 | 250 | 245 | 240 | 13 | 392 | 608 | 571 | 524 | 116 | 448 | 471 | 505 | 475 | 29 | 547 | 385 | 336 | 423 | 110 |
| | **IGHD2-11** | 396 | 414 | 464 | 425 | 35 | 99 | 104 | 97 | 100 | 4 | 130 | 145 | 151 | 142 | 11 | 328 | 365 | 366 | 353 | 22 | 295 | 331 | 316 | 314 | 18 | 375 | 307 | 276 | 319 | 51 |
| | **IGHD3-2** | 108 | 115 | 180 | 134 | 40 | 23 | 22 | 33 | 26 | 6 | 69 | 67 | 59 | 65 | 5 | 190 | 256 | 189 | 212 | 38 | 120 | 133 | 117 | 123 | 9 | 275 | 223 | 185 | 228 | 45 |
| | **IGHD4-1 (DQ52)** | 323 | 345 | 444 | 371 | 64 | 157 | 178 | 238 | 191 | 42 | 207 | 244 | 249 | 233 | 23 | 365 | 372 | 414 | 384 | 27 | 320 | 359 | 316 | 332 | 24 | 404 | 400 | 303 | 369 | 57 |
| | D usage in VHDJH joins | | | | | | | | | | | | | | | | | | | | | | | | | | | | | | |
| | **IGHD1-1 (DFL16.1)** | 500 | 561 | 576 | 546 | 40 | 237 | 220 | 183 | 213 | 28 | 669 | 893 | 825 | 795 | 115 | 413 | 309 | 315 | 346 | 59 | 488 | 524 | 574 | 528 | 43 | 300 | 329 | 211 | 280 | 62 |
| | **IGHD2-3** | 265 | 286 | 275 | 276 | 11 | 501 | 448 | 405 | 452 | 48 | 491 | 596 | 573 | 553 | 56 | 388 | 337 | 237 | 320 | 77 | 372 | 355 | 394 | 374 | 20 | 275 | 403 | 181 | 286 | 112 |
| | **IGHD2-2** | 149 | 118 | 116 | 128 | 19 | 303 | 318 | 228 | 283 | 48 | 259 | 281 | 279 | 273 | 12 | 248 | 152 | 118 | 173 | 67 | 164 | 191 | 173 | 176 | 14 | 163 | 207 | 87 | 152 | 61 |
| | **IGHD3-3** | 79 | 79 | 64 | 74 | 8 | 193 | 218 | 215 | 209 | 14 | 121 | 222 | 236 | 193 | 63 | 149 | 124 | 116 | 129 | 17 | 101 | 120 | 120 | 114 | 11 | 106 | 95 | 35 | 79 | 38 |
| | **IGHD1-2** | 236 | 284 | 260 | 260 | 24 | 491 | 464 | 448 | 468 | 22 | 346 | 460 | 410 | 405 | 58 | 392 | 298 | 237 | 309 | 78 | 286 | 359 | 362 | 336 | 43 | 286 | 447 | 190 | 308 | 130 |
| | **IGHD2-1** | 157 | 137 | 149 | 148 | 10 | 544 | 389 | 420 | 451 | 82 | 313 | 366 | 288 | 323 | 40 | 216 | 185 | 199 | 200 | 15 | 170 | 221 | 287 | 226 | 59 | 174 | 206 | 160 | 180 | 24 |
| | **IGHD2-4/2-9** | 404 | 446 | 433 | 428 | 22 | 1078 | 935 | 1044 | 1019 | 75 | 720 | 954 | 891 | 855 | 121 | 814 | 625 | 492 | 644 | 162 | 518 | 646 | 511 | 558 | 76 | 534 | 595 | 328 | 485 | 140 |
| | **IGHD2-14** | 432 | 580 | 544 | 519 | 77 | 1317 | 1232 | 1161 | 1237 | 78 | 929 | 1105 | 1106 | 1046 | 102 | 823 | 693 | 542 | 686 | 140 | 602 | 740 | 763 | 702 | 87 | 730 | 677 | 400 | 602 | 178 |
| | **IGHD2-10** | 292 | 362 | 277 | 310 | 45 | 1185 | 1176 | 1041 | 1134 | 80 | 735 | 946 | 787 | 823 | 110 | 557 | 499 | 433 | 496 | 62 | 395 | 508 | 537 | 480 | 75 | 382 | 427 | 233 | 347 | 102 |
| | **IGHD2-11** | 175 | 158 | 165 | 166 | 9 | 617 | 587 | 576 | 594 | 21 | 317 | 570 | 486 | 458 | 129 | 390 | 356 | 242 | 329 | 78 | 306 | 300 | 207 | 271 | 55 | 242 | 200 | 136 | 193 | 53 |
| | **IGHD3-2** | 110 | 115 | 145 | 123 | 19 | 353 | 321 | 378 | 351 | 29 | 262 | 297 | 337 | 299 | 37 | 216 | 158 | 140 | 171 | 40 | 138 | 178 | 186 | 167 | 26 | 122 | 220 | 89 | 144 | 68 |
| | **IGHD4-1 (DQ52)** | 172 | 172 | 169 | 171 | 2 | 621 | 546 | 541 | 570 | 45 | 360 | 410 | 347 | 372 | 33 | 369 | 267 | 219 | 285 | 77 | 237 | 261 | 227 | 241 | 18 | 193 | 300 | 192 | 228 | 62 |
| | D usage in DJH+VHDJH joins | | | | | | | | | | | | | | | | | | | | | | | | | | | | | | |
| | **IGHD1-1 (DFL16.1)** | 1667 | 2044 | 2365 | 2025 | 349 | 259 | 246 | 212 | 239 | 24 | 1182 | 1505 | 1404 | 1363 | 165 | 947 | 1099 | 928 | 991 | 94 | 1314 | 1509 | 1466 | 1429 | 103 | 884 | 862 | 512 | 753 | 209 |
| | **IGHD2-3** | 798 | 953 | 1078 | 943 | 140 | 587 | 523 | 488 | 533 | 50 | 682 | 818 | 791 | 764 | 72 | 835 | 897 | 694 | 808 | 104 | 809 | 810 | 840 | 820 | 18 | 848 | 820 | 456 | 708 | 219 |
| | **IGHD2-2** | 423 | 460 | 556 | 480 | 69 | 395 | 403 | 332 | 377 | 39 | 385 | 432 | 462 | 426 | 39 | 516 | 449 | 375 | 447 | 70 | 405 | 538 | 531 | 491 | 75 | 446 | 442 | 240 | 376 | 118 |
| | **IGHD3-3** | 80 | 80 | 65 | 75 | 8 | 194 | 218 | 215 | 209 | 13 | 121 | 222 | 237 | 193 | 63 | 149 | 124 | 118 | 130 | 16 | 101 | 121 | 124 | 115 | 12 | 106 | 95 | 37 | 79 | 37 |
| | **IGHD1-2** | 565 | 759 | 840 | 721 | 142 | 560 | 506 | 517 | 528 | 28 | 527 | 650 | 557 | 578 | 65 | 706 | 765 | 588 | 686 | 90 | 699 | 846 | 709 | 751 | 82 | 846 | 866 | 458 | 723 | 230 |
| | **IGHD2-1** | 649 | 724 | 841 | 738 | 97 | 723 | 568 | 603 | 632 | 81 | 589 | 641 | 601 | 611 | 27 | 730 | 862 | 740 | 777 | 74 | 710 | 759 | 735 | 735 | 25 | 838 | 735 | 495 | 689 | 176 |
| | **IGHD2-4/2-9** | 1185 | 1243 | 1435 | 1288 | 131 | 1191 | 1063 | 1170 | 1141 | 69 | 957 | 1231 | 1153 | 1113 | 141 | 1426 | 1392 | 1258 | 1359 | 89 | 1131 | 1316 | 1200 | 1216 | 94 | 1282 | 1251 | 751 | 1094 | 298 |
| | **IGHD2-14** | 1027 | 1166 | 1307 | 1167 | 140 | 1382 | 1291 | 1243 | 1306 | 70 | 1086 | 1316 | 1273 | 1225 | 123 | 1247 | 1297 | 1060 | 1202 | 125 | 1022 | 1291 | 1305 | 1206 | 159 | 1307 | 1086 | 745 | 1046 | 283 |
| | **IGHD2-10** | 873 | 855 | 949 | 892 | 50 | 1291 | 1267 | 1189 | 1249 | 53 | 961 | 1196 | 1032 | 1063 | 121 | 949 | 1107 | 1004 | 1020 | 80 | 843 | 979 | 1042 | 955 | 102 | 929 | 812 | 569 | 770 | 184 |
| | **IGHD2-11** | 571 | 572 | 629 | 591 | 33 | 716 | 691 | 673 | 694 | 22 | 447 | 715 | 637 | 600 | 138 | 718 | 721 | 608 | 682 | 64 | 601 | 631 | 523 | 585 | 56 | 617 | 507 | 412 | 512 | 103 |
| | **IGHD3-2** | 218 | 230 | 325 | 258 | 59 | 376 | 343 | 411 | 377 | 34 | 331 | 364 | 396 | 364 | 32 | 406 | 414 | 329 | 383 | 47 | 258 | 311 | 303 | 291 | 29 | 397 | 443 | 274 | 371 | 87 |
| | **IGHD4-1 (DQ52)** | 495 | 517 | 613 | 542 | 63 | 778 | 724 | 779 | 761 | 31 | 567 | 654 | 596 | 606 | 45 | 734 | 639 | 633 | 669 | 57 | 557 | 620 | 543 | 573 | 41 | 597 | 700 | 495 | 597 | 103 |

Notes:
1. Data were nomalized to 78,091 total reads.
2. Proximal V and D segments are denoted in bold.

**Supplementary Table 2 | Productive and non-productive VHDJH junctions in WT and IGCR1/CBEs mutated pro-B cells.**

| | | WT (n=3) | | | | | | CBE1&2 -/- (n=3) | | | | | | CBE1 -/- (n=3) | | | | | | CBE2 -/- (n=3) | | | | | | CBE1 inv/inv (n=3) | | | | | | CBE2 inv/inv (n=3) | | | | | |
|---|---|---|---|---|---|---|---|---|---|---|---|---|---|---|---|---|---|---|---|---|---|---|---|---|---|---|---|---|---|---|---|---|---|---|---|---|---|
| | | Productive | | | Non-productive | | | Productive | | | Non-productive | | | | | | | | | | | | | | | | | | | | | | | | | | |
| VH domains | | #1 | #2 | #3 | #1 | #2 | #3 | #1 | #2 | #3 | #1 | #2 | #3 | #1 | #2 | #3 | #1 | #2 | #3 | #1 | #2 | #3 | #1 | #2 | #3 | #1 | #2 | #3 | #1 | #2 | #3 | #1 | #2 | #3 | #1 | #2 | #3 |
| | Ighv1-84 | 11 | 14 | 10 | 13 | 16 | 15 | 1 | 0 | 7 | 1 | 0 | 6 | 4 | 3 | 2 | 1 | 5 | 5 | 16 | 2 | 3 | 6 | 3 | 2 | 3 | 2 | 6 | 8 | 13 | 7 | 6 | 12 | 6 | 6 | 8 | 0 |
| | Ighv1-82 | 1 | 1 | 2 | 1 | 3 | 1 | 1 | 1 | 0 | 0 | 2 | 1 | 1 | 1 | 0 | 2 | 2 | 0 | 5 | 0 | 2 | 1 | 0 | 0 | 2 | 2 | 0 | 3 | 1 | 0 | 4 | 3 | 0 | 1 | 1 | 2 |
| | Ighv1-81 | 0 | 3 | 6 | 3 | 1 | 2 | 2 | 0 | 1 | 1 | 0 | 0 | 0 | 3 | 0 | 1 | 0 | 2 | 2 | 0 | 0 | 1 | 0 | 0 | 3 | 2 | 2 | 2 | 4 | 1 | 4 | 3 | 3 | 2 | 2 | 2 |
| | Ighv1-80 | 12 | 10 | 6 | 6 | 12 | 13 | 2 | 1 | 1 | 0 | 3 | 1 | 4 | 3 | 5 | 5 | 2 | 0 | 12 | 4 | 4 | 22 | 1 | 0 | 10 | 8 | 12 | 6 | 4 | 3 | 11 | 22 | 12 | 7 | 11 | 7 |
| | Ighv1-78 | 7 | 3 | 2 | 3 | 3 | 3 | 1 | 0 | 1 | 0 | 0 | 0 | 2 | 4 | 2 | 0 | 2 | 0 | 1 | 1 | 2 | 0 | 1 | 0 | 5 | 2 | 6 | 2 | 3 | 4 | 7 | 7 | 9 | 6 | 7 | 1 |
| | Ighv1-77 | 9 | 4 | 4 | 7 | 4 | 7 | 1 | 1 | 2 | 1 | 2 | 1 | 3 | 3 | 3 | 0 | 3 | 2 | 10 | 2 | 2 | 1 | 3 | 0 | 0 | 4 | 10 | 5 | 4 | 9 | 14 | 10 | 5 | 5 | 6 | 2 |
| | Ighv1-76 | 7 | 14 | 1 | 3 | 7 | 10 | 2 | 0 | 1 | 2 | 1 | 0 | 9 | 3 | 1 | 0 | 5 | 2 | 16 | 13 | 2 | 8 | 7 | 2 | 7 | 4 | 5 | 4 | 3 | 11 | 17 | 22 | 9 | 6 | 5 | 6 |
| | Ighv1-75 | 6 | 0 | 3 | 3 | 1 | 0 | 0 | 0 | 2 | 1 | 0 | 0 | 2 | 2 | 0 | 1 | 1 | 0 | 3 | 3 | 0 | 1 | 0 | 0 | 1 | 1 | 2 | 2 | 3 | 2 | 0 | 5 | 2 | 3 | 1 | 0 |
| | Ighv1-74 | 3 | 3 | 1 | 0 | 1 | 2 | 0 | 0 | 0 | 0 | 0 | 0 | 3 | 2 | 0 | 1 | 0 | 4 | 2 | 2 | 0 | 1 | 1 | 0 | 0 | 3 | 2 | 2 | 4 | 4 | 0 | 5 | 1 | 1 | 5 | 4 |
| | Ighv1-72 | 1 | 1 | 0 | 0 | 0 | 0 | 0 | 0 | 0 | 0 | 0 | 0 | 0 | 0 | 1 | 0 | 0 | 0 | 0 | 0 | 0 | 0 | 0 | 0 | 0 | 0 | 0 | 1 | 1 | 0 | 0 | 1 | 0 | 0 | 1 | 0 |
| | Ighv1-71 | 1 | 1 | 0 | 0 | 0 | 0 | 0 | 0 | 0 | 0 | 0 | 1 | 1 | 0 | 1 | 0 | 0 | 0 | 0 | 0 | 0 | 0 | 0 | 0 | 0 | 3 | 1 | 0 | 0 | 0 | 0 | 0 | 0 | 0 | 3 | 0 |
| | Ighv8-12 | 28 | 18 | 31 | 14 | 22 | 22 | 9 | 6 | 15 | 5 | 6 | 6 | 15 | 18 | 18 | 5 | 8 | 4 | 49 | 21 | 11 | 16 | 14 | 11 | 16 | 22 | 19 | 9 | 8 | 13 | 34 | 44 | 29 | 15 | 22 | 19 |
| | Ighv1-69 | 2 | 7 | 4 | 4 | 4 | 4 | 1 | 0 | 0 | 0 | 0 | 0 | 3 | 3 | 1 | 1 | 2 | 3 | 2 | 3 | 2 | 0 | 1 | 0 | 3 | 9 | 6 | 3 | 9 | 3 | 0 | 3 | 1 | 3 | 6 | 0 |
| | Ighv1-67 | 30 | 31 | 20 | 28 | 26 | 38 | 8 | 10 | 8 | 2 | 5 | 1 | 17 | 11 | 8 | 7 | 15 | 8 | 41 | 18 | 27 | 23 | 14 | 14 | 22 | 17 | 15 | 19 | 8 | 21 | 17 | 54 | 37 | 21 | 27 | 15 |
| J558/3609 | Ighv1-66 | 1 | 5 | 0 | 6 | 1 | 1 | 0 | 0 | 0 | 0 | 0 | 0 | 5 | 0 | 2 | 0 | 0 | 0 | 0 | 3 | 0 | 3 | 1 | 0 | 2 | 2 | 1 | 0 | 0 | 1 | 1 | 1 | 0 | 1 | 1 | 3 |
| | Ighv8-11 | 13 | 13 | 14 | 6 | 10 | 20 | 4 | 1 | 3 | 3 | 0 | 2 | 7 | 7 | 7 | 5 | 5 | 4 | 57 | 15 | 6 | 10 | 4 | 8 | 10 | 12 | 5 | 3 | 9 | 9 | 13 | 17 | 29 | 7 | 14 | 7 |
| | Ighv1-64 | 17 | 26 | 10 | 11 | 13 | 20 | 8 | 3 | 4 | 0 | 2 | 1 | 10 | 12 | 14 | 5 | 7 | 7 | 52 | 13 | 9 | 23 | 7 | 14 | 17 | 23 | 21 | 18 | 15 | 19 | 23 | 37 | 17 | 23 | 37 | 5 |
| | Ighv1-63 | 3 | 10 | 6 | 1 | 10 | 3 | 1 | 4 | 1 | 1 | 0 | 1 | 2 | 3 | 1 | 0 | 1 | 2 | 11 | 7 | 2 | 5 | 3 | 3 | 3 | 4 | 0 | 1 | 3 | 3 | 3 | 14 | 2 | 4 | 9 | 6 |
| | Ighv1-62-3 | 4 | 5 | 7 | 3 | 4 | 3 | 1 | 1 | 0 | 1 | 2 | 4 | 3 | 2 | 4 | 4 | 2 | 0 | 6 | 0 | 3 | 4 | 1 | 1 | 3 | 4 | 0 | 3 | 2 | 4 | 8 | 5 | 5 | 2 | 3 | 2 |
| | Ighv1-61 | 6 | 4 | 3 | 4 | 3 | 0 | 0 | 0 | 0 | 0 | 1 | 0 | 3 | 4 | 1 | 2 | 0 | 0 | 3 | 0 | 1 | 4 | 2 | 1 | 3 | 5 | 3 | 0 | 1 | 5 | 1 | 6 | 4 | 2 | 4 | 0 |
| | Ighv1-59 | 4 | 6 | 3 | 5 | 2 | 4 | 0 | 1 | 4 | 0 | 1 | 0 | 4 | 1 | 2 | 1 | 0 | 0 | 4 | 5 | 2 | 1 | 0 | 0 | 1 | 6 | 4 | 1 | 1 | 6 | 3 | 6 | 3 | 3 | 3 | 1 |
| | Ighv1-58 | 2 | 0 | 1 | 3 | 0 | 1 | 0 | 0 | 0 | 0 | 0 | 0 | 0 | 1 | 1 | 0 | 0 | 1 | 0 | 0 | 0 | 0 | 0 | 0 | 2 | 0 | 0 | 0 | 0 | 1 | 2 | 2 | 2 | 0 | 0 | 3 |
| | Ighv8-8 | 0 | 2 | 2 | 2 | 1 | 0 | 0 | 0 | 0 | 1 | 0 | 0 | 2 | 1 | 1 | 0 | 0 | 1 | 0 | 0 | 0 | 0 | 0 | 0 | 0 | 1 | 1 | 0 | 0 | 1 | 1 | 1 | 0 | 0 | 0 | 1 |
| | Ighv1-56 | 1 | 1 | 0 | 3 | 8 | 4 | 0 | 0 | 0 | 0 | 0 | 0 | 2 | 0 | 0 | 0 | 1 | 3 | 1 | 2 | 0 | 6 | 0 | 2 | 2 | 2 | 1 | 0 | 1 | 0 | 1 | 0 | 4 | 1 | 3 | 0 |
| | Ighv1-55 | 16 | 9 | 10 | 10 | 11 | 10 | 1 | 5 | 4 | 1 | 2 | 1 | 5 | 5 | 5 | 5 | 6 | 1 | 24 | 9 | 3 | 16 | 4 | 3 | 6 | 14 | 6 | 9 | 5 | 9 | 8 | 19 | 16 | 13 | 18 | 5 |
| | Ighv1-54 | 15 | 20 | 22 | 20 | 29 | 15 | 2 | 7 | 2 | 1 | 2 | 5 | 11 | 20 | 17 | 6 | 3 | 6 | 15 | 14 | 15 | 27 | 4 | 7 | 23 | 21 | 15 | 11 | 12 | 13 | 25 | 34 | 32 | 21 | 15 | 2 |
| | Ighv1-53 | 7 | 5 | 15 | 4 | 8 | 4 | 0 | 3 | 3 | 1 | 2 | 1 | 6 | 5 | 2 | 0 | 3 | 0 | 3 | 2 | 2 | 2 | 0 | 0 | 8 | 9 | 4 | 5 | 5 | 2 | 3 | 2 | 0 | 6 | 0 | 2 |
| | Ighv1-52 | 3 | 2 | 3 | 1 | 6 | 1 | 0 | 1 | 0 | 0 | 0 | 0 | 4 | 3 | 1 | 1 | 0 | 2 | 5 | 5 | 0 | 0 | 2 | 4 | 2 | 2 | 0 | 2 | 3 | 3 | 2 | 7 | 7 | 4 | 1 | 2 |
| | Ighv1-50 | 2 | 2 | 2 | 1 | 1 | 0 | 0 | 0 | 0 | 0 | 1 | 0 | 0 | 1 | 0 | 0 | 2 | 1 | 2 | 1 | 2 | 3 | 1 | 0 | 0 | 1 | 1 | 0 | 1 | 6 | 4 | 2 | 2 | 1 | 2 | 1 |
| | Ighv8-4 | 1 | 2 | 1 | 6 | 4 | 7 | 0 | 0 | 0 | 0 | 1 | 1 | 1 | 1 | 1 | 0 | 1 | 3 | 2 | 3 | 3 | 1 | 5 | 2 | 0 | 3 | 2 | 2 | 4 | 3 | 2 | 0 | 0 | 3 | 0 | 2 |
| | Ighv1-47 | 13 | 4 | 6 | 31 | 14 | 13 | 1 | 1 | 0 | 1 | 0 | 1 | 1 | 1 | 4 | 2 | 6 | 7 | 4 | 5 | 3 | 17 | 11 | 16 | 2 | 3 | 8 | 9 | 8 | 19 | 9 | 2 | 1 | 18 | 10 | 8 |
| | Ighv1-43 | 1 | 7 | 2 | 7 | 4 | 3 | 2 | 0 | 0 | 0 | 0 | 0 | 3 | 2 | 0 | 2 | 2 | 2 | 5 | 5 | 0 | 4 | 2 | 2 | 3 | 3 | 1 | 3 | 4 | 4 | 0 | 5 | 0 | 2 | 7 | 4 |
| | Ighv1-42 | 13 | 10 | 6 | 9 | 9 | 13 | 4 | 2 | 5 | 1 | 2 | 4 | 5 | 6 | 6 | 2 | 9 | 3 | 18 | 16 | 10 | 11 | 9 | 6 | 6 | 12 | 15 | 9 | 12 | 14 | 8 | 19 | 12 | 12 | 16 | 6 |
| | Ighv1-39 | 3 | 11 | 5 | 10 | 2 | 6 | 4 | 8 | 2 | 0 | 0 | 2 | 7 | 3 | 6 | 1 | 5 | 0 | 10 | 12 | 2 | 7 | 6 | 5 | 8 | 4 | 7 | 5 | 6 | 3 | 10 | 14 | 5 | 12 | 6 | 1 |
| | Ighv1-37 | 3 | 0 | 1 | 2 | 2 | 2 | 1 | 1 | 1 | 0 | 0 | 0 | 2 | 4 | 2 | 0 | 0 | 0 | 3 | 1 | 1 | 1 | 0 | 1 | 2 | 1 | 2 | 2 | 4 | 0 | 2 | 3 | 1 | 2 | 1 | 0 |
| | Ighv1-34 | 5 | 5 | 10 | 8 | 5 | 11 | 0 | 4 | 0 | 0 | 0 | 0 | 3 | 3 | 1 | 4 | 4 | 3 | 10 | 2 | 3 | 13 | 1 | 2 | 7 | 5 | 7 | 6 | 15 | 6 | 4 | 15 | 2 | 5 | 5 | 0 |
| | Ighv1-26 | 21 | 14 | 14 | 13 | 13 | 11 | 2 | 3 | 1 | 3 | 0 | 2 | 4 | 6 | 6 | 4 | 4 | 6 | 29 | 9 | 12 | 16 | 12 | 2 | 8 | 11 | 7 | 11 | 12 | 12 | 10 | 32 | 11 | 21 | 8 | 8 |
| | Ighv1-23 | 2 | 4 | 1 | 0 | 1 | 0 | 0 | 1 | 1 | 0 | 1 | 0 | 4 | 1 | 0 | 0 | 0 | 0 | 1 | 6 | 0 | 1 | 0 | 0 | 2 | 4 | 1 | 6 | 5 | 0 | 2 | 3 | 2 | 0 | 9 | 1 |
| | Ighv1-22 | 2 | 3 | 1 | 1 | 1 | 0 | 1 | 1 | 0 | 0 | 0 | 0 | 0 | 1 | 0 | 0 | 0 | 0 | 2 | 6 | 1 | 0 | 4 | 0 | 0 | 0 | 1 | 0 | 1 | 0 | 3 | 2 | 2 | 2 | 1 | 1 |
| | Ighv1-19 | 6 | 13 | 6 | 9 | 2 | 6 | 1 | 1 | 0 | 0 | 0 | 0 | 8 | 1 | 1 | 1 | 1 | 2 | 14 | 9 | 6 | 7 | 2 | 2 | 5 | 8 | 6 | 5 | 7 | 6 | 7 | 15 | 12 | 3 | 7 | 3 |
| J558 | Ighv1-18 | 17 | 7 | 19 | 12 | 21 | 8 | 3 | 3 | 0 | 0 | 2 | 1 | 9 | 9 | 3 | 5 | 3 | 1 | 37 | 14 | 9 | 15 | 8 | 2 | 12 | 6 | 12 | 18 | 9 | 9 | 5 | 28 | 19 | 14 | 18 | 11 |
| | Ighv1-14 | 18 | 25 | 22 | 27 | 22 | 22 | 3 | 9 | 3 | 2 | 1 | 8 | 17 | 11 | 17 | 6 | 10 | 17 | 31 | 12 | 15 | 15 | 25 | 25 | 8 | 17 | 29 | 15 | 13 | 25 | 26 | 60 | 26 | 27 | 27 | 23 |
| | Ighv1-12 | 13 | 11 | 11 | 7 | 9 | 9 | 3 | 0 | 4 | 0 | 3 | 1 | 6 | 6 | 7 | 0 | 4 | 1 | 6 | 4 | 3 | 8 | 3 | 5 | 6 | 2 | 7 | 1 | 7 | 6 | 9 | 8 | 6 | 1 | 7 | 4 |
| | Ighv1-9 | 37 | 40 | 34 | 29 | 29 | 21 | 12 | 9 | 8 | 5 | 7 | 4 | 7 | 12 | 19 | 13 | 12 | 7 | 48 | 11 | 11 | 41 | 6 | 21 | 22 | 40 | 39 | 14 | 23 | 26 | 31 | 55 | 30 | 36 | 22 | 14 |
| | Ighv15-2 | 12 | 25 | 15 | 47 | 33 | 29 | 3 | 5 | 2 | 2 | 4 | 2 | 12 | 14 | 12 | 18 | 10 | 16 | 20 | 15 | 16 | 56 | 36 | 13 | 11 | 25 | 14 | 24 | 25 | 31 | 13 | 29 | 20 | 31 | 27 | 9 |
| | Ighv1-7 | 1 | 1 | 0 | 0 | 0 | 0 | 0 | 0 | 0 | 0 | 0 | 0 | 0 | 0 | 1 | 0 | 0 | 0 | 0 | 0 | 0 | 0 | 0 | 0 | 0 | 0 | 2 | 0 | 0 | 0 | 0 | 0 | 0 | 0 | 0 | 0 |
| | Ighv10-3 | 20 | 8 | 8 | 20 | 24 | 18 | 2 | 4 | 2 | 3 | 1 | 4 | 3 | 0 | 10 | 12 | 10 | 9 | 9 | 5 | 2 | 26 | 9 | 20 | 4 | 8 | 6 | 17 | 11 | 32 | 8 | 10 | 5 | 18 | 33 | 13 |
| | Ighv1-5 | 30 | 33 | 33 | 30 | 17 | 32 | 3 | 7 | 16 | 4 | 1 | 0 | 21 | 17 | 12 | 12 | 16 | 16 | 65 | 28 | 23 | 41 | 9 | 22 | 18 | 16 | 23 | 27 | 21 | 26 | 30 | 55 | 38 | 30 | 29 | 14 |
| | Ighv1-4 | 21 | 20 | 15 | 26 | 9 | 11 | 8 | 4 | 8 | 2 | 0 | 5 | 8 | 12 | 12 | 6 | 11 | 10 | 41 | 41 | 6 | 34 | 3 | 10 | 14 | 19 | 17 | 11 | 21 | 14 | 25 | 26 | 19 | 5 | 27 | 15 |
| | Ighv1-2P | 23 | 18 | 15 | 50 | 36 | 40 | 4 | 5 | 3 | 4 | 10 | 11 | 15 | 14 | 6 | 14 | 18 | 22 | 33 | 21 | 15 | 51 | 22 | 30 | 16 | 21 | 11 | 31 | 35 | 29 | 23 | 35 | 23 | 46 | 50 | 26 |
| | Ighv8-2 | 12 | 11 | 9 | 44 | 41 | 29 | 1 | 0 | 2 | 5 | 4 | 7 | 2 | 1 | 4 | 22 | 17 | 14 | 16 | 10 | 3 | 33 | 22 | 22 | 8 | 16 | 4 | 35 | 37 | 42 | 10 | 18 | 4 | 20 | 47 | 27 |
| | IGHV9-4 | 13 | 9 | 11 | 10 | 12 | 12 | 2 | 0 | 4 | 3 | 1 | 8 | 3 | 8 | 6 | 2 | 2 | 5 | 18 | 10 | 16 | 18 | 4 | 7 | 13 | 22 | 16 | 7 | 11 | 17 | 9 | 12 | 10 | 11 | 8 | 3 |
| | IGHV14-4 | 14 | 15 | 13 | 10 | 26 | 26 | 5 | 5 | 6 | 3 | 7 | 2 | 13 | 10 | 11 | 5 | 6 | 8 | 14 | 21 | 2 | 10 | 8 | 8 | 13 | 20 | 15 | 21 | 27 | 18 | 16 | 39 | 29 | 18 | 23 | 16 |
| | IGHV7-3 | 21 | 26 | 29 | 13 | 20 | 14 | 5 | 7 | 6 | 6 | 4 | 2 | 21 | 16 | 15 | 7 | 7 | 11 | 35 | 14 | 14 | 32 | 11 | 10 | 24 | 19 | 19 | 14 | 11 | 17 | 21 | 34 | 11 | 16 | 13 | 9 |
| | IGHV9-2-1 | 25 | 38 | 22 | 22 | 27 | 15 | 10 | 5 | 3 | 3 | 5 | 5 | 17 | 16 | 10 | 6 | 5 | 10 | 37 | 12 | 27 | 27 | 19 | 13 | 34 | 34 | 27 | 22 | 21 | 16 | 33 | 50 | 17 | 22 | 8 | 15 |
| | IGHV9-3-1 | 18 | 19 | 14 | 13 | 9 | 15 | 3 | 5 | 4 | 3 | 1 | 1 | 13 | 20 | 12 | 7 | 3 | 7 | 22 | 21 | 4 | 8 | 2 | 10 | 10 | 7 | 13 | 10 | 15 | 18 | 15 | 36 | 12 | 10 | 9 | 2 |
| | IGHV12-1-1 | 11 | 10 | 7 | 9 | 10 | 12 | 0 | 2 | 1 | 3 | 3 | 0 | 2 | 7 | 3 | 11 | 8 | 2 | 9 | 7 | 6 | 7 | 7 | 1 | 6 | 9 | 3 | 7 | 7 | 8 | 6 | 8 | 5 | 14 | 8 | 3 |
| | IGHV9-3 | 9 | 5 | 6 | 7 | 3 | 7 | 3 | 4 | 2 | 3 | 1 | 2 | 5 | 2 | 6 | 5 | 1 | 3 | 9 | 1 | 12 | 6 | 3 | 0 | 8 | 14 | 8 | 6 | 3 | 12 | 3 | 17 | 5 | 3 | 5 | 4 |
| | IGHV9-2 | 0 | 4 | 2 | 11 | 3 | 10 | 0 | 0 | 0 | 0 | 0 | 0 | 2 | 0 | 2 | 0 | 9 | 0 | 0 | 3 | 3 | 6 | 9 | 8 | 2 | 3 | 1 | 10 | 4 | 7 | 2 | 2 | 2 | 10 | 7 | 3 |
| Middle | IGHV9-1 | 6 | 8 | 7 | 5 | 8 | 13 | 2 | 2 | 2 | 2 | 0 | 2 | 2 | 3 | 4 | 3 | 1 | 0 | 13 | 6 | 3 | 7 | 5 | 1 | 6 | 4 | 5 | 6 | 11 | 7 | 12 | 18 | 7 | 3 | 3 | 3 |
| | IGHV14-3 | 22 | 33 | 25 | 16 | 35 | 23 | 3 | 12 | 6 | 5 | 14 | 4 | 24 | 32 | 12 | 14 | 9 | 7 | 23 | 40 | 11 | 15 | 5 | 2 | 32 | 52 | 38 | 12 | 36 | 24 | 24 | 53 | 11 | 27 | 8 | 7 |
| | IGHV11-2 | 10 | 7 | 7 | 26 | 23 | 22 | 2 | 0 | 4 | 4 | 2 | 11 | 3 | 5 | 4 | 9 | 18 | 22 | 9 | 16 | 2 | 19 | 20 | 16 | 6 | 8 | 2 | 20 | 16 | 13 | 4 | 9 | 8 | 21 | 20 | 16 |
| | IGHV3-2 | 25 | 35 | 41 | 35 | 37 | 36 | 7 | 22 | 9 | 4 | 8 | 8 | 28 | 25 | 17 | 9 | 23 | 16 | 38 | 7 | 10 | 24 | 5 | 12 | 32 | 51 | 54 | 27 | 33 | 30 | 18 | 31 | 14 | 24 | 9 | 7 |
| | IGHV4-2 | 7 | 8 | 11 | 14 | 11 | 13 | 6 | 1 | 1 | 8 | 2 | 3 | 1 | 11 | 3 | 11 | 10 | 12 | 4 | 7 | 2 | 13 | 7 | 3 | 3 | 11 | 5 | 10 | 18 | 23 | 3 | 4 | 2 | 15 | 15 | 3 |
| | IGHV3-1 | 10 | 11 | 8 | 19 | 8 | 13 | 4 | 7 | 2 | 0 | 2 | 2 | 9 | 8 | 10 | 10 | 4 | 8 | 13 | 6 | 12 | 11 | 5 | 4 | 14 | 11 | 8 | 8 | 14 | 13 | 7 | 17 | 4 | 7 | 8 | 3 |
| | IGHV4-1 | 15 | 14 | 30 | 7 | 19 | 21 | 5 | 6 | 7 | 4 | 4 | 2 | 16 | 15 | 16 | 12 | 12 | 10 | 24 | 15 | 9 | 15 | 10 | 1 | 24 | 17 | 21 | 16 | 20 | 17 | 17 | 18 | 7 | 17 | 8 | 6 |
| | IGHV14-1 | 5 | 12 | 2 | 10 | 15 | 11 | 2 | 1 | 2 | 1 | 3 | 3 | 13 | 9 | 8 | 3 | 1 | 3 | 14 | 20 | 11 | 5 | 3 | 5 | 4 | 10 | 13 | 12 | 10 | 6 | 13 | 17 | 11 | 7 | 5 | 8 |
| | IGHV7-1 | 37 | 33 | 25 | 23 | 34 | 28 | 23 | 6 | 8 | 3 | 10 | 5 | 20 | 26 | 31 | 14 | 16 | 22 | 67 | 31 | 17 | 33 | 22 | 16 | 28 | 35 | 18 | 24 | 29 | 26 | 43 | 47 | 30 | 12 | 21 | 7 |
| | IGHV2-9 | 30 | 29 | 56 | 38 | 36 | 40 | 12 | 19 | 21 | 7 | 15 | 4 | 27 | 31 | 35 | 10 | 38 | 37 | 64 | 31 | 30 | 31 | 24 | 29 | 38 | 57 | 48 | 29 | 50 | 43 | 28 | 64 | 25 | 26 | 31 | 20 |
| | IGHV5-17 | 29 | 15 | 13 | 24 | 23 | 30 | 5 | 3 | 7 | 5 | 10 | 2 | 23 | 18 | 15 | 5 | 10 | 16 | 31 | 23 | 19 | 25 | 17 | 8 | 18 | 25 | 27 | 22 | 21 | 20 | 22 | 37 | 26 | 14 | 33 | 11 |
| | IGHV5-15 | 6 | 4 | 1 | 12 | 9 | 6 | 5 | 3 | 0 | 3 | 1 | 3 | 5 | 4 | 3 | 1 | 0 | 4 | 12 | 14 | 1 | 5 | 12 | 14 | 4 | 17 | 7 | 8 | 4 | 11 | 9 | 17 | 13 | 16 | 8 | 9 |
| | IGHV2-6-7 | 30 | 41 | 29 | 33 | 21 | 34 | 14 | 10 | 10 | 8 | 4 | 4 | 22 | 19 | 23 | 7 | 23 | 14 | 39 | 38 | 25 | 26 | 9 | 18 | 32 | 31 | 29 | 32 | 27 | 30 | 28 | 33 | 33 | 22 | 11 | 16 |
| | IGHV2-6-6 | 0 | 4 | 4 | 0 | 0 | 1 | 0 | 0 | 0 | 1 | 1 | 1 | 1 | 0 | 2 | 0 | 1 | 0 | 5 | 0 | 1 | 2 | 0 | 0 | 0 | 1 | 1 | 0 | 0 | 0 | 1 | 0 | 0 | 0 | 0 | 0 |
| | IGHV2-6-5 | 9 | 1 | 6 | 4 | 6 | 2 | 0 | 4 | 0 | 0 | 0 | 1 | 1 | 5 | 4 | 1 | 2 | 3 | 3 | 3 | 7 | 2 | 1 | 1 | 14 | 7 | 3 | 2 | 4 | 4 | 3 | 5 | 3 | 0 | 2 | 5 |
| | IGHV2-2-2 | 4 | 4 | 5 | 5 | 5 | 17 | 0 | 5 | 1 | 1 | 0 | 1 | 4 | 6 | 5 | 1 | 5 | 2 | 15 | 8 | 11 | 9 | 3 | 0 | 6 | 5 | 9 | 2 | 8 | 5 | 14 | 12 | 6 | 7 | 5 | 1 |
| | IGHV2-6-4 | 10 | 10 | 13 | 7 | 15 | 15 | 0 | 2 | 5 | 1 | 3 | 4 | 8 | 2 | 11 | 2 | 4 | 6 | 17 | 9 | 2 | 9 | 1 | 1 | 7 | 7 | 5 | 5 | 5 | 3 | 5 | 7 | 8 | 3 | 7 | 2 |
| | IGHV5-6-5 | 11 | 10 | 14 | 15 | 18 | 17 | 5 | 3 | 5 | 2 | 0 | 0 | 11 | 8 | 8 | 6 | 3 | 2 | 11 | 14 | 13 | 2 | 5 | 0 | 11 | 11 | 17 | 8 | 12 | 13 | 7 | 14 | 4 | 7 | 9 | 9 |
| | IGHV5-12-2 | 13 | 16 | 6 | 7 | 8 | 10 | 6 | 3 | 7 | 5 | 0 | 2 | 12 | 12 | 6 | 3 | 2 | 9 | 38 | 22 | 13 | 13 | 15 | 3 | 9 | 11 | 9 | 5 | 9 | 5 | 17 | 14 | 17 | 9 | 18 | 4 |
| | IGHV5-6-4 | 9 | 11 | 12 | 11 | 3 | 6 | 2 | 0 | 2 | 5 | 0 | 5 | 6 | 9 | 7 | 4 | 5 | 1 | 33 | 8 | 23 | 8 | 6 | 6 | 12 | 15 | 13 | 9 | 7 | 11 | 13 | 19 | 11 | 23 | 7 | 4 |
| | IGHV2-6-3 | 1 | 2 | 2 | 8 | 4 | 8 | 0 | 1 | 0 | 0 | 2 | 0 | 1 | 1 | 0 | 3 | 4 | 0 | 0 | 0 | 0 | 2 | 3 | 9 | 1 | 0 | 3 | 2 | 7 | 10 | 2 | 1 | 0 | 5 | 14 | 8 |
| | IGHV5-6-3 | 3 | 12 | 9 | 3 | 5 | 17 | 1 | 2 | 2 | 3 | 4 | 3 | 3 | 1 | 0 | 1 | 6 | 2 | 16 | 4 | 2 | 3 | 5 | 2 | 2 | 4 | 4 | 9 | 14 | 3 | 3 | 5 | 4 | 5 | 6 | 1 |
| | IGHV5-12-1 | 7 | 7 | 12 | 9 | 9 | 10 | 1 | 0 | 2 | 1 | 3 | 1 | 6 | 2 | 6 | 2 | 2 | 2 | 6 | 8 | 4 | 2 | 7 | 3 | 4 | 6 | 15 | 6 | 6 | 7 | 2 | 7 | 5 | 9 | 2 | 6 |
| | IGHV5-9-5 | 0 | 1 | 1 | 1 | 7 | 2 | 0 | 0 | 1 | 1 | 0 | 0 | 1 | 0 | 2 | 0 | 0 | 2 | 7 | 2 | 0 | 0 | 0 | 0 | 3 | 1 | 1 | 0 | 1 | 0 | 0 | 3 | 0 | 1 | 2 | 4 |
| | IGHV2-6-2 | 7 | 0 | 1 | 2 | 2 | 3 | 0 | 0 | 0 | 0 | 0 | 0 | 1 | 0 | 1 | 0 | 0 | 5 | 1 | 0 | 1 | 0 | 1 | 1 | 1 | 0 | 1 | 5 | 2 | 1 | 3 | 0 | 3 | 1 | 3 | 0 |
| 7183/Q52 | IGHV5-6-2 | 7 | 11 | 14 | 9 | 8 | 11 | 2 | 2 | 2 | 1 | 1 | 2 | 7 | 9 | 6 | 0 | 4 | 9 | 2 | 5 | 7 | 4 | 12 | 15 | 5 | 7 | 10 | 6 | 4 | 5 | 9 | 18 | 14 | 8 | 9 | 10 |
| | IGHV2-9-2 | 2 | 7 | 15 | 10 | 5 | 5 | 1 | 2 | 1 | 0 | 2 | 0 | 6 | 4 | 4 | 5 | 1 | 1 | 2 | 5 | 4 | 18 | 5 | 4 | 6 | 10 | 5 | 1 | 4 | 3 | 6 | 13 | 6 | 6 | 2 | 6 |
| | IGHV5-9-4 | 8 | 8 | 18 | 16 | 16 | 16 | 2 | 8 | 0 | 6 | 1 | 4 | 7 | 3 | 7 | 2 | 2 | 2 | 22 | 15 | 12 | 3 | 14 | 14 | 4 | 9 | 6 | 8 | 4 | 14 | 13 | 9 | 7 | 16 | 7 | 14 |
| | IGHV2-4-1 | 6 | 5 | 1 | 4 | 7 | 4 | 0 | 1 | 0 | 3 | 0 | 0 | 0 | 6 | 0 | 0 | 5 | 3 | 5 | 1 | 3 | 1 | 3 | 0 | 2 | 1 | 5 | 4 | 6 | 5 | 5 | 4 | 3 | 6 | 4 | 3 |
| | IGHV5-9-3 | 22 | 15 | 8 | 10 | 19 | 15 | 3 | 11 | 6 | 1 | 4 | 1 | 3 | 5 | 5 | 3 | 5 | 5 | 14 | 20 | 4 | 28 | 9 | 8 | 11 | 15 | 10 | 12 | 8 | 7 | 21 | 29 | 14 | 12 | 12 | 12 |
| | IGHV5-9-2 | 2 | 4 | 4 | 0 | 2 | 2 | 5 | 0 | 0 | 2 | 0 | 0 | 5 | 4 | 5 | 3 | 0 | 1 | 6 | 9 | 4 | 3 | 2 | 1 | 1 | 0 | 5 | 1 | 4 | 3 | 2 | 7 | 2 | 2 | 3 | 5 |
| | IGHV2-6-1 | 3 | 4 | 5 | 5 | 8 | 7 | 0 | 0 | 0 | 2 | 1 | 0 | 2 | 3 | 1 | 0 | 3 | 3 | 1 | 3 | 1 | 6 | 1 | 2 | 1 | 3 | 4 | 4 | 2 | 4 | 0 | 1 | 2 | 4 | 5 | 0 |
| | IGHV5-9-1 | 13 | 19 | 21 | 22 | 22 | 31 | 9 | 6 | 8 | 2 | 5 | 4 | 21 | 23 | 6 | 7 | 15 | 10 | 22 | 47 | 15 | 35 | 37 | 26 | 17 | 40 | 28 | 22 | 23 | 17 | 32 | 46 | 32 | 22 | 26 | 15 |
| | IGHV2-6 | 2 | 4 | 1 | 7 | 3 | 7 | 0 | 1 | 0 | 1 | 1 | 2 | 2 | 5 | 0 | 0 | 3 | 0 | 0 | 2 | 3 | 5 | 4 | 9 | 4 | 2 | 2 | 8 | 9 | 5 | 1 | 3 | 2 | 3 | 9 | 2 |
| | IGHV5-12 | 23 | 28 | 24 | 21 | 43 | 27 | 5 | 9 | 13 | 3 | 4 | 7 | 10 | 5 | 10 | 9 | 9 | 5 | 38 | 66 | 61 | 56 | 38 | 49 | 7 | 19 | 12 | 21 | 26 | 22 | 19 | 44 | 32 | 46 | 38 | 26 |
| | IGHV5-9 | 2 | 4 | 7 | 2 | 9 | 0 | 0 | 0 | 0 | 1 | 2 | 1 | 5 | 6 | 3 | 1 | 5 | 2 | 9 | 14 | 1 | 20 | 2 | 20 | 4 | 1 | 6 | 2 | 2 | 1 | 2 | 8 | 4 | 8 | 7 | 4 |
| | IGHV2-4 | 5 | 2 | 3 | 0 | 5 | 5 | 2 | 2 | 1 | 2 | 5 | 0 | 3 | 3 | 1 | 6 | 8 | 6 | 1 | 1 | 4 | 2 | 8 | 8 | 0 | 2 | 0 | 3 | 7 | 1 | 0 | 7 | 1 | 3 | 3 | 4 |
| | IGHV5-6 | 11 | 14 | 11 | 57 | 45 | 59 | 7 | 7 | 7 | 10 | 5 | 9 | 16 | 4 | 5 | 9 | 9 | 9 | 23 | 52 | 49 | 94 | 84 | 81 | 2 | 15 | 17 | 17 | 18 | 12 | 24 | 30 | 9 | 92 | 54 | 16 |
| | IGHV2-3 | 27 | 38 | 16 | 52 | 65 | 71 | 8 | 13 | 14 | 24 | 33 | 26 | 12 | 16 | 8 | 29 | 27 | 24 | 42 | 55 | 43 | 120 | 115 | 102 | 14 | 14 | 17 | 19 | 17 | 27 | 23 | 44 | 29 | 94 | 67 | 25 |
| | IGHV5-4 | 30 | 38 | 25 | 65 | 136 | 85 | 21 | 18 | 24 | 95 | 90 | 92 | 30 | 28 | 30 | 73 | 121 | 109 | 58 | 117 | 100 | 221 | 178 | 153 | 15 | 12 | 24 | 52 | 62 | 56 | 38 | 55 | 39 | 170 | 129 | 58 |
| | IGHV2-2 | 43 | 44 | 41 | 144 | 211 | 182 | 57 | 89 | 73 | 559 | 539 | 509 | 109 | 103 | 71 | 520 | 611 | 627 | 132 | 163 | 110 | 434 | 405 | 369 | 60 | 58 | 83 | 277 | 303 | 297 | 89 | 78 | 66 | 276 | 259 | 141 |
| | IGHV5-2 | 47 | 66 | 71 | 273 | 335 | 328 | 940 | 879 | 896 | 5019 | 4380 | 4327 | 655 | 874 | 751 | 2932 | 4110 | 3798 | 175 | 224 | 136 | 862 | 859 | 758 | 233 | 283 | 305 | 1312 | 1502 | 1478 | 129 | 93 | 43 | 522 | 468 | 199 |

Notes:

1. Data were nomalized to 78,091 total reads.
2. Pseudo V are not listed in this table.

**Supplementary Table 3 | Absolute utilization of Igh VH and D segments in single Igh WAPL-degron v-Abl pro-B cells.**

| | | WT untreated (n=3) | | | | | WT+Dox/IAA (n=3) | | | | | 3'CBE del untreated (n=3) | | | | | 3'CBE del+Dox/IAA (n=3) | | | | | 3'CBE inv untreated (n=3) | | | | | 3'CBE inv +Dox/IAA (n=3) | | | | |
|---|---|---|---|---|---|---|---|---|---|---|---|---|---|---|---|---|---|---|---|---|---|---|---|---|---|---|---|---|---|---|---|
| **VH domains** | | #1 | #2 | #3 | Average | s.d. | #1 | #2 | #3 | Average | s.d. | #1 | #2 | #3 | Average | s.d. | #1 | #2 | #3 | Average | s.d. | #1 | #2 | #3 | Average | s.d. | #1 | #2 | #3 | Average | s.d. |
| | Ighv1-86P | 0 | 0 | 0 | 0 | 0 | 0 | 0 | 5 | 2 | 3 | 0 | 0 | 0 | 0 | 0 | 0 | 2 | 2 | 1 | 1 | 0 | 0 | 0 | 0 | 0 | 0 | 3 | 0 | 1 | 2 |
| | Ighv1-85 | 1 | 0 | 0 | 0 | 1 | 17 | 18 | 19 | 18 | 1 | 0 | 0 | 0 | 0 | 0 | 7 | 6 | 2 | 5 | 3 | 0 | 0 | 4 | 1 | 2 | 6 | 4 | 1 | 4 | 3 |
| | Ighv1-84 | 0 | 0 | 0 | 0 | 0 | 0 | 0 | 8 | 3 | 5 | 0 | 0 | 0 | 0 | 0 | 2 | 1 | 4 | 2 | 2 | 0 | 0 | 0 | 0 | 0 | 0 | 1 | 0 | 0 | 1 |
| | Ighv1-83P | 0 | 0 | 0 | 0 | 0 | 5 | 2 | 0 | 2 | 3 | 0 | 0 | 0 | 0 | 0 | 1 | 4 | 3 | 3 | 2 | 0 | 0 | 1 | 0 | 1 | 0 | 2 | 4 | 2 | 2 |
| | Ighv1-82 | 0 | 0 | 0 | 0 | 0 | 4 | 9 | 21 | 11 | 9 | 0 | 0 | 0 | 0 | 0 | 8 | 7 | 5 | 7 | 2 | 0 | 0 | 0 | 0 | 0 | 4 | 10 | 9 | 8 | 3 |
| | Ighv1-81 | 0 | 0 | 0 | 0 | 0 | 1 | 12 | 8 | 7 | 6 | 0 | 0 | 0 | 0 | 0 | 11 | 8 | 5 | 8 | 3 | 0 | 4 | 0 | 1 | 2 | 6 | 5 | 7 | 6 | 1 |
| | Ighv1-80 | 0 | 0 | 1 | 0 | 1 | 8 | 7 | 14 | 10 | 4 | 0 | 0 | 0 | 0 | 0 | 10 | 6 | 1 | 6 | 5 | 0 | 0 | 1 | 0 | 1 | 1 | 11 | 0 | 4 | 6 |
| | Ighv1-79P | 0 | 0 | 0 | 0 | 0 | 1 | 8 | 2 | 4 | 4 | 1 | 0 | 0 | 0 | 1 | 0 | 1 | 2 | 1 | 1 | 0 | 0 | 0 | 0 | 0 | 0 | 5 | 0 | 2 | 3 |
| | Ighv1-78 | 0 | 0 | 8 | 3 | 5 | 28 | 50 | 48 | 42 | 12 | 1 | 8 | 0 | 3 | 4 | 23 | 18 | 26 | 22 | 4 | 12 | 8 | 9 | 10 | 2 | 14 | 49 | 29 | 31 | 18 |
| | Ighv1-77 | 0 | 0 | 0 | 0 | 0 | 4 | 3 | 22 | 10 | 11 | 0 | 0 | 1 | 0 | 1 | 3 | 7 | 3 | 4 | 2 | 0 | 0 | 0 | 0 | 0 | 2 | 3 | 5 | 3 | 2 |
| | Ighv1-76 | 0 | 0 | 5 | 2 | 3 | 42 | 35 | 23 | 33 | 10 | 1 | 0 | 3 | 1 | 2 | 16 | 19 | 28 | 21 | 6 | 0 | 1 | 4 | 2 | 2 | 27 | 25 | 45 | 32 | 11 |
| | Ighv1-75 | 0 | 0 | 0 | 0 | 0 | 14 | 18 | 22 | 18 | 4 | 0 | 0 | 0 | 0 | 0 | 2 | 1 | 17 | 7 | 9 | 0 | 0 | 0 | 0 | 0 | 8 | 7 | 23 | 13 | 9 |
| | Ighv1-74 | 0 | 1 | 4 | 2 | 2 | 36 | 51 | 38 | 42 | 8 | 4 | 4 | 4 | 4 | 0 | 12 | 13 | 14 | 13 | 1 | 5 | 5 | 6 | 5 | 1 | 20 | 24 | 33 | 26 | 7 |
| | Ighv1-73P | 0 | 3 | 0 | 1 | 2 | 16 | 18 | 36 | 23 | 11 | 0 | 0 | 0 | 0 | 0 | 4 | 7 | 9 | 7 | 3 | 0 | 0 | 0 | 0 | 0 | 2 | 6 | 10 | 6 | 4 |
| | Ighv1-72 | 0 | 0 | 1 | 0 | 1 | 4 | 7 | 13 | 8 | 5 | 0 | 0 | 1 | 0 | 1 | 4 | 7 | 2 | 4 | 3 | 1 | 7 | 0 | 3 | 4 | 3 | 24 | 5 | 11 | 12 |
| | Ighv1-71 | 0 | 0 | 0 | 0 | 0 | 0 | 0 | 0 | 0 | 0 | 0 | 0 | 0 | 0 | 0 | 0 | 0 | 0 | 0 | 0 | 0 | 0 | 0 | 0 | 0 | 0 | 0 | 0 | 0 | 0 |
| | Ighv1-70P | 2 | 0 | 4 | 2 | 2 | 95 | 73 | 102 | 90 | 15 | 1 | 1 | 10 | 4 | 5 | 44 | 22 | 40 | 35 | 12 | 0 | 4 | 5 | 3 | 3 | 45 | 56 | 86 | 62 | 21 |
| | Ighv8-12 | 19 | 3 | 12 | 11 | 8 | 91 | 101 | 79 | 90 | 11 | 8 | 2 | 4 | 5 | 3 | 28 | 27 | 40 | 32 | 7 | 1 | 19 | 11 | 10 | 9 | 61 | 80 | 86 | 76 | 13 |
| | Ighv1-69 | 0 | 0 | 2 | 1 | 1 | 6 | 13 | 13 | 11 | 4 | 5 | 0 | 1 | 2 | 3 | 8 | 9 | 6 | 8 | 2 | 1 | 5 | 2 | 3 | 2 | 8 | 17 | 8 | 11 | 5 |
| | Ighv1-67 | 2 | 0 | 0 | 1 | 1 | 13 | 13 | 13 | 13 | 0 | 0 | 0 | 3 | 1 | 2 | 8 | 1 | 1 | 3 | 4 | 0 | 1 | 3 | 1 | 2 | 12 | 4 | 10 | 9 | 4 |
| | Ighv1-66 | 0 | 0 | 0 | 0 | 0 | 1 | 1 | 6 | 3 | 3 | 0 | 0 | 0 | 0 | 0 | 0 | 0 | 1 | 0 | 1 | 0 | 0 | 0 | 0 | 0 | 1 | 0 | 1 | 1 | 1 |
| J558/3609 | Ighv8-11 | 1 | 5 | 1 | 2 | 2 | 37 | 23 | 35 | 32 | 8 | 2 | 2 | 11 | 5 | 5 | 27 | 20 | 15 | 21 | 6 | 2 | 2 | 7 | 4 | 3 | 26 | 25 | 32 | 28 | 4 |
| | Ighv1-64 | 0 | 0 | 0 | 0 | 0 | 9 | 4 | 12 | 8 | 4 | 2 | 2 | 3 | 2 | 1 | 8 | 17 | 2 | 9 | 8 | 1 | 0 | 1 | 1 | 1 | 7 | 4 | 7 | 6 | 2 |
| | Ighv1-63 | 2 | 0 | 2 | 1 | 1 | 19 | 29 | 32 | 27 | 7 | 0 | 1 | 6 | 2 | 3 | 4 | 10 | 17 | 10 | 7 | 1 | 0 | 1 | 1 | 1 | 19 | 29 | 22 | 23 | 5 |
| | Ighv8-9 | 0 | 0 | 0 | 0 | 0 | 19 | 8 | 10 | 12 | 6 | 0 | 0 | 0 | 0 | 0 | 0 | 7 | 5 | 4 | 4 | 2 | 0 | 3 | 2 | 2 | 2 | 3 | 12 | 6 | 6 |
| | Ighv1-62-3 | 1 | 1 | 0 | 1 | 1 | 3 | 3 | 3 | 3 | 0 | 0 | 0 | 1 | 0 | 1 | 3 | 0 | 1 | 1 | 2 | 0 | 0 | 2 | 1 | 1 | 2 | 2 | 3 | 2 | 1 |
| | Ighv1-62-2 | 0 | 0 | 0 | 0 | 0 | 0 | 0 | 0 | 0 | 0 | 0 | 0 | 0 | 0 | 0 | 0 | 0 | 0 | 0 | 0 | 0 | 0 | 0 | 0 | 0 | 1 | 0 | 0 | 0 | 1 |
| | Ighv1-62P | 0 | 0 | 2 | 1 | 1 | 1 | 6 | 7 | 5 | 3 | 0 | 0 | 0 | 0 | 0 | 0 | 3 | 3 | 2 | 2 | 0 | 0 | 1 | 0 | 1 | 2 | 1 | 1 | 1 | 1 |
| | Ighv1-61 | 1 | 4 | 1 | 2 | 2 | 23 | 19 | 26 | 23 | 4 | 1 | 1 | 4 | 2 | 2 | 8 | 14 | 6 | 9 | 4 | 4 | 8 | 7 | 6 | 2 | 21 | 21 | 20 | 21 | 1 |
| | Ighv1-59 | 0 | 0 | 0 | 0 | 0 | 9 | 12 | 11 | 11 | 2 | 2 | 0 | 1 | 1 | 1 | 3 | 4 | 5 | 4 | 1 | 2 | 3 | 1 | 2 | 1 | 3 | 13 | 8 | 8 | 5 |
| | Ighv1-58 | 3 | 0 | 0 | 1 | 2 | 23 | 28 | 30 | 27 | 4 | 0 | 0 | 1 | 0 | 1 | 13 | 14 | 9 | 12 | 3 | 4 | 0 | 5 | 3 | 3 | 15 | 11 | 13 | 13 | 2 |
| | Ighv8-8 | 2 | 9 | 2 | 4 | 4 | 60 | 73 | 72 | 68 | 7 | 11 | 5 | 8 | 8 | 3 | 23 | 30 | 46 | 33 | 12 | 29 | 18 | 20 | 22 | 6 | 41 | 45 | 64 | 50 | 12 |
| | Ighv8-7P | 3 | 0 | 9 | 4 | 5 | 29 | 23 | 29 | 27 | 3 | 0 | 4 | 4 | 3 | 2 | 32 | 24 | 29 | 28 | 4 | 7 | 3 | 7 | 6 | 2 | 27 | 23 | 37 | 29 | 7 |
| | Ighv1-56 | 0 | 0 | 1 | 0 | 1 | 2 | 3 | 5 | 3 | 2 | 0 | 0 | 0 | 0 | 0 | 0 | 2 | 1 | 1 | 1 | 0 | 0 | 1 | 0 | 1 | 1 | 1 | 5 | 2 | 2 |
| | Ighv1-55 | 0 | 1 | 2 | 1 | 1 | 10 | 7 | 11 | 9 | 2 | 0 | 0 | 0 | 0 | 0 | 4 | 0 | 6 | 3 | 3 | 0 | 3 | 3 | 2 | 2 | 3 | 2 | 10 | 5 | 4 |
| | Ighv1-54 | 1 | 5 | 7 | 4 | 3 | 46 | 47 | 34 | 42 | 7 | 5 | 9 | 8 | 7 | 2 | 7 | 26 | 26 | 20 | 11 | 12 | 14 | 3 | 10 | 6 | 19 | 33 | 38 | 30 | 10 |
| | Ighv8-6 | 0 | 0 | 2 | 1 | 1 | 21 | 13 | 5 | 13 | 8 | 0 | 0 | 2 | 1 | 1 | 6 | 11 | 1 | 6 | 5 | 0 | 4 | 5 | 3 | 3 | 8 | 7 | 9 | 8 | 1 |
| | Ighv1-53 | 0 | 1 | 2 | 1 | 1 | 17 | 4 | 8 | 10 | 7 | 0 | 6 | 1 | 2 | 3 | 8 | 8 | 7 | 8 | 1 | 1 | 1 | 1 | 1 | 0 | 3 | 7 | 15 | 8 | 6 |
| | Ighv1-52 | 1 | 2 | 4 | 2 | 2 | 41 | 13 | 14 | 23 | 16 | 2 | 1 | 1 | 1 | 1 | 15 | 16 | 13 | 15 | 2 | 6 | 7 | 3 | 5 | 2 | 20 | 13 | 19 | 17 | 4 |
| | Ighv1-51P | 0 | 0 | 0 | 0 | 0 | 2 | 0 | 0 | 1 | 1 | 7 | 0 | 0 | 2 | 4 | 0 | 0 | 1 | 0 | 1 | 0 | 0 | 0 | 0 | 0 | 3 | 0 | 0 | 1 | 2 |
| | Ighv1-50 | 0 | 0 | 0 | 0 | 0 | 3 | 5 | 9 | 6 | 3 | 0 | 2 | 3 | 2 | 2 | 1 | 1 | 2 | 1 | 1 | 0 | 1 | 2 | 1 | 1 | 2 | 5 | 5 | 4 | 2 |
| | Ighv8-5 | 0 | 0 | 0 | 0 | 0 | 8 | 3 | 1 | 4 | 4 | 0 | 0 | 0 | 0 | 0 | 5 | 0 | 2 | 2 | 3 | 0 | 1 | 8 | 3 | 4 | 2 | 0 | 3 | 2 | 2 |
| | Ighv1-49 | 0 | 0 | 0 | 0 | 0 | 2 | 5 | 3 | 3 | 2 | 0 | 1 | 3 | 1 | 2 | 0 | 7 | 2 | 3 | 4 | 0 | 4 | 0 | 1 | 2 | 8 | 1 | 6 | 5 | 4 |
| | Ighv8-4 | 0 | 0 | 0 | 0 | 0 | 15 | 12 | 16 | 14 | 2 | 0 | 0 | 3 | 1 | 2 | 3 | 14 | 6 | 8 | 6 | 1 | 1 | 0 | 1 | 1 | 14 | 3 | 8 | 8 | 6 |
| | Ighv1-47 | 8 | 15 | 15 | 13 | 4 | 40 | 42 | 39 | 40 | 2 | 1 | 7 | 6 | 5 | 3 | 25 | 25 | 19 | 23 | 3 | 16 | 14 | 25 | 18 | 6 | 35 | 33 | 27 | 32 | 4 |
| | Ighv1-46P | 0 | 0 | 0 | 0 | 0 | 5 | 0 | 0 | 2 | 3 | 4 | 0 | 0 | 1 | 2 | 1 | 3 | 2 | 2 | 1 | 0 | 0 | 0 | 0 | 0 | 0 | 4 | 0 | 1 | 2 |
| | Ighv1-43 | 0 | 0 | 0 | 0 | 0 | 3 | 0 | 0 | 1 | 2 | 0 | 0 | 0 | 0 | 0 | 1 | 0 | 0 | 0 | 1 | 0 | 2 | 0 | 1 | 1 | 3 | 0 | 7 | 3 | 4 |
| | Ighv1-42 | 3 | 0 | 1 | 1 | 2 | 11 | 25 | 28 | 21 | 9 | 2 | 0 | 1 | 1 | 1 | 1 | 11 | 0 | 4 | 6 | 13 | 1 | 2 | 5 | 7 | 6 | 1 | 2 | 3 | 3 |
| | Ighv1-40P | 0 | 0 | 0 | 0 | 0 | 4 | 0 | 0 | 1 | 2 | 0 | 0 | 0 | 0 | 0 | 0 | 0 | 0 | 0 | 0 | 2 | 0 | 5 | 2 | 3 | 0 | 0 | 0 | 0 | 0 |
| | Ighv1-39 | 3 | 0 | 3 | 2 | 2 | 21 | 12 | 24 | 19 | 6 | 0 | 2 | 3 | 2 | 2 | 7 | 12 | 6 | 8 | 3 | 8 | 1 | 6 | 5 | 4 | 7 | 9 | 23 | 13 | 9 |
| | Ighv1-38P | 0 | 0 | 2 | 1 | 1 | 1 | 2 | 1 | 1 | 1 | 0 | 1 | 1 | 1 | 1 | 0 | 0 | 0 | 0 | 0 | 0 | 0 | 0 | 0 | 0 | 1 | 0 | 1 | 1 | 1 |
| | Ighv1-36 | 0 | 0 | 1 | 0 | 1 | 2 | 1 | 0 | 1 | 1 | 0 | 0 | 0 | 0 | 0 | 0 | 0 | 4 | 1 | 2 | 0 | 0 | 0 | 0 | 0 | 0 | 0 | 0 | 0 | 0 |
| | Ighv1-34 | 0 | 0 | 0 | 0 | 0 | 0 | 1 | 0 | 0 | 1 | 0 | 0 | 0 | 0 | 0 | 4 | 1 | 0 | 2 | 2 | 0 | 0 | 0 | 0 | 0 | 0 | 0 | 0 | 0 | 0 |
| | Ighv1-33P | 0 | 0 | 0 | 0 | 0 | 1 | 0 | 4 | 2 | 2 | 0 | 0 | 1 | 0 | 1 | 0 | 1 | 2 | 1 | 1 | 0 | 0 | 0 | 0 | 0 | 2 | 1 | 0 | 1 | 1 |
| | Ighv1-32P | 0 | 0 | 0 | 0 | 0 | 0 | 0 | 0 | 0 | 0 | 0 | 0 | 0 | 0 | 0 | 0 | 0 | 0 | 0 | 0 | 0 | 0 | 0 | 0 | 0 | 0 | 0 | 0 | 0 | 0 |
| | Ighv1-31 | 0 | 0 | 0 | 0 | 0 | 2 | 1 | 0 | 1 | 1 | 0 | 0 | 0 | 0 | 0 | 0 | 0 | 5 | 2 | 3 | 0 | 0 | 0 | 0 | 0 | 4 | 0 | 0 | 1 | 2 |
| | Ighv1-26 | 0 | 0 | 0 | 0 | 0 | 2 | 6 | 4 | 4 | 2 | 0 | 0 | 0 | 0 | 0 | 6 | 9 | 2 | 6 | 4 | 0 | 0 | 0 | 0 | 0 | 0 | 2 | 0 | 1 | 1 |
| | Ighv1-25P | 0 | 0 | 0 | 0 | 0 | 2 | 17 | 15 | 11 | 8 | 0 | 0 | 0 | 0 | 0 | 0 | 11 | 1 | 4 | 6 | 0 | 2 | 7 | 3 | 4 | 9 | 6 | 11 | 9 | 3 |
| | Ighv1-23 ORF | 0 | 0 | 0 | 0 | 0 | 1 | 7 | 5 | 4 | 3 | 1 | 1 | 0 | 1 | 1 | 2 | 5 | 4 | 4 | 2 | 0 | 0 | 0 | 0 | 0 | 0 | 5 | 4 | 3 | 3 |
| | Ighv1-22 | 1 | 1 | 0 | 1 | 1 | 4 | 7 | 10 | 7 | 3 | 0 | 0 | 1 | 0 | 1 | 0 | 0 | 1 | 0 | 1 | 0 | 0 | 0 | 0 | 0 | 0 | 2 | 0 | 1 | 1 |
| | Ighv1-21P | 0 | 0 | 0 | 0 | 0 | 0 | 7 | 7 | 5 | 4 | 0 | 0 | 0 | 0 | 0 | 0 | 3 | 0 | 1 | 2 | 0 | 0 | 0 | 0 | 0 | 8 | 6 | 0 | 5 | 4 |
| J558 | Ighv1-21-1P | 0 | 0 | 1 | 0 | 1 | 0 | 0 | 3 | 1 | 2 | 0 | 0 | 0 | 0 | 0 | 0 | 0 | 0 | 0 | 0 | 0 | 0 | 0 | 0 | 0 | 0 | 3 | 3 | 2 | 2 |
| | Ighv1-20 | 0 | 0 | 1 | 0 | 1 | 0 | 0 | 0 | 0 | 0 | 0 | 1 | 0 | 0 | 1 | 1 | 0 | 0 | 0 | 1 | 0 | 0 | 0 | 0 | 0 | 0 | 0 | 5 | 2 | 3 |
| | Ighv1-19 | 0 | 0 | 0 | 0 | 0 | 4 | 0 | 0 | 1 | 2 | 0 | 0 | 2 | 1 | 1 | 2 | 0 | 7 | 3 | 4 | 0 | 0 | 2 | 1 | 1 | 3 | 3 | 7 | 4 | 2 |
| | Ighv1-18 | 0 | 0 | 1 | 0 | 1 | 7 | 6 | 8 | 7 | 1 | 0 | 0 | 4 | 1 | 2 | 0 | 1 | 2 | 1 | 1 | 1 | 0 | 0 | 0 | 1 | 3 | 4 | 10 | 6 | 4 |
| | Ighv1-17 | 0 | 0 | 0 | 0 | 0 | 0 | 1 | 9 | 3 | 5 | 0 | 0 | 0 | 0 | 0 | 0 | 0 | 0 | 0 | 0 | 0 | 0 | 0 | 0 | 0 | 0 | 0 | 0 | 0 | 0 |
| | Ighv1-15 | 0 | 0 | 0 | 0 | 0 | 2 | 9 | 2 | 4 | 4 | 1 | 0 | 0 | 0 | 1 | 0 | 0 | 3 | 1 | 2 | 0 | 0 | 0 | 0 | 0 | 0 | 0 | 1 | 0 | 1 |
| | Ighv1-14 | 0 | 0 | 1 | 0 | 1 | 0 | 2 | 0 | 1 | 1 | 0 | 0 | 1 | 0 | 1 | 0 | 0 | 1 | 0 | 1 | 0 | 0 | 0 | 0 | 0 | 0 | 2 | 0 | 1 | 1 |
| | Ighv1-13P | 0 | 1 | 3 | 1 | 2 | 0 | 6 | 3 | 3 | 3 | 0 | 0 | 0 | 0 | 0 | 0 | 5 | 0 | 2 | 3 | 0 | 0 | 4 | 1 | 2 | 1 | 0 | 0 | 0 | 1 |
| | Ighv1-12 | 0 | 0 | 8 | 3 | 5 | 42 | 33 | 19 | 31 | 12 | 7 | 2 | 0 | 3 | 4 | 20 | 15 | 6 | 14 | 7 | 10 | 4 | 10 | 8 | 3 | 23 | 29 | 25 | 26 | 3 |
| | Ighv1-11 | 14 | 16 | 11 | 14 | 3 | 85 | 43 | 67 | 65 | 21 | 21 | 14 | 21 | 19 | 4 | 39 | 35 | 63 | 46 | 15 | 35 | 53 | 47 | 45 | 9 | 41 | 31 | 66 | 46 | 18 |
| | Ighv1-9 | 4 | 2 | 17 | 8 | 8 | 15 | 12 | 17 | 15 | 3 | 0 | 6 | 4 | 3 | 3 | 10 | 5 | 3 | 6 | 4 | 5 | 6 | 8 | 6 | 2 | 6 | 11 | 18 | 12 | 6 |
| | Ighv15-2 | 7 | 0 | 2 | 3 | 4 | 17 | 12 | 16 | 15 | 3 | 4 | 5 | 3 | 4 | 1 | 1 | 10 | 6 | 6 | 5 | 18 | 14 | 7 | 13 | 6 | 6 | 5 | 18 | 10 | 7 |
| | Ighv1-7 | 1 | 0 | 0 | 0 | 1 | 3 | 9 | 12 | 8 | 5 | 0 | 0 | 7 | 2 | 4 | 1 | 0 | 11 | 4 | 6 | 1 | 3 | 6 | 3 | 3 | 11 | 1 | 0 | 4 | 6 |
| | Ighv10-3 | 0 | 2 | 5 | 2 | 3 | 13 | 7 | 16 | 12 | 5 | 0 | 2 | 2 | 1 | 1 | 1 | 6 | 19 | 9 | 9 | 2 | 5 | 10 | 6 | 4 | 13 | 8 | 7 | 9 | 3 |
| | Ighv1-5 | 0 | 0 | 2 | 1 | 1 | 7 | 0 | 0 | 2 | 4 | 0 | 0 | 1 | 0 | 1 | 4 | 0 | 3 | 2 | 2 | 0 | 0 | 0 | 0 | 0 | 0 | 0 | 6 | 2 | 3 |
| | Ighv1-4 | 0 | 0 | 1 | 0 | 1 | 0 | 0 | 0 | 0 | 0 | 0 | 1 | 0 | 0 | 1 | 1 | 0 | 0 | 0 | 1 | 0 | 4 | 1 | 2 | 2 | 0 | 1 | 1 | 1 | 1 |
| | Ighv10-1 | 3 | 9 | 1 | 4 | 4 | 13 | 30 | 18 | 20 | 9 | 1 | 2 | 5 | 3 | 2 | 7 | 6 | 12 | 8 | 3 | 1 | 15 | 26 | 14 | 13 | 7 | 13 | 31 | 17 | 12 |
| | Ighv1-2P | 1 | 2 | 5 | 3 | 2 | 2 | 2 | 1 | 2 | 1 | 0 | 0 | 5 | 2 | 3 | 1 | 1 | 6 | 3 | 3 | 1 | 0 | 1 | 1 | 1 | 1 | 3 | 0 | 1 | 2 |
| | Ighv8-2 ORF | 0 | 0 | 3 | 1 | 2 | 3 | 0 | 0 | 1 | 2 | 0 | 0 | 4 | 1 | 2 | 0 | 0 | 7 | 2 | 4 | 0 | 0 | 2 | 1 | 1 | 3 | 0 | 0 | 1 | 2 |
| | Ighv6-7 | 0 | 0 | 1 | 0 | 1 | 0 | 0 | 2 | 1 | 1 | 0 | 1 | 0 | 0 | 1 | 1 | 0 | 0 | 0 | 1 | 0 | 0 | 0 | 0 | 0 | 0 | 0 | 0 | 0 | 0 |
| | Ighv6-6 | 4 | 12 | 1 | 6 | 6 | 23 | 45 | 27 | 32 | 12 | 2 | 4 | 2 | 3 | 1 | 4 | 9 | 11 | 8 | 4 | 2 | 12 | 11 | 8 | 6 | 12 | 15 | 20 | 16 | 4 |
| | Ighv6-5 | 0 | 0 | 2 | 1 | 1 | 3 | 4 | 1 | 3 | 2 | 0 | 0 | 0 | 0 | 0 | 1 | 0 | 6 | 2 | 3 | 1 | 1 | 0 | 1 | 1 | 0 | 4 | 2 | 2 | 2 |
| | Ighv6-4 | 0 | 0 | 0 | 0 | 0 | 1 | 3 | 5 | 3 | 2 | 0 | 0 | 0 | 0 | 0 | 0 | 0 | 2 | 1 | 1 | 0 | 0 | 0 | 0 | 0 | 0 | 1 | 7 | 3 | 4 |
| | Ighv6-3 | 3 | 0 | 0 | 1 | 2 | 11 | 13 | 14 | 13 | 2 | 0 | 7 | 2 | 3 | 4 | 5 | 18 | 13 | 12 | 7 | 10 | 2 | 6 | 6 | 4 | 2 | 12 | 18 | 11 | 8 |
| | Ighv12-3 | 7 | 2 | 26 | 12 | 13 | 50 | 80 | 73 | 68 | 16 | 13 | 9 | 11 | 11 | 2 | 22 | 45 | 36 | 34 | 12 | 11 | 23 | 22 | 19 | 7 | 35 | 45 | 52 | 44 | 9 |
| | Ighv13-2 | 8 | 11 | 6 | 8 | 3 | 30 | 53 | 47 | 43 | 12 | 10 | 16 | 8 | 11 | 4 | 28 | 15 | 27 | 23 | 7 | 37 | 23 | 41 | 34 | 9 | 18 | 30 | 40 | 29 | 11 |
| | Ighv3-8 | 2 | 0 | 7 | 3 | 4 | 48 | 23 | 36 | 36 | 13 | 6 | 8 | 11 | 8 | 3 | 10 | 15 | 20 | 15 | 5 | 9 | 11 | 24 | 15 | 8 | 14 | 14 | 24 | 17 | 6 |
| | Ighv5-21P | 0 | 2 | 0 | 1 | 1 | 4 | 0 | 0 | 1 | 2 | 0 | 6 | 0 | 2 | 3 | 0 | 0 | 0 | 0 | 0 | 2 | 2 | 0 | 1 | 1 | 0 | 0 | 0 | 0 | 0 |
| | Ighv9-4 | 0 | 0 | 0 | 0 | 0 | 6 | 11 | 7 | 8 | 3 | 0 | 0 | 5 | 2 | 3 | 0 | 5 | 3 | 3 | 3 | 7 | 7 | 7 | 7 | 0 | 8 | 0 | 5 | 4 | 4 |
| | Ighv3-6 | 1 | 7 | 4 | 4 | 3 | 0 | 14 | 20 | 11 | 10 | 5 | 1 | 1 | 2 | 2 | 5 | 3 | 14 | 7 | 6 | 1 | 15 | 12 | 9 | 7 | 9 | 15 | 12 | 12 | 3 |
| | Ighv13-1 | 0 | 0 | 0 | 0 | 0 | 0 | 0 | 0 | 0 | 0 | 0 | 0 | 0 | 0 | 0 | 0 | 0 | 0 | 0 | 0 | 0 | 0 | 0 | 0 | 0 | 0 | 0 | 0 | 0 | 0 |
| | Ighv3-5 | 0 | 0 | 0 | 0 | 0 | 2 | 0 | 0 | 1 | 1 | 0 | 0 | 0 | 0 | 0 | 1 | 0 | 1 | 1 | 1 | 0 | 2 | 0 | 1 | 1 | 0 | 0 | 9 | 3 | 5 |
| | Ighv3-4 | 1 | 6 | 1 | 3 | 3 | 11 | 14 | 3 | 9 | 6 | 5 | 2 | 2 | 3 | 2 | 3 | 5 | 8 | 5 | 3 | 1 | 5 | 3 | 3 | 2 | 4 | 2 | 13 | 6 | 6 |
| | Ighv7-4 | 0 | 2 | 0 | 1 | 1 | 9 | 2 | 15 | 9 | 7 | 0 | 0 | 0 | 0 | 0 | 0 | 4 | 0 | 1 | 2 | 1 | 0 | 0 | 0 | 1 | 3 | 6 | 0 | 3 | 3 |
| | Ighv3-3 | 1 | 6 | 5 | 4 | 3 | 6 | 23 | 9 | 13 | 9 | 1 | 4 | 12 | 6 | 6 | 9 | 5 | 5 | 6 | 2 | 0 | 14 | 9 | 8 | 7 | 2 | 17 | 28 | 16 | 13 |
| | Ighv14-4 | 0 | 4 | 7 | 4 | 4 | 10 | 12 | 6 | 9 | 3 | 0 | 3 | 5 | 3 | 3 | 4 | 4 | 7 | 5 | 2 | 13 | 6 | 13 | 11 | 4 | 2 | 1 | 6 | 3 | 3 |
| Middle | Ighv7-3 | 3 | 0 | 4 | 2 | 2 | 7 | 8 | 21 | 12 | 8 | 0 | 2 | 1 | 1 | 1 | 2 | 5 | 14 | 7 | 6 | 3 | 10 | 46 | 20 | 23 | 9 | 12 | 11 | 11 | 2 |
| | Ighv9-3 | 0 | 2 | 11 | 4 | 6 | 5 | 11 | 9 | 8 | 3 | 2 | 3 | 9 | 5 | 4 | 0 | 9 | 0 | 3 | 5 | 6 | 9 | 4 | 6 | 3 | 8 | 11 | 8 | 9 | 2 |
| | Ighv12-2P | 0 | 0 | 1 | 0 | 1 | 0 | 0 | 0 | 0 | 0 | 0 | 0 | 0 | 0 | 0 | 0 | 0 | 0 | 0 | 0 | 0 | 0 | 0 | 0 | 0 | 0 | 0 | 0 | 0 | 0 |
| | Ighv9-2 | 0 | 0 | 3 | 1 | 2 | 2 | 5 | 3 | 3 | 2 | 0 | 0 | 0 | 0 | 0 | 0 | 10 | 0 | 3 | 6 | 5 | 8 | 1 | 5 | 4 | 0 | 2 | 0 | 1 | 1 |
| | Ighv12-1 | 2 | 10 | 0 | 4 | 5 | 0 | 9 | 2 | 4 | 5 | 2 | 0 | 3 | 2 | 2 | 0 | 1 | 2 | 1 | 1 | 1 | 0 | 0 | 0 | 1 | 2 | 5 | 0 | 2 | 3 |
| | Ighv9-1 | 1 | 0 | 0 | 0 | 1 | 0 | 1 | 2 | 1 | 1 | 1 | 0 | 1 | 1 | 1 | 0 | 0 | 2 | 1 | 1 | 2 | 13 | 2 | 6 | 6 | 2 | 2 | 0 | 1 | 1 |
| | Ighv16-1 | 0 | 0 | 0 | 0 | 0 | 0 | 0 | 0 | 0 | 0 | 0 | 0 | 0 | 0 | 0 | 0 | 0 | 0 | 0 | 0 | 0 | 0 | 0 | 0 | 0 | 0 | 0 | 1 | 0 | 1 |
| | Ighv14-3 | 0 | 0 | 0 | 0 | 0 | 0 | 1 | 13 | 5 | 7 | 3 | 1 | 2 | 2 | 1 | 1 | 6 | 9 | 5 | 4 | 10 | 0 | 3 | 4 | 5 | 11 | 0 | 0 | 4 | 6 |
| | Ighv11-2 | 3 | 1 | 4 | 3 | 2 | 21 | 15 | 32 | 23 | 9 | 15 | 3 | 10 | 9 | 6 | 10 | 3 | 4 | 6 | 4 | 27 | 3 | 33 | 21 | 16 | 8 | 15 | 23 | 15 | 8 |
| | Ighv3-2 | 10 | 1 | 0 | 4 | 6 | 20 | 4 | 9 | 11 | 8 | 3 | 4 | 12 | 6 | 5 | 2 | 9 | 5 | 5 | 4 | 11 | 14 | 16 | 14 | 3 | 4 | 4 | 24 | 11 | 12 |
| | Ighv4-2 | 0 | 0 | 3 | 1 | 2 | 6 | 4 | 3 | 4 | 2 | 2 | 5 | 6 | 4 | 2 | 2 | 4 | 2 | 3 | 1 | 22 | 5 | 9 | 12 | 9 | 3 | 2 | 12 | 6 | 6 |
| | Ighv14-2 | 8 | 1 | 4 | 4 | 4 | 12 | 1 | 9 | 7 | 6 | 17 | 0 | 5 | 7 | 9 | 8 | 5 | 5 | 6 | 2 | 32 | 5 | 24 | 20 | 14 | 1 | 6 | 6 | 4 | 3 |
| | Ighv11-1 | 10 | 8 | 2 | 7 | 4 | 7 | 7 | 11 | 8 | 2 | 12 | 14 | 12 | 13 | 1 | 1 | 3 | 1 | 2 | 1 | 27 | 25 | 45 | 32 | 11 | 12 | 7 | 8 | 9 | 3 |
| | Ighv3-1 | 0 | 0 | 5 | 2 | 3 | 5 | 6 | 7 | 6 | 1 | 1 | 4 | 3 | 3 | 2 | 1 | 0 | 0 | 0 | 1 | 5 | 3 | 6 | 5 | 2 | 0 | 0 | 9 | 3 | 5 |
| | Ighv4-1 | 0 | 3 | 2 | 2 | 2 | 5 | 5 | 0 | 3 | 3 | 4 | 8 | 5 | 6 | 2 | 5 | 2 | 0 | 2 | 3 | 10 | 12 | 20 | 14 | 5 | 4 | 3 | 2 | 3 | 1 |
| | Ighv14-1 | 0 | 0 | 1 | 0 | 1 | 0 | 0 | 3 | 1 | 2 | 0 | 3 | 0 | 1 | 2 | 0 | 0 | 0 | 0 | 0 | 0 | 0 | 3 | 1 | 2 | 0 | 4 | 0 | 1 | 2 |
| | Ighv7-2 | 0 | 0 | 0 | 0 | 0 | 0 | 0 | 0 | 0 | 0 | 0 | 0 | 0 | 0 | 0 | 0 | 0 | 0 | 0 | 0 | 0 | 0 | 0 | 0 | 0 | 0 | 0 | 0 | 0 | 0 |
| | Ighv7-1 | 11 | 17 | 12 | 13 | 3 | 14 | 30 | 7 | 17 | 12 | 21 | 13 | 24 | 19 | 6 | 0 | 10 | 20 | 10 | 10 | 40 | 40 | 58 | 46 | 10 | 20 | 11 | 14 | 15 | 5 |
| | Ighv2-9 | 10 | 8 | 15 | 11 | 4 | 13 | 3 | 5 | 7 | 5 | 38 | 12 | 11 | 20 | 15 | 8 | 11 | 22 | 14 | 7 | 25 | 32 | 48 | 35 | 12 | 22 | 13 | 17 | 17 | 5 |

| | | | | | | | | | | | | | | | | | | | | | | | | | | | | | | | |
|---|---|---|---|---|---|---|---|---|---|---|---|---|---|---|---|---|---|---|---|---|---|---|---|---|---|---|---|---|---|---|---|
| | Ighv5-17 | 32 | 10 | 12 | 18 | 12 | 5 | 17 | 8 | 10 | 6 | 23 | 22 | 13 | 19 | 6 | 5 | 4 | 12 | 7 | 4 | 30 | 48 | 66 | 48 | 18 | 3 | 24 | 10 | 12 | 11 |
| | Ighv5-16 | 3 | 3 | 6 | 4 | 2 | 8 | 6 | 8 | 7 | 1 | 3 | 4 | 5 | 4 | 1 | 3 | 7 | 5 | 5 | 2 | 2 | 11 | 12 | 8 | 6 | 2 | 6 | 16 | 8 | 7 |
| | Ighv5-15 | 13 | 9 | 11 | 11 | 2 | 4 | 7 | 5 | 5 | 2 | 21 | 9 | 14 | 15 | 6 | 0 | 8 | 8 | 5 | 5 | 5 | 12 | 27 | 15 | 11 | 2 | 14 | 8 | 8 | 6 |
| | Ighv2-7 | 0 | 0 | 1 | 0 | 1 | 1 | 0 | 0 | 0 | 1 | 0 | 0 | 0 | 0 | 0 | 0 | 0 | 0 | 0 | 0 | 0 | 0 | 0 | 0 | 0 | 0 | 0 | 3 | 1 | 2 |
| | Ighv2-6-8 | 7 | 7 | 11 | 8 | 2 | 6 | 9 | 0 | 5 | 5 | 11 | 8 | 12 | 10 | 2 | 6 | 5 | 6 | 6 | 1 | 0 | 5 | 8 | 4 | 4 | 0 | 2 | 16 | 6 | 9 |
| | Ighv2-9-1 | 2 | 9 | 7 | 6 | 4 | 3 | 4 | 0 | 2 | 2 | 13 | 35 | 18 | 22 | 12 | 7 | 3 | 8 | 6 | 3 | 15 | 22 | 26 | 21 | 6 | 4 | 3 | 3 | 3 | 1 |
| | Ighv5-12-4 | 0 | 0 | 1 | 0 | 1 | 0 | 0 | 0 | 0 | 0 | 0 | 0 | 0 | 0 | 0 | 0 | 0 | 0 | 0 | 0 | 0 | 0 | 1 | 0 | 1 | 0 | 0 | 0 | 0 | 0 |
| | Ighv5-9-1 | 4 | 14 | 14 | 11 | 6 | 3 | 5 | 11 | 6 | 4 | 21 | 27 | 32 | 27 | 6 | 5 | 0 | 3 | 3 | 3 | 29 | 58 | 83 | 57 | 27 | 1 | 2 | 1 | 1 | 1 |
| 7183/Q52 | Ighv2-6 | 29 | 42 | 18 | 30 | 12 | 2 | 3 | 6 | 4 | 2 | 15 | 35 | 14 | 21 | 12 | 11 | 1 | 1 | 4 | 6 | 36 | 44 | 62 | 47 | 13 | 1 | 1 | 7 | 3 | 3 |
| | Ighv5-12 | 49 | 54 | 46 | 50 | 4 | 11 | 14 | 12 | 12 | 2 | 65 | 92 | 95 | 84 | 17 | 9 | 4 | 3 | 5 | 3 | 128 | 169 | 188 | 162 | 31 | 19 | 20 | 10 | 16 | 6 |
| | Ighv2-5 | 7 | 25 | 18 | 17 | 9 | 4 | 0 | 2 | 2 | 2 | 12 | 19 | 16 | 16 | 4 | 8 | 1 | 2 | 4 | 4 | 26 | 44 | 23 | 31 | 11 | 2 | 12 | 15 | 10 | 7 |
| | Ighv5-9 | 3 | 1 | 8 | 4 | 4 | 4 | 16 | 0 | 7 | 8 | 3 | 0 | 4 | 2 | 2 | 0 | 2 | 5 | 2 | 3 | 0 | 6 | 5 | 4 | 3 | 0 | 0 | 6 | 2 | 3 |
| | Ighv2-4 | 0 | 2 | 2 | 1 | 1 | 0 | 0 | 0 | 0 | 0 | 8 | 2 | 2 | 4 | 3 | 5 | 0 | 0 | 2 | 3 | 2 | 5 | 3 | 3 | 2 | 2 | 1 | 3 | 2 | 1 |
| | **Ighv5-6** | 78 | 62 | 65 | 68 | 9 | 3 | 15 | 12 | 10 | 6 | 86 | 83 | 81 | 83 | 3 | 3 | 4 | 10 | 6 | 4 | 122 | 215 | 262 | 200 | 71 | 26 | 13 | 6 | 15 | 10 |
| | **Ighv2-3** | 96 | 135 | 128 | 120 | 21 | 14 | 22 | 1 | 12 | 11 | 87 | 92 | 73 | 84 | 10 | 13 | 8 | 14 | 12 | 3 | 55 | 161 | 141 | 119 | 56 | 23 | 4 | 11 | 13 | 10 |
| | **Ighv5-4** | 128 | 164 | 149 | 147 | 18 | 9 | 16 | 17 | 14 | 4 | 147 | 172 | 135 | 151 | 19 | 15 | 14 | 15 | 15 | 1 | 185 | 200 | 243 | 209 | 30 | 66 | 17 | 16 | 33 | 29 |
| | **Ighv2-2** | 318 | 442 | 393 | 384 | 62 | 27 | 59 | 36 | 41 | 17 | 296 | 315 | 222 | 278 | 49 | 19 | 13 | 29 | 20 | 8 | 285 | 361 | 419 | 355 | 67 | 77 | 8 | 85 | 57 | 42 |
| | **Ighv5-2** | 866 | 960 | 992 | 939 | 65 | 44 | 74 | 87 | 68 | 22 | 466 | 526 | 477 | 490 | 32 | 40 | 36 | 43 | 40 | 4 | 593 | 746 | 822 | 720 | 117 | 101 | 47 | 81 | 76 | 27 |
| | **Ighv5-1P** | 35 | 38 | 62 | 45 | 15 | 4 | 16 | 8 | 9 | 6 | 33 | 20 | 16 | 23 | 9 | 0 | 3 | 5 | 3 | 3 | 27 | 15 | 20 | 21 | 6 | 2 | 5 | 2 | 3 | 2 |
| | D usage in DJH joins | | | | | | | | | | | | | | | | | | | | | | | | | | | | | | |
| | IGHD1-1 (DFL16.1) | 75965 | 80946 | 78077 | 78329 | 2500 | 5292 | 5843 | 5868 | 5668 | 326 | 66716 | 61828 | 58299 | 62281 | 4227 | 4715 | 5485 | 4728 | 4976 | 441 | 44816 | 64508 | 81613 | 63646 | 18414 | 6502 | 6205 | 8223 | 6977 | 1090 |
| | IGHD6-1 | 343 | 277 | 312 | 311 | 33 | 59 | 46 | 50 | 52 | 7 | 227 | 243 | 199 | 223 | 22 | 30 | 39 | 56 | 42 | 13 | 304 | 371 | 337 | 337 | 34 | 108 | 36 | 62 | 69 | 36 |
| | IGHD2-3 | 16495 | 17329 | 17930 | 17251 | 721 | 4502 | 4742 | 5180 | 4808 | 344 | 13499 | 19300 | 14294 | 15698 | 3145 | 4428 | 3515 | 3647 | 3863 | 493 | 24403 | 21368 | 23121 | 22964 | 1524 | 12387 | 5664 | 6816 | 8289 | 3595 |
| | IGHD6-2 | 65 | 56 | 71 | 64 | 8 | 20 | 25 | 21 | 22 | 3 | 41 | 116 | 43 | 67 | 43 | 15 | 1 | 9 | 8 | 7 | 144 | 98 | 106 | 116 | 25 | 52 | 24 | 29 | 35 | 15 |
| | IGHD2-4 | 3320 | 3134 | 3336 | 3263 | 112 | 1109 | 1146 | 1273 | 1176 | 86 | 2618 | 2580 | 2379 | 2526 | 128 | 867 | 997 | 824 | 896 | 90 | 5340 | 2994 | 4256 | 4197 | 1174 | 2821 | 1331 | 1696 | 1949 | 777 |
| | IGHD2-5 | 616 | 506 | 499 | 540 | 66 | 245 | 264 | 244 | 251 | 11 | 330 | 358 | 357 | 348 | 16 | 123 | 156 | 136 | 138 | 17 | 818 | 403 | 559 | 593 | 210 | 528 | 187 | 235 | 317 | 185 |
| | IGHD2-6 | 450 | 390 | 427 | 422 | 30 | 212 | 204 | 173 | 196 | 21 | 312 | 232 | 286 | 277 | 41 | 125 | 106 | 134 | 122 | 14 | 537 | 307 | 378 | 407 | 118 | 351 | 133 | 221 | 235 | 110 |
| | IGHD2-7 | 348 | 356 | 439 | 381 | 50 | 190 | 148 | 207 | 182 | 30 | 287 | 225 | 182 | 231 | 53 | 110 | 169 | 115 | 131 | 33 | 495 | 181 | 365 | 347 | 158 | 422 | 71 | 144 | 212 | 185 |
| | IGHD2-8 | 3678 | 4022 | 3850 | 3850 | 172 | 1802 | 1984 | 2015 | 1934 | 115 | 2914 | 2581 | 2522 | 2672 | 211 | 1315 | 1501 | 1221 | 1346 | 142 | 5040 | 2623 | 3792 | 3818 | 1209 | 3615 | 1360 | 2096 | 2357 | 1150 |
| | IGHD3-2 | 357 | 488 | 493 | 446 | 77 | 189 | 357 | 248 | 265 | 85 | 338 | 230 | 239 | 269 | 60 | 140 | 188 | 123 | 150 | 34 | 599 | 416 | 507 | 507 | 92 | 368 | 279 | 317 | 321 | 45 |
| | IGHD4-1 (DQ52) | 1863 | 2177 | 2192 | 2077 | 186 | 1922 | 2067 | 2049 | 2013 | 79 | 1455 | 1411 | 1254 | 1373 | 106 | 999 | 1091 | 973 | 1021 | 62 | 1643 | 1512 | 1788 | 1648 | 138 | 1349 | 1159 | 1571 | 1360 | 206 |
| | D usage in VHDJH joins | | | | | | | | | | | | | | | | | | | | | | | | | | | | | | |
| | IGHD1-1 (DFL16.1) | 743 | 815 | 837 | 798 | 49 | 472 | 516 | 612 | 533 | 71 | 774 | 772 | 624 | 723 | 86 | 256 | 372 | 315 | 314 | 58 | 780 | 1109 | 1496 | 1128 | 358 | 244 | 459 | 577 | 427 | 169 |
| | IGHD6-1 | 0 | 0 | 6 | 2 | 3 | 1 | 0 | 5 | 2 | 3 | 1 | 3 | 0 | 1 | 2 | 0 | 2 | 2 | 1 | 1 | 1 | 5 | 2 | 3 | 2 | 2 | 2 | 1 | 2 | 1 |
| | IGHD2-3 | 485 | 575 | 573 | 544 | 52 | 630 | 641 | 626 | 633 | 8 | 387 | 515 | 570 | 491 | 94 | 347 | 291 | 358 | 332 | 36 | 734 | 875 | 963 | 857 | 115 | 528 | 467 | 589 | 528 | 61 |
| | IGHD6-2 | 0 | 6 | 5 | 4 | 3 | 4 | 0 | 2 | 2 | 2 | 0 | 0 | 0 | 0 | 0 | 0 | 0 | 0 | 0 | 0 | 7 | 1 | 2 | 3 | 4 | 2 | 2 | 5 | 3 | 2 |
| | IGHD2-4 | 141 | 163 | 164 | 156 | 13 | 116 | 174 | 100 | 130 | 39 | 104 | 108 | 99 | 104 | 5 | 49 | 47 | 81 | 59 | 19 | 177 | 169 | 233 | 193 | 35 | 127 | 62 | 111 | 100 | 34 |
| | IGHD2-5 | 49 | 60 | 56 | 55 | 5 | 35 | 47 | 30 | 37 | 9 | 26 | 27 | 33 | 29 | 4 | 17 | 16 | 18 | 17 | 1 | 37 | 47 | 46 | 43 | 5 | 29 | 15 | 38 | 27 | 12 |
| | IGHD2-6 | 49 | 60 | 56 | 55 | 5 | 35 | 47 | 30 | 37 | 9 | 26 | 27 | 33 | 29 | 4 | 17 | 16 | 18 | 17 | 1 | 37 | 47 | 46 | 43 | 5 | 29 | 15 | 38 | 27 | 12 |
| | IGHD2-7 | 33 | 22 | 58 | 38 | 18 | 8 | 25 | 21 | 18 | 9 | 13 | 19 | 22 | 18 | 5 | 1 | 8 | 10 | 7 | 5 | 44 | 22 | 2 | 23 | 21 | 18 | 5 | 5 | 9 | 8 |
| | IGHD2-8 | 160 | 186 | 184 | 177 | 15 | 151 | 182 | 143 | 158 | 21 | 76 | 67 | 65 | 69 | 5 | 34 | 46 | 93 | 58 | 31 | 94 | 103 | 138 | 111 | 24 | 71 | 80 | 118 | 90 | 25 |
| | IGHD3-2 | 19 | 32 | 39 | 30 | 10 | 12 | 17 | 30 | 20 | 10 | 11 | 12 | 9 | 10 | 2 | 7 | 5 | 14 | 9 | 5 | 16 | 41 | 21 | 26 | 13 | 9 | 14 | 13 | 12 | 2 |
| | IGHD4-1 (DQ52) | 45 | 60 | 44 | 50 | 9 | 22 | 28 | 58 | 36 | 19 | 24 | 23 | 9 | 18 | 8 | 9 | 17 | 33 | 20 | 12 | 11 | 35 | 54 | 33 | 22 | 13 | 8 | 32 | 18 | 12 |
| | D usage in DJH+VHDJH joins | | | | | | | | | | | | | | | | | | | | | | | | | | | | | | |
| | IGHD1-1 (DFL16.1) | 76708 | 81761 | 78914 | 79128 | 2534 | 5764 | 6359 | 6480 | 6201 | 383 | 67490 | 62600 | 58923 | 63004 | 4297 | 4971 | 5857 | 5043 | 5290 | 492 | 45596 | 65617 | 83109 | 64774 | 18771 | 6746 | 6664 | 8800 | 7403 | 1210 |
| | IGHD6-1 | 343 | 277 | 318 | 313 | 33 | 60 | 46 | 55 | 54 | 7 | 228 | 246 | 199 | 224 | 24 | 30 | 41 | 58 | 43 | 14 | 305 | 376 | 339 | 340 | 35 | 110 | 38 | 63 | 70 | 37 |
| | IGHD2-3 | 16980 | 17904 | 18503 | 17796 | 768 | 5132 | 5383 | 5806 | 5441 | 340 | 13886 | 19815 | 14864 | 16188 | 3179 | 4775 | 3806 | 4005 | 4195 | 512 | 25137 | 22243 | 24084 | 23821 | 1465 | 12915 | 6131 | 7405 | 8817 | 3606 |
| | IGHD6-2 | 65 | 62 | 76 | 68 | 7 | 24 | 25 | 23 | 24 | 1 | 41 | 116 | 43 | 67 | 43 | 15 | 1 | 9 | 8 | 7 | 151 | 99 | 108 | 119 | 28 | 54 | 26 | 34 | 38 | 15 |
| | IGHD2-4 | 3461 | 3297 | 3500 | 3419 | 108 | 1225 | 1320 | 1373 | 1306 | 75 | 2722 | 2688 | 2478 | 2629 | 133 | 916 | 1044 | 905 | 955 | 77 | 5517 | 3163 | 4489 | 4390 | 1180 | 2948 | 1393 | 1807 | 2049 | 806 |
| | IGHD2-5 | 665 | 566 | 555 | 595 | 61 | 280 | 311 | 274 | 288 | 20 | 356 | 385 | 390 | 377 | 19 | 140 | 172 | 154 | 155 | 16 | 855 | 450 | 605 | 637 | 204 | 557 | 202 | 273 | 344 | 188 |
| | IGHD2-6 | 499 | 450 | 483 | 477 | 25 | 247 | 251 | 203 | 234 | 27 | 338 | 259 | 319 | 306 | 41 | 142 | 122 | 152 | 139 | 15 | 574 | 354 | 424 | 451 | 112 | 380 | 148 | 259 | 262 | 116 |
| | IGHD2-7 | 381 | 378 | 497 | 419 | 67 | 198 | 173 | 228 | 200 | 27 | 300 | 244 | 204 | 249 | 48 | 111 | 177 | 125 | 138 | 35 | 539 | 203 | 367 | 370 | 168 | 440 | 76 | 149 | 222 | 193 |
| | IGHD2-8 | 3838 | 4208 | 4034 | 4027 | 185 | 1953 | 2166 | 2158 | 2092 | 121 | 2990 | 2648 | 2587 | 2742 | 217 | 1349 | 1547 | 1314 | 1403 | 126 | 5134 | 2726 | 3930 | 3930 | 1204 | 3686 | 1440 | 2214 | 2447 | 1141 |
| | IGHD3-2 | 376 | 520 | 532 | 476 | 87 | 201 | 374 | 278 | 284 | 87 | 349 | 242 | 248 | 279 | 60 | 147 | 193 | 137 | 159 | 30 | 615 | 457 | 528 | 533 | 79 | 377 | 293 | 330 | 333 | 42 |
| | IGHD4-1 (DQ52) | 1908 | 2237 | 2236 | 2127 | 190 | 1944 | 2095 | 2107 | 2049 | 91 | 1479 | 1434 | 1263 | 1392 | 114 | 1008 | 1108 | 1006 | 1041 | 58 | 1654 | 1547 | 1842 | 1681 | 150 | 1362 | 1167 | 1603 | 1377 | 218 |

Notes:
1. Data were nomalized to 1,964,102 total reads.
2. Proximal V and D segments are denoted in bold.

**Supplementary Table 4 | Absolute utilization of Igh D to JHs1-4 in WAPL-degron v-Abl pro-B cells. Data was extracted from previously published data (22: GSM4593296-GSM4593303)**

| D usage in DJH joins | WT untreated JH1 bait primer (n=3) | | | | | WT untreated JH2 bait primer (n=3) | | | | | WT untreated JH3 bait primer (n=3) | | | | | WT untreated JH4 bait primer (n=3) | | | | |
|---|---|---|---|---|---|---|---|---|---|---|---|---|---|---|---|---|---|---|---|---|
| | #1 | #2 | #3 | Average | s.d. | #1 | #2 | #3 | Average | s.d. | #1 | #2 | #3 | Average | s.d. | #1 | #2 | #3 | Average | s.d. |
| **IGHD1-1 (DFL16.1)** | 108484 | 113218 | 96778 | 106160 | 8463 | 173771 | 182460 | 164495 | 173575 | 8984 | 46828 | 53634 | 47131 | 49198 | 3845 | 23338 | 23810 | 20867 | 22672 | 1581 |
| **IGHD6-1** | 104 | 89 | 85 | 93 | 10 | 736 | 555 | 553 | 615 | 105 | 195 | 163 | 137 | 165 | 29 | 63 | 45 | 27 | 45 | 18 |
| **IGHD2-3** | 6500 | 6020 | 4998 | 5839 | 767 | 13214 | 13239 | 10743 | 12399 | 1434 | 4732 | 4528 | 3744 | 4335 | 522 | 2510 | 2244 | 1834 | 2196 | 341 |
| **IGHD6-2** | 12 | 22 | 15 | 16 | 5 | 73 | 72 | 70 | 72 | 2 | 16 | 27 | 21 | 21 | 6 | 7 | 3 | 11 | 7 | 4 |
| **IGHD2-4** | 1749 | 1662 | 1562 | 1658 | 94 | 4546 | 4121 | 3897 | 4188 | 330 | 1456 | 1456 | 1288 | 1400 | 97 | 669 | 582 | 483 | 578 | 93 |
| **IGHD2-5** | 522 | 445 | 427 | 465 | 50 | 1140 | 1010 | 1066 | 1072 | 65 | 292 | 278 | 239 | 270 | 27 | 173 | 106 | 119 | 133 | 36 |
| **IGHD2-6** | 425 | 352 | 346 | 374 | 44 | 839 | 766 | 791 | 799 | 37 | 205 | 182 | 180 | 189 | 14 | 114 | 100 | 100 | 105 | 8 |
| **IGHD2-7** | 181 | 216 | 195 | 197 | 18 | 595 | 624 | 506 | 575 | 61 | 158 | 159 | 122 | 146 | 21 | 60 | 71 | 40 | 57 | 16 |
| **IGHD2-8** | 2743 | 2782 | 2107 | 2544 | 379 | 5239 | 4995 | 4562 | 4932 | 343 | 1386 | 1399 | 1236 | 1340 | 91 | 816 | 757 | 534 | 702 | 149 |
| **IGHD3-2** | 135 | 131 | 114 | 127 | 11 | 1702 | 1603 | 1335 | 1547 | 190 | 426 | 422 | 316 | 388 | 62 | 124 | 101 | 104 | 110 | 13 |
| **IGHD4-1 (DQ52)** | 12801 | 12213 | 9782 | 11599 | 1601 | 48201 | 44001 | 38932 | 43711 | 4641 | 7441 | 7342 | 5820 | 6868 | 909 | 1342 | 1135 | 877 | 1118 | 233 |

| D usage in DJH joins | WT+Dox/IAA JH1 bait primer (n=3) | | | | | WT+Dox/IAA JH2 bait primer (n=3) | | | | | WT+Dox/IAA JH3 bait primer (n=3) | | | | | WT+Dox/IAA JH4 bait primer (n=3) | | | | |
|---|---|---|---|---|---|---|---|---|---|---|---|---|---|---|---|---|---|---|---|---|
| | #1 | #2 | #3 | Average | s.d. | #1 | #2 | #3 | Average | s.d. | #1 | #2 | #3 | Average | s.d. | #1 | #2 | #3 | Average | s.d. |
| **IGHD1-1 (DFL16.1)** | 5362 | 5383 | 5609 | 5451 | 137 | 8355 | 7916 | 8341 | 8204 | 250 | 1857 | 1814 | 1796 | 1822 | 31 | 789 | 816 | 899 | 835 | 57 |
| **IGHD6-1** | 11 | 7 | 13 | 10 | 3 | 79 | 66 | 58 | 68 | 11 | 9 | 9 | 10 | 9 | 1 | 4 | 1 | 6 | 4 | 3 |
| **IGHD2-3** | 1051 | 1042 | 1215 | 1103 | 97 | 2053 | 1973 | 2036 | 2021 | 42 | 595 | 592 | 581 | 589 | 7 | 336 | 366 | 387 | 363 | 26 |
| **IGHD6-2** | 1 | 0 | 2 | 1 | 1 | 15 | 11 | 23 | 16 | 6 | 3 | 1 | 1 | 2 | 1 | 2 | 1 | 0 | 1 | 1 |
| **IGHD2-4** | 414 | 411 | 402 | 409 | 6 | 852 | 782 | 854 | 829 | 41 | 228 | 225 | 261 | 238 | 20 | 132 | 129 | 123 | 128 | 5 |
| **IGHD2-5** | 122 | 103 | 139 | 121 | 18 | 225 | 202 | 215 | 214 | 12 | 68 | 46 | 50 | 55 | 12 | 27 | 21 | 36 | 28 | 8 |
| **IGHD2-6** | 86 | 98 | 123 | 102 | 19 | 165 | 161 | 182 | 169 | 11 | 55 | 28 | 34 | 39 | 14 | 45 | 27 | 36 | 36 | 9 |
| **IGHD2-7** | 80 | 87 | 76 | 81 | 6 | 144 | 154 | 155 | 151 | 6 | 34 | 21 | 29 | 28 | 7 | 24 | 22 | 16 | 21 | 4 |
| **IGHD2-8** | 784 | 716 | 904 | 801 | 95 | 1332 | 1263 | 1268 | 1288 | 38 | 342 | 321 | 320 | 328 | 12 | 165 | 185 | 229 | 193 | 33 |
| **IGHD3-2** | 23 | 29 | 20 | 24 | 5 | 324 | 299 | 291 | 305 | 17 | 75 | 79 | 80 | 78 | 3 | 36 | 34 | 34 | 35 | 1 |
| **IGHD4-1 (DQ52)** | 7214 | 7232 | 7914 | 7453 | 399 | 32811 | 31355 | 33050 | 32405 | 917 | 4880 | 4831 | 5007 | 4906 | 91 | 336 | 348 | 390 | 358 | 28 |

Notes:
1. Data using different bait primer were nomalized to the indicated total reads: 672.689 reads for JH1 primer: 946.994 reads for JH2 primer: 410.987 reads for JH3 primer: 614.285 reads for JH4 primer.
2. Data was extracted from previously published data (22: GSM4593296-GSM4593303)

**Supplementary Table 5 | Oligos used in this study.**

| | |
|---|---|
| **Genotyping primers** | |
| CBE1 WT Forward | AGTCTCTGCTTCCCCCTTGT |
| CBE1 WT Reverse | AGCACATGACCCTTCTCACC |
| CBE1 scramble Forward | AGCACATGACCCTTCTCACC |
| CBE1 scramble Reverse | CTCTCCTGCTAGCCTTCTCG |
| CBE1 inversion Forward | CAGTAATGCTCATGGCCACAAG |
| CBE1 inversion Reverse | AGCACATGACCCTTCTCACC |
| CBE2 WT Forward | TCTCCACAAGAGGGCAGAAT |
| CBE2 WT Reverse | CTTATGGAATGGGTGGGTTG |
| CBE2 scramble Forward | CTTATGGAATGGGTGGGTTG |
| CBE2 scramble Reverse | GGGAGATCTGTATCTCTTCCACTAGTAA |
| CBE2 inversion Forward | ATCTCCACAAGAGGGCAGAAG |
| CBE2 inversion Reverse | TGTCACCCTTGAGGCCATTC |
| **sgRNA used for 3'CBE deletion and inversion** | |
| 3'CBE sgRNA-1 | AGCCAGTGTTATAGTATCTC |
| 3'CBE sgRNA-2 | AGTCCCAACTTTTGCTTTAG |
| 3'CBE sgRNA-3 | GTCCCAACTTTTGCTTTAGC |
| **Oligos for generating HTGTS-V(D)J-seq and 3C-HTGTS libraries** | |
| **HTGTS-VDJ-seq bait primers** | |
| JH4 Coding end bio primer (129SV background) | /5BiosG/CCCTCAGGGACAAATATCCA |
| JH4 Coding end nested primer (129SV background) | CTGCAATGCTCAGAAAACTCC |
| JH4 Coding end bio primer (C57B6/L background) | /5BiosG/CTGCAATGCTCAGAAAACTCC |
| JH4 Coding end nested primer (C57B6/L background) | CTTACCTGAGGAGACGGTGAC |
| **3C-HTGTS bait primers** | |
| iEμ Bio primer | /5BiosG/GGTTATGTAAGAAATTGAAGGACTTTAGTG |
| iEμ Nested primer | CTCTATTATTCTTCCCTCTGATTATTGG |
| VH81X Bio primer | /5BiosG/AAATAGAAGATGAAATGGAAGATTTGAAGG |
| VH81X Nested primer | TGAGAAACACCAATATTGTCAACTAACC |
| CBE1 Bio primer | /5BiosG/CATGAGGCTCCTGGAGGTGAC |
| CBE1 Nested primer | GGTGACCAGGAAAGCTGCTTAG |
